# Dynamics of ML-based Morphological Features Indicate a Shear Stress-Dependent Bifurcation of hiPSC-Derived Endothelial Cell States

**DOI:** 10.64898/2026.07.07.736803

**Authors:** Erin Angelini, Chantelle L. Leveille, Serge E. Parent, Rebecca J. Zaunbrecher, Tiffany Barszczewski, Julie C. Dixon, Fatwir S. Mohammed, Benjamin Morris, Jessica S. Yu, Joy Arakaki, Renske J. Dupar, Jacqueline H. Edmonds, Erik A. Ehlers, Clare R. Gamlin, Maxwell J. Hedayati, Caroline Hookway, Jacob McCarley, Saurabh Mogre, Amber Phan, Brock Roberts, Emmanuel E. Sanchez, John Paul Thottam, Chamari S. Wijesooriya, Jie Yao, Matthew L. Kutys, Ehssan Nazockdast, Jin Wang, Julie A. Theriot, Gokhan Dalgin, Susanne M. Rafelski, Matheus P. Viana

## Abstract

Cell states are increasingly conceptualized as attractors of high-dimensional dynamical systems, yet quantitative approaches for integrating phenotypic information into this framework remain limited. Here, we take an image-based approach that combines unsupervised machine learning (ML) with timelapse imaging to extract and characterize the temporal dynamics of morphological features. Using a cell line with endogenously tagged VE-cadherin, we acquired brightfield and fluorescence timelapse images of human induced pluripotent stem cell-derived endothelial cell (hiPSC-EC) monolayers, which adopt distinct phenotypes at two different magnitudes of shear stress in terms of their morphology, behavior, and VE-cadherin organization. To quantify these phenotypic cell states without segmentation, we trained a diffusion autoencoder to predict VE-cadherin signal from brightfield images. We identified interpretable ML-based features representing cell orientation, elongation, and local density. Treating these variables as dimensions of a morphological state space, we estimated a data-driven vector field and found that the two observed phenotypic cell states correspond to stable fixed points of the inferred dynamical system. Mapping measured cell migration coherence onto this space further distinguished the states. Imaging cells across intermediate shear stresses revealed a regime of bistability in which both states coexist, indicating that the shear-stress-dependent transition between endothelial cell states occurs as a bifurcation of the inferred dynamical system. Finally, we applied this method to study an N-terminal truncation of VE-cadherin, finding that mutated cells preserve alignment and coherent migration, but exhibit altered morphology and increased migration speed. This work demonstrates the applicability of a dynamical systems approach to quantitatively characterize morphological aspects of cell state from interpretable ML-based features.

## Introduction

A fundamental understanding of cell and developmental biology requires rigorously defining and characterizing diverse cell states and their functions. However, how different two cells are from each other can vary widely. At one extreme, terminally differentiated cell types (such as neurons, cardiomyocytes, etc.), can be unambiguously defined functionally, visually and molecularly as distinct cell states. Cells can also be assigned to distinct states at identifiable intermediate steps along the continuous developmental pathways toward these terminal cell fates.^1,2^ Independently of differentiation trajectory, cells are also described as being in distinct, more or less reversible states such as proliferative, quiescent or apoptotic.^3^ Assigning cells to more intermediate, less easily distinguishable cell states is more difficult, given the continuous differences between states and the heterogeneity among cells within one state.^4^ At the other extreme, cell state can be defined as all measurable characteristics of a cell at any moment in time.^5^ In this case it is difficult to set a criterion, such as an acceptable range of measurement values, to categorize a group of cells into a state.

A well-established mathematical framework reconciles these disparate views by treating the discrete and continuous extremes as two features of a single description.^6–8^ A cell is represented as a collection of cell state variables whose possible values define a theoretical continuous cell state space. The framework’s core assumption is that an equation of motion describes how a cell traverses this state space, as a dynamical system in which a theoretical vector field maps values of cell state variables to velocities.^6–8^ Discrete cell states then emerge as points acting as sinks, or attractors, of this theoretical vector field.^6–8^

This framework was developed for gene expression-related cell state variables, given well-established mechanistic, control theory-based dynamical systems models for gene regulatory networks.^9–12^ The focus on dynamical systems in this framework was also motivated by the goal of quantifying a cell state “landscape” (analogous to Waddington’s epigenetic landscape and the energy landscapes of protein folding) that accommodates the energy-driven, active nature of living systems.^7,8,12–16^ In particular, only by considering cell state dynamics and the theoretical vector field can one reconcile a cell state landscape with the fact that biological systems are far from thermodynamic equilibrium .^6–8,17,18^ This combination of landscape and cell state dynamics has been used to visualize and characterize cell state transitions, from cell fate decisions in development^7,14,15,19–22^ to phenotypic plasticity in cancer cell populations.^23–26^

In the inverse problem of inferring the theoretical vector field from data, this framework has been primarily applied to molecular data such as single-cell RNA sequencing. The measured dynamic behavior of many cells can be used to estimate this theoretical vector field as a data-driven vector field.^21,22,27,28^ Since these data are snapshots, inferring a dynamical systems representation requires assumptions about change over time (i.e., cell state velocities) in the absence of directly measured dynamics.^29–31^

Although this framework was developed for molecular cell states, it can be generalized to a broader selection of cell state variables. Cell types and states have historically been defined by a cell’s observed phenotype: its appearance, location, and function.^32^ Molecular markers have since been mapped onto these observed phenotypes to create molecular representations of phenotypic cell states.^33–35^ Many aspects of the cell phenotype, including morphology, intracellular organization, and behavior are simultaneously accessible via timelapse microscopy. This modality offers an additional advantage: the cell state velocities used to build the data-driven vector field can be measured directly rather than inferred.

This approach has already been applied to image-based phenotypic cell state during cell fate decision processes,^31^ cellular response to external stimuli,^36,37^ and the epithelial-to-mesenchymal transition (EMT).^28,38,39^ Although the biological contexts and the exact methods vary, each of these works build a data-driven vector field and use it to characterize the cell state transition in question as a bifurcation in which one cell state destabilizes over time while another stable state emerges.^28,31,36–39^ We build on these works in two ways. First, we use unsupervised ML to extract biologically interpretable cell state variables from image data rather than from segmentation-based features. Second, we use a model system in which, unlike EMT or cell differentiation, the underlying biological parameters driving the state change are not inherently time-dependent and are experimentally controllable.

We selected endothelial cells cultured under shear stress as a model system because distinct image-based phenotypic cell states can be induced by changing the shear stress magnitude. Previous studies in primary endothelial have shown that different shear stress levels induce distinct orientations, parallel at low shear stress and perpendicular at higher shear stress, while maintaining endothelial identity.^40,41^ These controllable states let us investigate morphological dynamics within each state and how they change between states. Another advantage is that the flat morphology of endothelial cells allows most spatial data to be analyzed in 2D.^42^ We chose to work specifically with human induced pluripotent stem cell-derived endothelial cells (hiPSC-ECs), given their broad applicability in drug testing, disease modeling, and developmental studies.^43–46^

## Results

### hiPSC-derived endothelial cells exhibit distinct image-based cellular phenotypes under 6 dyn/cm^2^ and 21 dyn/cm^2^ fluid shear stress

We aimed to develop an imaging assay in which we could observe distinct phenotypic cell states of hiPSC-ECs based on shear stress magnitude. Previous studies have extensively demonstrated that primary vascular endothelial cells align parallel when cultured under many magnitudes of shear stress, the exact values of which are dependent upon the cell type used.^40,47,48^ In addition to aligning with fluid flow, these cells have also been found to elongate and migrate collectively, although the direction of migration varies between specific cell systems.^47,49,50^ A small number of studies indicate that some primary endothelial cell types align perpendicular when cultured at higher shear stress magnitudes.^40,50–52^ In hiPSC-ECs, parallel alignment has been observed in shear stress levels ranging from 0.4 to 15.4 dyn/cm^2^,^53–55^ and perpendicular alignment has not been reported. Human umbilical vein endothelial cells have been reported to align perpendicular when cultured in shear stress levels above 20 dyn/cm^2^.^40^ Based on these results, we first studied cells under 6 dyn/cm^2^, to induce parallel alignment and 21 dyn/cm^2^, to promote perpendicular alignment.

We generated a cell line for the Allen Cell Collection (https://cell-catalog.allencell.org/) in which VE-cadherin is endogenously tagged with monomeric enhanced green fluorescent protein (mEGFP) at its C-terminus to permit quantification of the morphology and behavior of hiPSC-ECs. We acquired fluorescent and brightfield channel timelapse images of cultured monolayers of hiPSC-ECs under continuous fluid flow in 3D every 5 minutes for 48 hours. We observed that within 18 hours of application of fluid flow, hiPSC-ECs adopted consistent phenotypes of cell behavior and morphology that persisted for hours. We identified this duration of a given timelapse dataset as the “steady state” period (Methods).

Under 6 dyn/cm^2^ shear stress, we observed that cells elongate and orient parallel to the direction of the fluid flow and collectively migrate upstream relative to the direction of fluid flow, as had been previously reported in primary endothelial cells^47,49,50^ (Figure 1A). In contrast, under 21 dyn/cm^2^ shear stress, we observed that cells still elongate but orient perpendicular to the flow direction and migrate bidirectionally and largely perpendicular to the direction of fluid flow with less coordination in migration (Figure 1A). We additionally observed differences in VE-cadherin organization between the 6 dyn/cm^2^ and 21 dyn/cm^2^ conditions. Under 6 dyn/cm^2^ the cells exhibit persistent “blobs” of mEGFP-tagged VE-cadherin polarized to the downstream edge that are brighter than the surrounding junctional fluorescent signal (Figure 1A). In contrast, under 21 dyn/cm^2^ the cells present a more homogenous VE-cadherin distribution along the periphery of the cell, with only transient polarized blobs (Figure 1A). These blobs are reminiscent of previously described organizational centers in primary endothelial cells exposed to shear stress, although VE-cadherin has not previously been observed localizing to these domains.^56–60^ Due to these dramatic visual differences in cell morphology, migratory behavior and VE-cadherin organization observed in hiPSC-ECs exposed to 6 or 21 dyn/cm^2^ shear stress we consider hiPSC-ECs at these two different shear stress magnitudes to be in distinct image-based phenotypic cell states.

**Figure 1.**
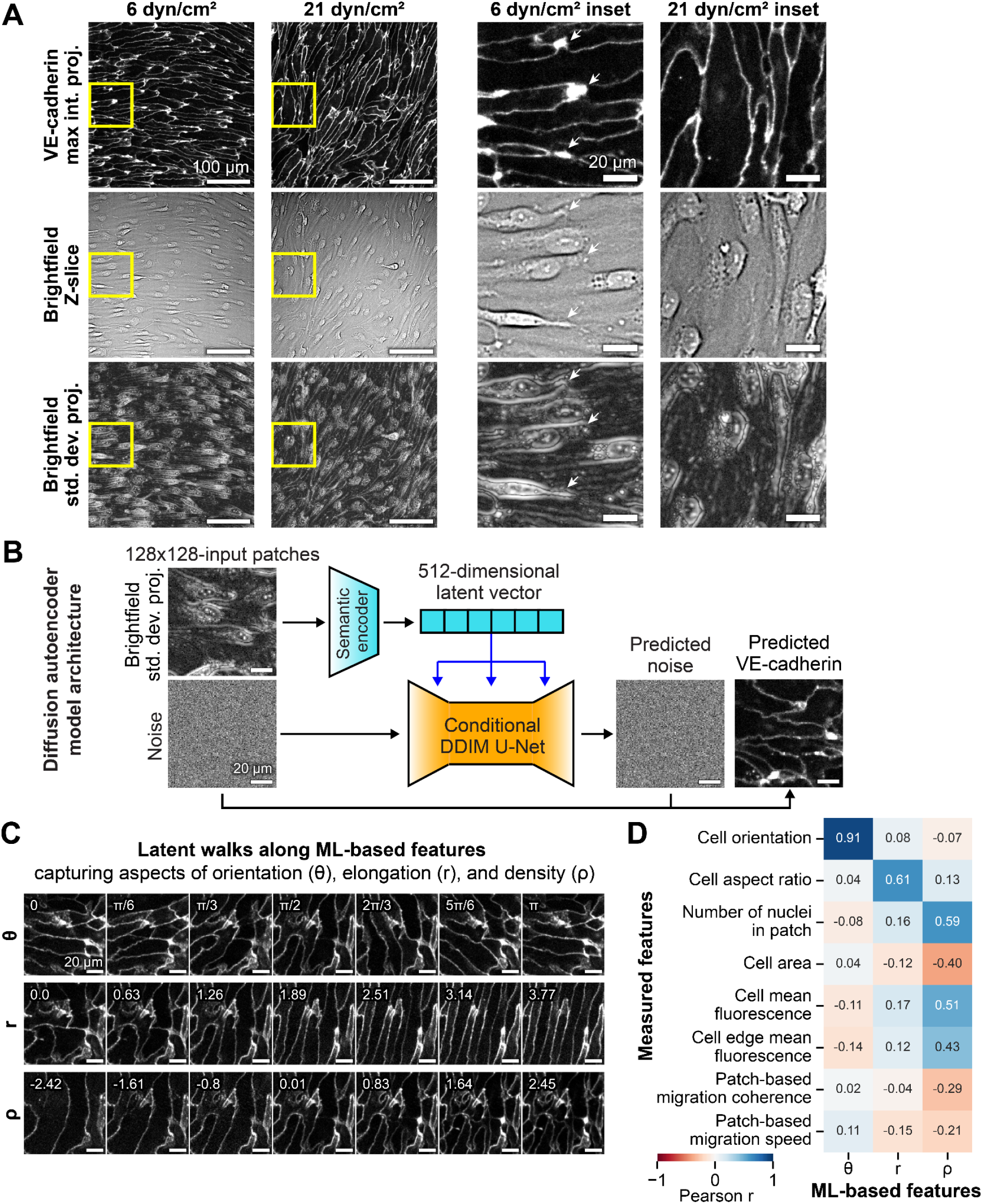
Machine learning-derived image features capture biologically relevant phenotypes of hiPSC-derived endothelial cells exposed to shear stress. **A.** 3D timelapse imaging of mEGFP-tagged VE-cadherin and brightfield was performed over 48 hours while hiPSC-ECs were exposed to shear stress. Shown are representative fields of view (FOVs) and corresponding inset regions (yellow) of cells in steady state when cultured under shear stress of 6 dyn/cm^2^ and 21 dyn/cm^2^. Downstream analysis of 2D imaging data was performed on VE-cadherin maximum intensity Z-projections and brightfield standard deviation Z-projections (same as displayed here). A corresponding brightfield Z-slice is also shown for reference. Fluid flow direction is from left to right in all images. The associated VE-cadherin maximum intensity Z-projection and brightfield standard deviation Z-projection timelapses stitched across all FOVs in the 6 dyn/cm^2^ and 21 dyn/cm^2^ replicates shown are provided in Videos S1, S4. The associated VE-cadherin maximum intensity Z-projection, brightfield Z-slice, and brightfield standard deviation Z-projection timelapses of the single FOVs and insets in the 6 dyn/cm^2^ and 21 dyn/cm^2^ replicates shown are provided in Videos S2, S3, S5, S6. White arrows on 6 dyn/cm^2^ inset images show examples of persistent VE-cadherin downstream polarized blobs. Scale bars, 100 µm (FOVs) and 20 µm (insets). **B**. Schematic showing the diffusion autoencoder (DiffAE) architecture and inference workflow. The semantic encoder compresses a 128×128 brightfield standard deviation Z-projection patch into a 512-dimensional latent representation that captures high-level morphological information. This latent representation conditions the denoising diffusion implicit model (DDIM), enabling generation of the corresponding VE-cadherin localization from an image initialized with pure noise. Scale bars, 20 µm. **C**. DiffAE-generated reconstructions of VE-cadherin patches from latent walks along interpretable ML-based features (derived in Figure S5) capturing aspects of orientation (θ), elongation (*r*), and density (ρ), spanning the range of variation observed in the *DiffAE model training dataset* for each feature. Scale bars, 20 µm. **D**. Pearson correlation coefficients (r) of interpretable ML-based features and segmentation- and patch-based measured features (for additional feature correlations see Figure S5).

### A label- and segmentation-free unsupervised ML model learns semantic image features linking brightfield to VE-cadherin

We initially pursued a single-cell segmentation-based approach to quantitatively characterize these two phenotypic cell states via the mEGFP-tagged VE-cadherin signal (Methods). Most of the relevant structural information in these relatively flat cells was confined to the XY plane. Therefore, we projected all 3D image data into 2D for downstream analysis to simplify the analyses. The VE-cadherin-mEGFP signal through the depth of the cell monolayer was captured in 2D by a maximum intensity Z-projection of the fluorescent channel (Methods, Figure 1A). The features in the brightfield image stack were enhanced and transformed into 2D through a computed log-normal standard deviation Z-projection (Methods, Figure 1A).

From single cell segmentations, we were able to directly measure many of the features that we used to visually differentiate the phenotypic cell states of hiPSC-ECs exposed to 6 and 21 dyn/cm^2^, including cell orientation, elongation, and migration angle (Methods, Figure S2).

However, segmentation accuracy for a given cell was not consistent over time, making it infeasible to track many cells over long time periods without extensive manual curation. As a result, after applying tracking algorithms only approximately 30% of the original segmentations remained (Methods, Table S5).

These limitations motivated the development of a more scalable and generalizable image-based approach for quantitatively characterizing hiPSC-EC phenotypic cell states that does not rely on cell segmentation and tracking. To learn unsupervised image representations, we used a diffusion autoencoder (DiffAE), a type of image-to-image conditioned diffusion model^61,62^ (Figure 1B, Figure S3A). Compared to standard convolutional autoencoders, diffusion models produce sharper and more realistic image reconstructions.^62,63^ In general, the network architecture of a DiffAE model consists of two jointly trained components: a denoising diffusion implicit model (DDIM) and a semantic encoder network. The trained DDIM component is capable of generating images from pure noise by learning to predict the noise added to a target image during training, effectively “denoising” the target image. The denoising process is guided by “conditioning” the DDIM on the vector obtained by passing the target reconstruction image through the semantic encoder network. This vector embedding of the image is referred to as the “semantic” or “latent” representation of the image, providing a compressed encoding of high-level image structure and morphology.

Whereas the original DiffAE implementation reconstructs the same image used as the conditioning input,^61^ we modified the training procedure to enable cross-modal prediction using paired standard deviation brightfield Z-projection (brightfield input) and VE-cadherin-mEGFP maximum intensity Z-projection (VE-cadherin target) image patches. In particular, we used the brightfield patch as the conditioning input and computed the image reconstruction loss on the corresponding VE-cadherin patch. Thus, during training the model learns two coupled tasks: encoding brightfield morphology into a 512-dimensional latent vector and reconstructing the corresponding VE-cadherin patch via a denoising process (Methods, Figure S3A). This training architecture allows for featurization of timelapse images without VE-cadherin labeling while still maintaining the potential to interpret the abstract latent representations via a prediction of the VE-cadherin signal.

We trained a DiffAE model in this manner on 128 pixels × 128 pixels (∼98 µm × 98 µm) patches from the larger timelapse images (∼6-fold larger) in the *DiffAE model training FOV dataset* (Methods, Table S2). We selected this patch size because it facilitates featurization of both a cell (average length of elongated endothelial cell: ∼70 µm) and its local neighborhood. Patches were randomly sampled across spatial locations, with a new random set drawn at each training batch. This sampling strategy increases the diversity of training examples, reduces spatial bias within the field of view, and ensures variability in whether cells are centered or fully contained within a patch, which is particularly important in a segmentation- and tracking-free approach. A subset of timelapse replicates capturing a broad range of endothelial morphologies and behaviors were selected to create the *DiffAE model training FOV dataset* (Methods, Table S2).

Once trained, this model can be used in two complementary ways. First, it encodes a brightfield patch as a compact 512-dimensional semantic latent vector representing high-level morphological features (a process called model inference), reducing the dimensionality from 128 pixels × 128 pixels (16,384 dimensions) by ∼97%. Second, the model can generate VE-cadherin patches conditioned on this latent vector, producing a reconstruction of the corresponding predicted VE-cadherin signal from a noise-initialized image (Figure 1B). We used the generative capability of this model to validate the quality of the semantic encoding by taking the latent vector encoded from a brightfield patch previously not seen by the model and using it to condition the image denoising process to transform a patch of noise into a predicted VE-cadherin patch (Methods). We then compared the resulting reconstructed VE-cadherin patch to the ground truth target VE-cadherin patch. We found that predictions successfully captured the overall cellular morphology from the real VE-cadherin signal across a wide range of VE-cadherin patterns (Figure S3B).

We performed additional controls to confirm that the morphological features in the reconstructed images are actively driven by the encoded latent vector. We first conditioned VE-cadherin reconstruction on a “scrambled” latent vector obtained by randomly permuting the latent encoding of the brightfield patch. We also conditioned reconstruction on a latent vector generated from a scrambled brightfield patch, in which pixel values were randomly shuffled (Methods, Figure S3B). Despite the realistic-looking reconstructed VE-cadherin patches generated by the model when we scrambled the latent vector, in both controls the model failed to reconstruct the morphological patterns of the VE-cadherin signal observed in the ground truth patches (Figure S3B).

These results confirm that the semantic representation of the brightfield patch encodes relevant information about the localization of VE-cadherin, enabling the prediction of a corresponding VE-cadherin pattern. To investigate how much information is lost by using brightfield rather than VE-cadherin as the DiffAE conditioning input and to estimate the upper bound on performance, we trained a separate DiffAE model with the same architecture (i.e., same number of latent dimensions, same training objective), differing only in the conditioning modality (VE-cadherin vs. brightfield). We then compared the VE-cadherin reconstruction accuracy of this VE-cadherin-conditioned DiffAE to the brightfield-conditioned DiffAE. We found that the VE-cadherin-conditioned DiffAE exhibits overall higher reconstruction quality than its brightfield-conditioned counterpart (Figure S4A, B). This result was expected since brightfield images contain only partial information about the corresponding fluorescence signal. The improved performance VE-cadherin-conditioned model did not, however, outweigh the greater generalizability of the brightfield conditioned model. The brightfield-conditioned model retained sufficient morphological information to accurately reconstruct VE-cadherin to a suitable degree for visual interpretation. We therefore selected the trained brightfield-conditioned model, hereafter referred to as the DiffAE model, for downstream analyses.

### Principal component analysis on latent feature space identifies biologically-relevant interpretable morphological state variables

Next, we aimed to characterize the information contained in the latent representations and determine whether they capture interpretable biological variation. To do this, model inference using the semantic encoder was performed on a 6x6 grid of non-overlapping patches extracted from all the timelapse images in the *DiffAE model training FOV dataset*, generating a representative set of 512-dimensional latent vectors that constitute the *DiffAE model training grid-based analysis dataset* (Methods). This grid-based sampling provides uniform coverage of the imaging field. We then fit and applied principal component analysis (PCA) to these latent vectors (Methods). We observed that the total variance is spread across many principal components (PCs), with the first 97 PCs capturing 95% of the variance and the top 10 PCs explaining only 2.04% variance on average (Figure S5A).

Because the DiffAE model can generate reconstructed VE-cadherin patches from latent vectors, the PCA-reduced feature space can be interrogated visually by reconstructing representative VE-cadherin patches at particular points in this space. To interpret the axes of biological variation captured by each PC in this manner, we performed generative “latent walks” by systematically varying one component at a time while holding the others fixed at their mean value, generating latent vectors spanning the range of each PC (Methods, Figure S5B). These latent vectors were then used as inputs to the DiffAE model to condition VE-cadherin reconstruction, providing a visual interpretation of the morphological variation captured by each PC (Figure S5B).

The latent walks along PC1 and PC2 showcase two distinct aspects of cell orientation within a patch. PC1 captures changes from parallel (0°=180°) to perpendicular (90°) orientation, while PC2 separates the “diagonal” orientations 45° and 135° (Figure S5B). In addition, cells appear more elongated at the extreme values of PC1 and PC2. The latent walk along PC3 shows variation in both the overall amount of VE-cadherin signal and the apparent cellular density within a patch, going from crowded cellular regions with extensive junctional boundaries to sparser regions with reduced visible VE-cadherin (Figure S5B). These results suggest that PC3 captures variation in local cell density at the patch scale. Visualization of PCs beyond the first three did not reveal clear or interpretable patterns (Figure S5B).

To disentangle the representation of cell orientation from that of cell elongation, we applied a polar coordinate transformation to PC1 and PC2 to obtain two new features denoted θ and *r* (Methods, Figure S5C). Furthermore, to obtain a ML-based feature representing local density, we defined an additional feature variable ρ =− *PC*3, such that higher values of ρ correspond to greater local density (Methods, Figure S5B). The latent walk along θ captures the continuum of possible cell orientations at the patch level, going from 0° to 180° (Figure 1C). At the endpoints, the reconstructions are equivalent, reflecting the circular nature of the variable. Accordingly, θ is periodic with period 180°, such that 0°=180°. The latent walk along *r* captures an increase in the elongation of cells within a patch, and the latent walk along ρ confirms the previous interpretation of PC3 as being density-related (Figure 1C).

To quantitatively validate these visual interpretations of the latent walks, we leveraged the paired measured features and ML-based features in the *DiffAE model training cell-centered analysis dataset* (Methods). We computed pairwise Pearson correlations between ML-based features and both single-cell and patch-based measured features for corresponding cell-centered patches during steady state timepoints (Figure 1D, Figure S5D).

The ML-based feature θ is strongly correlated with the measured, segmentation-based feature cell orientation (Pearson correlation, r = 0.91). In contrast, PC1 and PC2 show little correlation with cell orientation (r = 0.08 and -0.07, respectively; Figure 1D, Figure S5D). These relationships are consistent with the latent walk interpretations and indicate that the polar coordinate transformation enhances the representation of cell orientation within the latent space. While none of the original PCs correlate with the measured, segmentation-based feature of cell aspect ratio, the ML-based feature *r* correlates strongly with this measured feature (r = 0.61; Methods, Figure 1D, Figure S5D). The correlation results show that the ML-based feature ρ indeed encodes information about local cell density, specifically correlating positively with measured features related to both local cell density and VE-cadherin-mEGFP fluorescence intensity (Methods, Figure 1D, Figure S5D). PCs beyond the first three did not show strong correlations with any of the measured features (r < 0.28; Figure S5D).

These correlations indicate that the features θ, *r*, and ρ derived from the top three PCs capture key morphological properties of endothelial cells in a segmentation-free manner despite explaining only a small fraction of the total variance. While we used segmentation-based measured features to validate the morphological interpretation of these features as representations of cell morphology, it is important to remember that these representations are at the length scale of a patch, as highlighted by the generative latent walks (Figure 1C). In particular, the ML-based feature *r* does not just encode the aspect ratio (i.e., degree of elongation) of a single cell, but the degree to which all cells within a patch are elongated. Similarly, the ML-based feature ρ is a representation of the overall density of cells within a patch as inferred from 2D Z-projections of the brightfield and VE-cadherin-mEGFP images. As such, ρ is a representation of multiple entangled aspects of local cell density arising from projecting a 3D structure into 2D, including 2D-projected cell area, overall intensity in the VE-cadherin-mEGFP maximum intensity Z-projection, and overall brightness in the brightfield standard deviation Z-projection.

The axes of variation in the latent space of the trained DiffAE model did not capture several qualitative aspects of the observed hiPSC-EC phenotypic cell states. The dynamic behavioral features such as migration coherence and migration speed were not well represented by this morphological feature space (r < 0.26; Figure 1D, Figure S5D). Additionally, the observed downstream-polarized blobs were neither quantified by the cell segmentations nor systematically captured by the unsupervised learning approach. Although the DiffAE model did not give us clear representations of the organizational and behavioral aspects of the hiPSC-EC phenotypic cell states, the ML-based features θ, *r* and ρ provide complete quantitative representations of the observed morphological aspects of these cell states. We therefore defined a data-driven, morphological cell state space from these three features, permitting us to characterize the morphological aspects of the hiPSC-EC phenotypic cell states via the dynamical systems framework for cell states.

### Analysis of dynamics within ML-based morphological state space identifies shear-stress-dependent cell states as stable fixed points

#### Applying the dynamical systems framework for cell states to trajectories in ML-based morphological state space

Using the dynamical systems framework for cell states,^6–8^ cellular morphologies can be described as evolving continuously over time within a structured state space. In this general mathematical framework, the state of a cell at a given timepoint *t* is defined by a set of selected state variables that together specify its position in the state space. The collection of these locations over all timepoints in a cell’s past and future makes up a trajectory through cell state space. In the context of hiPSC-EC morphology, we approximate the full, theoretical state space of all measurable morphological properties of a cell using the data-driven state space defined by the ML-based morphological features θ, *r*, and ρ. That is, based on previous observations, we assume that the morphological aspects of cell orientation, cell aspect ratio, and local cell density are sufficient to characterize and distinguish the two observed hiPSC-EC phenotypic cell states at 6 dyn/cm^2^ and 21 dyn/cm^2^. We additionally assume that the ML- and patch-based representations of these aspects in θ, *r*, and ρ are suitable for characterizing cell states.

In this morphological state space, we represent individual points in vector form as **x** = (θ, *r*, ρ). The core assumption of the dynamical systems framework for cell states is that the paths a cell may take through the underlying state space can be described by an equation of motion for this state vector **x**.^6–8^ More formally, this equation of motion is assumed to be a stochastic differential equation,

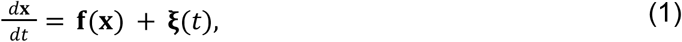

where *d***x***/dt* denotes the velocity of the state vector **x, f**(**x**) is a vector field that maps points in state space to deterministic velocities, and **ξ**(t) represents noise. The specific form of **f**(**x**) depends on the choice of state variables as well as the biological context and may vary across experimental conditions. We allow **f**(**x**) to differ across replicates and assume that it is modulated by shear stress as an external control parameter: **f**(**x**) = **f**(**x**; *shear stress*).

Some locations in cell state space have zero velocity in all directions, such that trajectories arriving at those locations remain there indefinitely. These locations are known as fixed points. In terms of the theoretical vector field **f**(**x**), a fixed point is defined as a point **x** in state space for which **f**(**x**) = 0. Fixed points can be further classified in terms of their stability, which is determined by the behavior of trajectories close to the fixed points. Stable fixed points, which we denote **x**^∗^, are those where trajectories are driven back to the fixed point following small perturbations. Formally, stability requires that the Jacobian of **f**(**x**) at the fixed point has only eigenvalues with negative real parts. These points **x**^∗^ are of interest because in the dynamical systems framework for cell states, discrete phenotypic cell states emerge exactly as the stable fixed points of the theoretical vector field **f**(**x**).^6–8^

Our goal was to use trajectory data to estimate the underlying theoretical vector field f(x) in equation (1), yielding a data-driven vector field 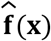 that approximates the observed dynamics in morphological state space. From 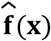, we can then identify stable fixed points **x**^∗^ (if they exist) and determine whether they correspond to observed morphological phenotypes of hiPSC-ECs. To do so, we must define what constitutes a trajectory in morphological state space, given that the state variables **x** = (θ, *r*, ρ) are patch-based and not single-cell-based.

We therefore adopted a fixed frame of reference in which image patches at constant XY locations are evaluated over time. Each patch is represented by **x**(*t*) = (θ(*t*), *r*(*t*), ρ(*t*)), yielding a trajectory in state space. These trajectories are obtained from a non-overlapping grid tiled across each field of view (*Shear stress grid-based analysis dataset*; Methods), without requiring cell segmentation or tracking. While these trajectories do not correspond to individual cells moving through space, cell motion is minimal over the 5-minute imaging interval (∼6 pixels, corresponding to ∼2% of the patch side length). Therefore, the temporal evolution of a fixed-location (grid-based) patch should closely approximate that of a cell-centered patch translating through XY-space.

#### Data-driven vector fields for representative datasets at 6 dyn/cm^2^ and 21 dyn/cm^2^ reveal dynamic relationships between cell aspect ratio and local density

For a single representative replicate at each of 6 dyn/cm^2^ and 21 dyn/cm^2^, we obtained numerical estimates of the data-driven vector field 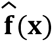 at discrete points in morphological state space using a kernel-based method applied to single-frame displacements along the grid-based trajectories from the steady state timepoints of the data (Methods, Figure S6). For each estimated vector field from these representative replicates, we found a single stable fixed point (θ^∗^, *r*^∗^, ρ^∗^) (Figure 2A). However, it was difficult to observe any obvious trends in the dynamic interdependence of θ, *r*, and ρ solely through visualization of 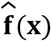 in three dimensions (Figure 2A). To get a better sense of how the variables in morphological state space depended on one another, we computed the data-driven vector field in 2D for pairwise combinations of these variables (Methods). We found that *d*θ*/dt* primarily depended on θ, and that *dr/dt* and *d*ρ*/dt* appeared qualitatively independent of θ. Therefore, we chose to compute and visualize the velocity *d*θ*/dt* as a function of θ, and the velocities *dr/dt* and *d*ρ*/dt* each as functions of (*r*, ρ). These qualitative trends held across replicates (Methods).

**Figure 2.**
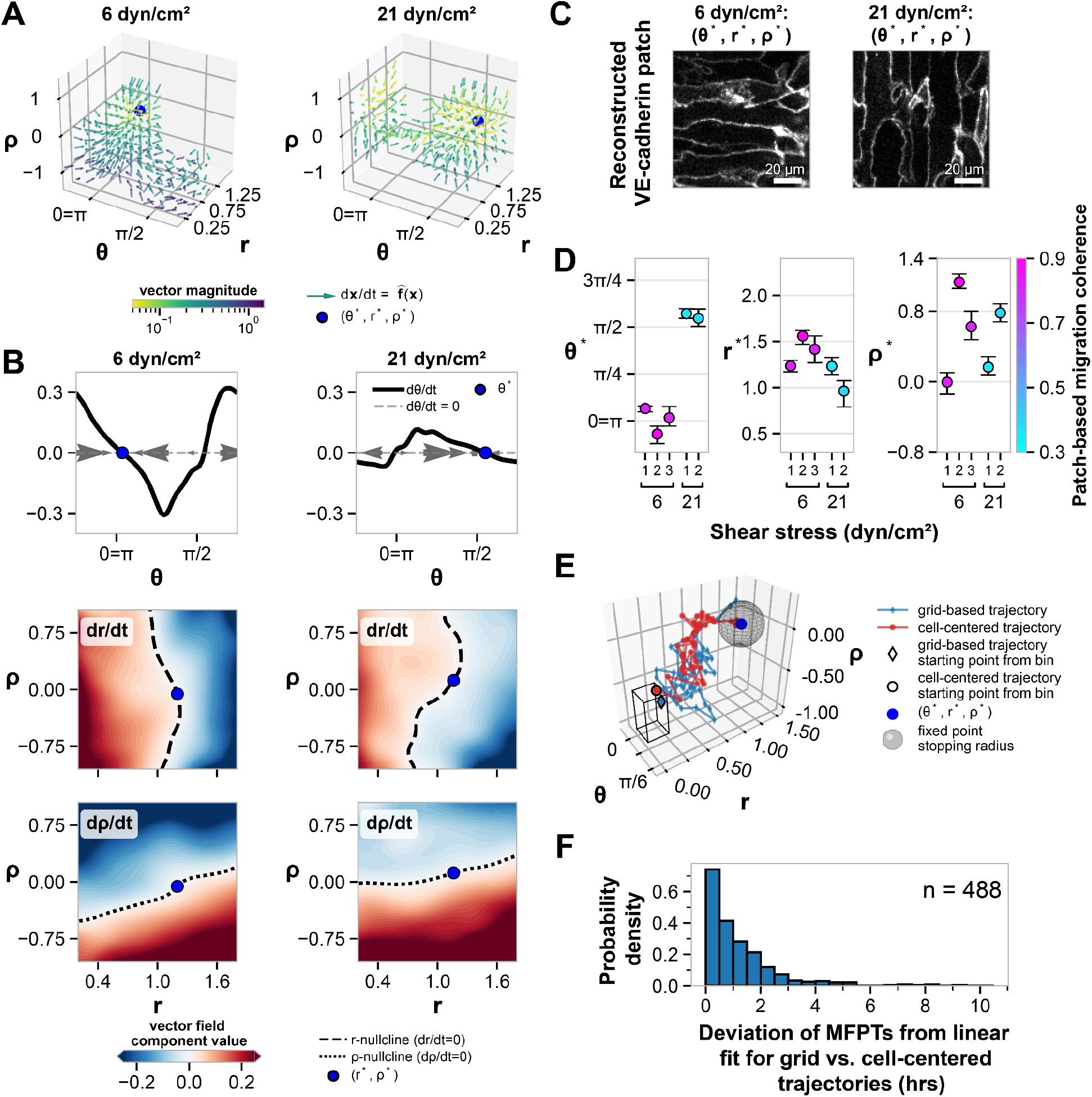
Stable fixed points of data-driven vector fields over morphological state space reveal distinct cell states at shear stress magnitudes of 6 dyn/cm^2^ and 21 dyn/cm^2^. **A.** Data-driven vector field 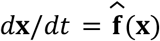 for **x** = (θ, *r*, ρ) as computed from a representative replicate at 6 dyn/cm^2^ (left) and 21 dyn/cm^2^ (right). The single stable fixed point (θ^∗^, *r*^∗^, ρ^∗^) is shown as a blue point. Arrows are normalized to unit length and indicate the local direction of the vector field, while the color indicates the magnitude of the vectors on a logarithmic scale, thresholded between 0.05 and 1.5 for visualization. **B**. Visual decomposition of the 3D vector field 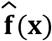 at 6 dyn/cm^2^ (left) and 21 dyn/cm^2^ (right) into 1D and 2D component parts. The topmost plot shows 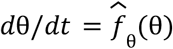 with stable fixed point θ^∗^ shown as a blue point. The black line shows the value of *d*θ*/dt* at all values of θ. The grey arrows (plotted along the line *y* = 0; dashed gray line) indicate the values (direction and magnitude) of *d*θ*/dt* at equally spaced values of θ to indicate movement along the θ axis towards the stable fixed point (phase line diagram). The lower two plots show contour plots of the values of 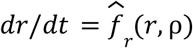 and 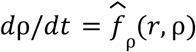, respectively, as heatmaps with stable fixed point (*r*^∗^, ρ^∗^) shown as a blue point. The range of the colormap shared by these contour plots is thresholded to be between -0.25 and 0.25 for ease of visualization. The nullcline curves defined by *dr/dt* = 0 and *d*ρ*/dt* = 0 are shown as black dashed (*r*-nullcline) and black dotted (ρ-nullcline) lines, respectively, overlaid on the contour plots. **C**. Example reconstruction of a VE-cadherin patch using the stable fixed point (θ^∗^, *r*^∗^, ρ^∗^) at 6 dyn/cm (left) and 21 dyn/cm (right) as the conditioning vector. **D**. Identified stable fixed points (θ^∗^, *r*^∗^, ρ^∗^) across replicates at 6 dyn/cm^2^ and 21 dyn/cm^2^, colored by the measured migration coherence at the stable fixed points (Methods, Figure S8D). Error bars show bootstrapped 90% confidence intervals (Methods). Panels **A**-**C** show representative examples corresponding to 6 dyn/cm^2^ replicate 1 and 21 dyn/cm^2^ replicate 1. **E**. An example of a grid-based trajectory (blue) and a cell-centered trajectory (red) moving from an example initial position (black outlined point and diamond marker, respectively) in the same bin (black box; uniform widths of 0.26 rad, 0.25 a.u., and 0.5 a.u. respectively in θ, *r*, and ρ) to a stable fixed point (blue point). The trajectory is considered to have “reached” the fixed point once it passes a given radius (0.25 in each direction) around the fixed point (grey sphere). These trajectories were used to calculate the mean first passage times (MFPTs) to the stable fixed point from each bin. **F**. Probability density of residuals from the linear fit between grid-based and cell-centered MFPTs across all replicates at 6 dyn/cm^2^ and 21 dyn/cm^2^ (see Figure S9; Methods). Residuals are reported in hours. N = 488 initial-state bins.

The 1D velocity field in θ shows the existence of single stable fixed point in terms of θ, denoted θ^∗^, at both 6 dyn/cm^2^ and 21 dyn/cm^2^ (Figure 2B). The stable fixed point at 6 dyn/cm^2^ is located at θ*≈ 0, which corresponds to parallel orientation of cells relative to the direction of fluid flow within a patch (Figure 2B). The stable fixed point at 21 dyn/cm^2^ is located at θ*≈π*/*2, corresponds to perpendicular orientation of cells relative to the direction of fluid flow within a patch (Figure 2B). Thus, not only are there single stable fixed points at each of 6 dyn/cm^2^ and 21 dyn/cm^2^, their locations in terms of θ correspond to empirical observations of cell orientation.

We found that the 2D vector fields in *r* and ρ at 6 dyn/cm^2^ are qualitatively similar to those at 21 dyn/cm^2^. The velocity *dr/dt* shows the existence of a stable value of *r* as a function of ρ, as evidenced by the stable *nullcline* (*dr/dt* = 0) dividing the (*r*, ρ)-plane (Figure 2B). Walking along this *r*-nullcline from low to high values of ρ, the corresponding *r*-coordinate first increases with ρ and then decreases with ρ past a certain value of ρ (Figure 2B). This trend points to a tradeoff between increasing local density and the corresponding degree of cell elongation within a patch. On the one hand, cells might respond to increasing local density by “compacting” along their short axis, leading to a more elongated, higher cell aspect ratio. On the other hand, the degree of local density could be so extreme that cells also get compacted along their long axis, leading to a less elongated, lower cell aspect ratio. To validate this interpretation, we leveraged the generative aspect of the DiffAE to perform a latent walk from negative to positive values along the *r*-nullcline, using the latent vectors corresponding to equally spaced points about the stable fixed point (*r*^∗^, ρ^∗^) along the nullcline to condition the image generation (Figure S7A, C). Doing so, we see that as we walk along the *r*-nullcine for increasing values of ρ the elongation of cells within a patch initially becomes more pronounced (Figure S7B, D). Once ρ is sufficiently large, the VE-cadherin pattern resembles that of cell crowding, a pattern that includes less prominently elongated cells.

The velocity *d*ρ*/dt* similarly shows the existence of a stable value of ρ as a function of *r*, again evidenced by a stable nullcline (*d*ρ*/dt* = 0) dividing the (*r*, ρ)-plane (Figure 2B). Unlike the *r* -nullcline, the ρ-nullcline shows a moderate upward trend of the stable value of ρ with increasing *r* (Figure 2B). This trend is intuitive given that the more elongated cells are, the more cells can fit in a fixed 2D area (i.e., a patch). As we did with the *r*-nullcline, we performed a generative latent walk along the ρ-nullcline through the stable fixed point (*r*^∗^, ρ^∗^) to validate this morphological interpretation of the shape of the nullcline (Figure S7A, C). We see that as we walk along the ρ-nullcine for increasing values of *r*, the local density of cells within a patch indeed increases as cell aspect ratio increases (Figure S7B, D).

Although these representative replicates at 6 dyn/cm^2^ and 21 dyn/cm^2^ differ in the location of their stable fixed points in the θ-component, the locations of their stable fixed points in *r* and ρ components have similar values of *r*^∗^ ≈ 1 and ρ^∗^ ≈ 0. Within the range of all replicates in the *DiffAE model training grid-based analysis dataset*, these values correspond to patches that contain cells with an elongated morphology and a moderate level of local density (Figure 1C, Figure 2C). When we compare the locations of the identified stable fixed points across all replicates at 6 dyn/cm^2^ and 21 dyn/cm^2^ (*Shear stress grid-based analysis dataset*; Methods) we see that at 6 dyn/cm^2^, locations θ^∗^ are consistent with parallel orientation of cells relative to fluid flow (θ^∗^ ≈ 0 for all replicates; Figure 2D). Across replicates at 21 dyn/cm^2^, we found that the locations θ^∗^ are consistent with perpendicular orientation of cells relative to fluid flow (θ^∗^≈ π*/*2 for all replicates; Figure 2D). Across replicates for both shear stress magnitudes, the locations *r*^∗^ take on a range of values close to 1 and greater, which is consistent with a more elongated phenotype (Figure 2D). The lowest stable extent of cell elongation within a patch, as quantified by *r*^∗^, observed at 21 dyn/cm is lower than that observed at 6 dyn/cm (Figure 2D). However, we observe replicates at both 21 dyn/cm^2^ and 6 dyn/cm^2^ that have the same value of *r*^∗^ (overlapping 90% confidence interval; Methods, Figure 2D). Thus, we cannot conclusively say that the state variable *r* separates the two phenotypic cell states. The values of ρ^∗^ do not show a clear trend with respect to shear stress and vary considerably across all replicates. This variation in the stable local density may arise from experimental and/or biological sources, such as initial plating density or cell proliferation rates and does not separate the two phenotypic cell states.

#### Mapping dynamic measured features of cell migration onto stable fixed points in morphological state space enhances characterization of cell states

The ML-based features θ, *r*, and ρ successfully capture morphological aspects of the two observed phenotypic cell states but do not capture the observed presence or absence of the downstream-polarized VE-cadherin blobs (Figure 1A, Figure S5B) and the notable collective migration that these cells display. Upon closer qualitative assessment we found that VE-cadherin blobs were associated with retraction fibers and migration behavior (Figure S8A). We also observed that blobs occur persistently wherever cell populations migrate collectively, whereas they only occur transiently where collective migration is absent. To quantify the degree of collective migration at the patch level, we measured optical flow between consecutive brightfield standard deviation Z-projection images and averaged normalized per-pixel flow directions within each patch to obtain a measure of migration coherence (Figure S8B). Values close to 1 indicate coherent migration in a common direction, whereas values close to 0 indicate directional cancellation consistent with incoherent or isotropic motion (Methods, Figure S8A). Using this patch-based measured feature, we visually confirmed the association between migration coherence and persistent downstream-polarized blobs, supporting its use as a proxy for their emergence and persistence (Figure S8C).

Because migration coherence is computed from brightfield patches, it can be directly mapped onto the morphological cell state space. We quantified the mean migration coherence within a small region around each fixed point (θ^∗^, *r*^∗^, ρ^∗^) (Methods, Figure S8D). At 6 dyn/cm^2^, the stable fixed point found for each replicate corresponds to high migration coherence values (∼0.9; Figure 2D, Figure S8B), near the upper bound observed in the *Shear stress grid-based analysis dataset* (95th percentile = 0.9; Methods). In contrast, at 21 dyn/cm^2^, the stable fixed point found for each replicate corresponds to low coherence values (∼0.4; Figure 2D, Figure S8B), approaching the lower bound observed for the migration coherence feature (5th percentile = 0.3; Methods). This observed lower bound likely reflects within-cell pixel-level coherence that persists even in the absence of coordinated multicellular motion. Overall, we found that the brightfield-derived, patch-based measured feature of cell migration coherence separates the observed cell migration behavior at 6 dyn/cm^2^ versus 21 dyn/cm^2^ into two distinct quantitative regimes. These two regimes of high and low migration coherence correspond to the distinct stable fixed points in the morphological state space at 6 dyn/cm^2^ and 21 dyn/cm^2^.

#### Grid-based dynamics capture behavior of features from cell-centered patches across timescales

We have defined the set of patch-based features **x** = (θ, *r*, ρ) as a morphological cell state space for hiPSC-ECs, and used grid-based trajectories in this state space to identify discrete cell states as stable fixed points of a data-driven vector field. Underlying this characterization is the assumption that both the patch-based state variables and the corresponding grid-based trajectory velocities in these variables accurately capture what cells are doing at the population level. To address whether choosing the grid-based frame of reference over the cell-centered frame of reference for quantifying cell state velocities introduces systematic mischaracterization of population-level cell state dynamics, we performed a statistical comparison of the trajectories from the two frames of reference.

We leveraged the subset of reliably tracked cell segmentations to obtain cell-centered patch trajectories and compared them with the grid-based patch trajectories. Rather than directly comparing the paths of noisy individual trajectories, we tested whether trajectories from both frames of reference approach the stable cell state on similar timescales. Specifically, we computed the average time it took grid-based and cell-centered trajectories from the same starting binned location in morphological state space to reach the stable fixed points (θ^∗^, *r*^∗^, ρ^∗^) of the grid-based data-driven vector field 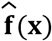 at each of 6 dyn/cm^2^ and 21 dyn/cm^2^. This quantity, known as the mean first passage time (MFPT), extends the analysis of cell state dynamics beyond the single-frame displacements used to construct the data-driven vector field 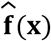 by considering the behavior of trajectories over their full duration. If the MFPT from the grid-based trajectories does not correlate with that from the cell-centered trajectories, it would indicate that grid-based data-driven vector fields (and their fixed points) are insufficient to capture the dynamic behavior of the cell-centered trajectories.

For each of the datasets at 6 dyn/cm^2^ and 21 dyn/cm^2^, we discretized morphological state space into bins and computed the MFPT from that bin as the mean time it took for trajectories starting from that location to “reach” the corresponding stable fixed point (Figure 2E). A trajectory was considered to have reached the fixed point when it entered a small radius around it (Methods, Figure 2E). We computed the MFPT per-bin in this manner separately for grid-based and cell-centered trajectories and computed the Pearson correlation between these two MFPTs across all bins within each replicate (Methods, Figure S9A, B). For 6 dyn/cm^2^ and 21 dyn/cm^2^, we found that the MFPTs were correlated across replicates with Pearson correlation values ranging from ∼0.7-0.8 (Figure 2F, Figure S9E). Furthermore, linear regression revealed grid-based trajectories exhibit comparable or slower MFPTs than the cell-centered trajectories with line fit slopes ranging from ∼0.5-0.9 for 6 and 21 dyn/cm^2^ replicates (Methods, Figure S9F).

Overall, the dynamical systems representation of the DiffAE features derived from grid-based trajectories provides a robust approximation of cell-centered trajectory dynamics across timescales. In regards to the slight discrepancies between frames of reference (deviations from 1.0 in the Pearson correlations of ∼0.2-0.3 and regression fit slopes of ∼0.1-0.5), there are several possible contributions. Cell-centered trajectories may follow more direct paths through morphological state space than grid-based trajectories because they remain centered on the same cell rather than sampling a fixed spatial location, an effect that may become pronounced over longer timescales (Figure S9A). Residual tracking errors in cell-centered trajectories may also introduce artificial jumps in state space. Together, these factors may contribute to the observed deviations in the correlation and linear fit of the MFPTs.

### A region of bistability at intermediate shear stress magnitudes is consistent with a bifurcation of cell states

We have demonstrated that by creating an ML-based morphological state space and applying a dynamical systems framework to grid-based patch trajectories in that state space, we can quantitatively characterize and differentiate the phenotypic cell states of hiPSC-ECs at shear stress magnitudes of 6 dyn/cm^2^ and 21 dyn/cm^2^. The existence of these distinct phenotypic cell states raises the question of how the transition between these states occurs as a function of shear stress magnitude. In terms of the dynamical systems framework, we are asking how exactly the underlying vector field over morphological state space depends on shear stress as a control parameter: **f**(**x**) = **f**(**x**; *shear stress*). This conceptual framing of a state transition being driven by a change in the theoretical vector field that describes cell state dynamics has been used to study cell state transitions in the contexts of both developmental^19,20,30,64–68^ and cancer biology.^23,25,26^

We hypothesized two possible mechanisms for this shear-stress-dependent transition: a continuous transition or a bifurcation (Figure 3A). In the case of a continuous transition, the underlying vector field would have a single fixed point whose location smoothly traverses the morphological state space over the range of intermediate shear stress magnitudes (Figure 3A, bottom). In a bifurcation, there would be a critical shear stress magnitude between 6 dyn/cm^2^ and 21 dyn/cm^2^ at which the underlying vector field would undergo a drastic structural change, such as the emergence of multiple, coexisting fixed points across intermediate shear stress magnitudes (Figure 3A, top). A conceptually simple possibility for the bifurcation scenario is one where the underlying dynamical system would undergo a series of saddle-node bifurcations going from 6 dyn/cm^2^ to 21 dyn/cm^2^, where the distinction is clear for which of the stable fixed points is the continuation of the 6 dyn/cm^2^-like state versus the 21 dyn/cm^2^-like state (Figure 3A, top). In practice, because of the high dimensionality of the system along with the potential influence of any number of unobserved biological parameters, the topology of the bifurcation could be more complex.^69^ Even in the idealized scenario of a saddle-node bifurcation, the locations of the stable fixed points can drift as the system approaches a critical point. For example, the continuation of the stable fixed point observed at 6 dyn/cm^2^ could exhibit an increasing value of θ^∗^ as shear stress increases, such that cells would maintain other characteristics of the 6 dyn/cm^2^ state without orientation parallel to the direction of fluid flow. This expected increased variation of the locations of stable fixed points poses a challenge for identifying the mechanism by which the 21 dyn/cm^2^ state emerges as shear stress increases.

**Figure 3.**
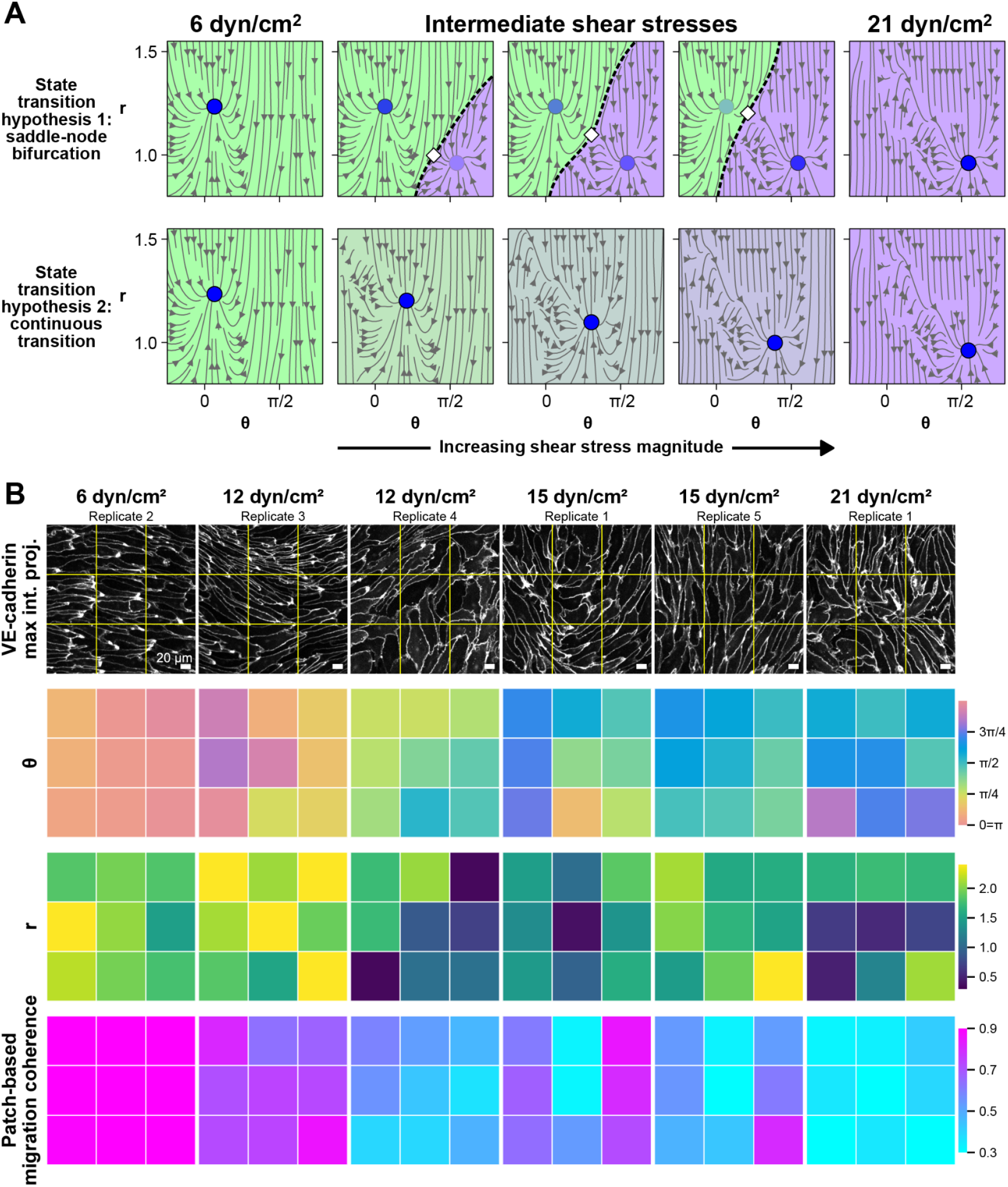
Hypothesized dynamics and imaging data characterizing the transition between 6 dyn/cm^2^ and 21 dyn/cm^2^ shear stress states. **A.** Two hypothesized mechanisms for the transition between the 6 dyn/cm^2^ and 21 dyn/cm^2^ states. 2D streamplots (grey arrows) for a single section through morphological state space are shown for illustrative purposes. **Top:** Cell state bifurcation hypothesis, in which a second stable fixed point (blue point) emerges at intermediate shear stresses. The two stable fixed points coexist and are separated by a saddle point (white diamond) and its stable manifold (black dashed line), which partitions state space into distinct basins of attraction (shaded in green and purple). The transparency of the blue points indicates the emergence and subsequent disappearance of stable fixed points, ultimately leaving the single stable 21 dyn/cm^2^ state. **Bottom:** Continuous-transition hypothesis, in which a single stable fixed point (blue point) moves smoothly through state space as shear stress magnitude changes, resulting in a gradual transformation of the vector field without the appearance of additional stable fixed points. The green-to-purple color gradient illustrates the continuous transition from the 6 dyn/cm^2^ state to the 21 dyn/cm^2^ state while maintaining a single fixed point and basin of attraction. **B**. Representative mEGFP-tagged VE-cadherin maximum intensity Z-projections at steady state timepoints from 48-hour 3D timelapse imaging of hiPSC-ECs at 6 dyn/cm^2^, 12 dyn/cm^2^, 15 dyn/cm^2^, and 21 dyn/cm^2^ shear stress magnitudes are shown (for brightfield standard deviation Z-projections see Figure S10A). The associated VE-cadherin maximum intensity Z-projections and brightfield standard deviation Z-projection timelapses stitched across all FOVs for the 6 dyn/cm^2^, 12 dyn/cm^2^, 15 dyn/cm^2^, and 21 dyn/cm^2^ replicates are provided in Videos S1, S4, S8, S10, S12, S14. The associated VE-cadherin maximum intensity Z-projection, brightfield Z-slice, and brightfield standard deviation Z-projection timelapses of the shown images for the 6 dyn/cm^2^, 12 dyn/cm^2^, 15 dyn/cm^2^, and 21 dyn/cm^2^ are provided in Videos S7, S9, S11, S13, S15, S16. Fluid flow direction is from left to right in all images. Scale bars, 20 µm. Grid-based patch locations are shown in yellow, with corresponding ML-based and measured features for each patch shown below. θ is colorized using a circular colormap, *r* using the viridis colormap, and migration coherence using a diverging colormap from magenta (high) to cyan (low; for ρ see Figure S10A).

To test whether the continuous transition or the bifurcation is the more likely scenario, we imaged hiPSC-ECs at intermediate shear stress magnitudes of 9 dyn/cm^2^, 12 dyn/cm^2^ and 15 dyn/cm^2^. At 9 dyn/cm^2^, cells are qualitatively indistinguishable from cells exposed to 6 dyn/cm^2^, exhibiting elongated morphology and cell orientation parallel to the direction of fluid flow, coherent migration upstream relative to fluid flow, and the presence of persistent downstream-polarized VE-cadherin blobs. In contrast, at 12 dyn/cm^2^ and 15 dyn/cm^2^, cells display increased phenotypic heterogeneity. To visualize this heterogeneity, we complemented the image data by color-coding the corresponding patch locations with ML-based and measured features (Figure 3B, Figure S10A).

In some 12 dyn/cm^2^ replicates, the observed phenotype closely resembles that at 6 dyn/cm^2^. In these replicates, cells orient parallel to the direction of fluid flow (θ ≈ 0, π), elongate (*r* > 1), and exhibit persistent downstream-polarized VE-cadherin blobs consistent with high migration coherence (> 0.6; Figure 3B, 6 dyn/cm^2^ in column 1 and 12 dyn/cm^2^ in column 2). Similarly, some 15 dyn/cm^2^ replicates closely resemble the phenotype observed at 21 dyn/cm^2^, with cells that orient perpendicular to the direction of fluid flow (θ ≈ π*/*2), elongate (*r* > 1), and exhibit transient polarized VE-cadherin blobs consistent with low migration coherence (< 0.6; Figure 3B, 15 dyn/cm^2^ in column 5 and 21 dyn/cm^2^ in column 6). Unlike the 6 dyn/cm^2^ and 21 dyn/cm^2^ conditions, where phenotypes were consistent across the six FOVs within each replicate, some 12 dyn/cm^2^ and 15 dyn/cm^2^ replicates exhibit increased heterogeneity both between FOVs and within individual FOVs, with neighboring local regions displaying distinct phenotypes (Figure 3B, columns 3 and 4).

In other 12 dyn/cm^2^ and 15 dyn/cm^2^ replicates, we observed substantially greater variation in cell orientation and migration coherence within individual replicates. In these individual replicates, the observed cell orientations span the full range from parallel to perpendicular relative to the direction of fluid flow, rather than being concentrated around a single preferred orientation (Figure 3B, row 2, columns 3 and 4). Migration coherence likewise exhibits a broader distribution both within individual patches and across FOVs (Figure 3B, row 4, columns 3 and 4). In addition, spatially organized regions emerge within FOVs from these replicates, in which neighboring cells share similar orientations, exhibit greater elongation, and migrate more coherently. These spatially organized regions are in contrast to adjacent regions that contain cells with different orientations, reduced elongation, and lower migration coherence (Figure 3B, columns 3 and 4). Spatial regions containing persistent downstream-polarized VE-cadherin blobs corresponded to patches with higher migration coherence; this relationship reflects their dynamic behavior, best visualized over time (Video S10-13).

While the association between persistent downstream-polarized VE-cadherin blobs and high migration coherence is maintained at intermediate shear stress magnitudes, the coupling between cell orientation and migration coherence is disrupted. Unlike the 6 dyn/cm^2^ and 21 dyn/cm^2^ states, where orientation was associated with the degree of migration coherence, there are individual patches in the intermediate shear stress conditions that exhibit both high and low migration coherence across the full range of cell orientations (Figure 3B, rows 1 and 4, columns 3 and 4). Despite these differences in cell orientation, the observed association between cell elongation and migration coherence at 6 dyn/cm^2^ and 21 dyn/cm^2^ persisted in the intermediate shear stress regimes. Regions with elongated cells and high migration coherence resembled the 6 dyn/cm^2^ state, whereas regions with reduced elongation and low migration coherence more closely resembled the 21 dyn/cm^2^ state (Figure 3B, rows 3 and 4, columns 3 and 4). The coexistence of these distinct phenotypes within individual replicates at intermediate shear stress magnitudes suggests that the transition between the 6 dyn/cm^2^ and 21 dyn/cm^2^ states occurs through a bifurcation instead of a continuous transition (Figure 3A).

To quantitatively distinguish between the bistable and continuous-transition hypotheses, we estimated the data-driven vector field 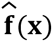 over morphological state space for grid-based trajectories from multiple replicates at 9 dyn/cm^2^, 12 dyn/cm^2^, and 15 dyn/cm^2^ (*Shear stress grid-based analysis dataset*, Methods). We then applied the same analysis used for the 6 dyn/cm^2^ and 21 dyn/cm^2^ replicates to identify the stable fixed points **x** = (θ^∗^, *r*^∗^, ρ^∗^) for each of these data-driven vector fields and quantify the measured migration coherence at each fixed point (Figure 4A). Consistent with our qualitative observations, the stable fixed points at 9 dyn/cm^2^ occupy locations similar to those observed at 6 dyn/cm^2^, with values of θ^∗^ corresponding to orientation of cells parallel to the direction of fluid flow and values of *r*^∗^ corresponding to elongated cells (Figure 4A). In contrast, there is high variability of the location and number of stable fixed points in all variables across replicates at both 12 dyn/cm^2^ and 15 dyn/cm^2^ (Figure 4A, Figure S10B).

**Figure 4.**
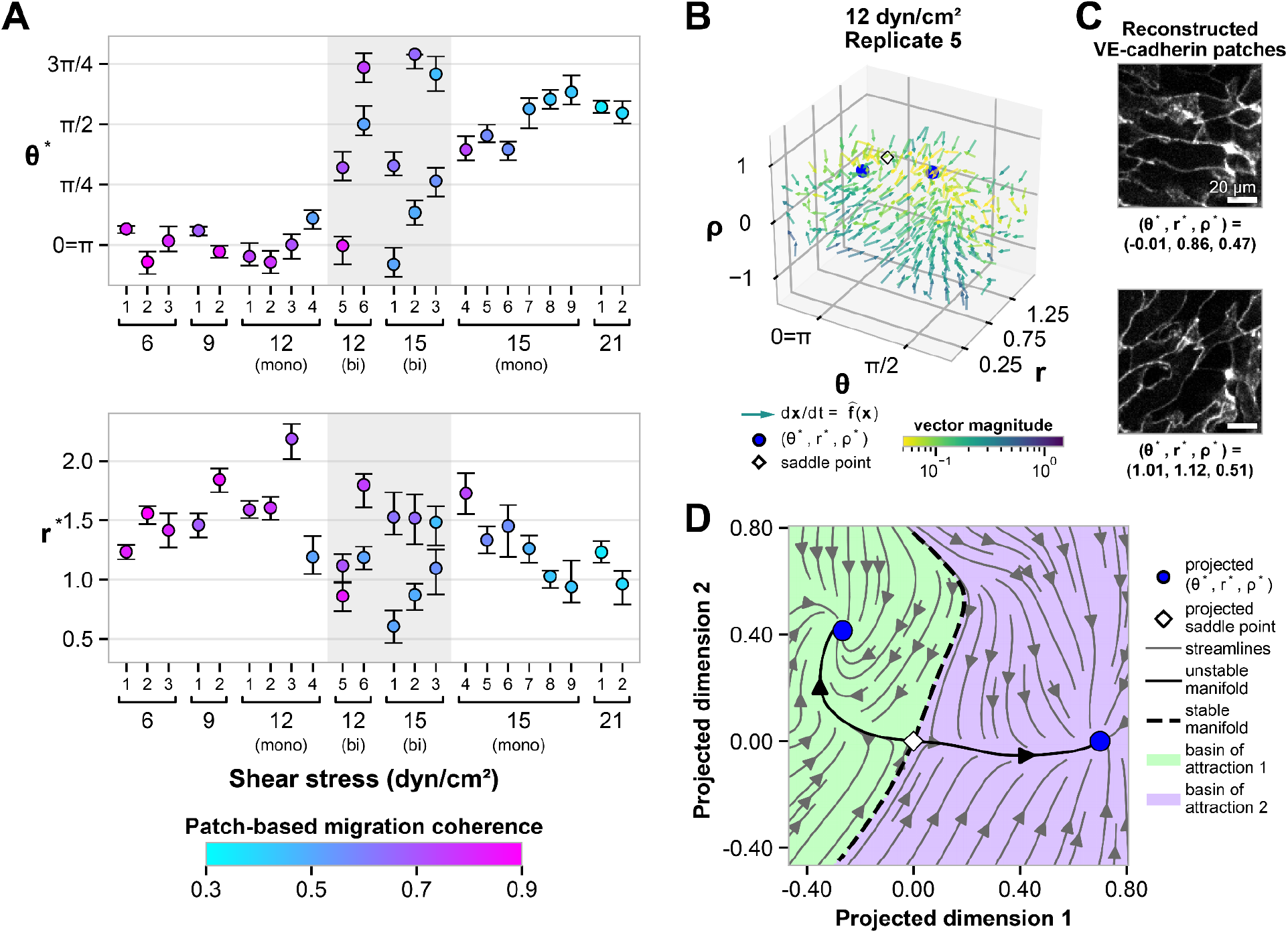
Properties of the data-driven vector field are consistent with a saddle node bifurcation at intermediate shear stress magnitudes. **A.** Identified stable fixed point locations θ^∗^ and *r*^∗^ across replicates at 6 dyn/cm^2^, 9 dyn/cm^2^, 12 dyn/cm^2^, 15 dyn/cm^2^ and 21 dyn/cm^2^, colored by the measured migration coherence at each stable fixed point for all replicates in the *Shear stress grid-based analysis dataset* (for ρ^∗^ see Figure S10B). Error bars show bootstrapped 90% confidence intervals (Methods). Replicates exhibiting bistable fixed points at 12 dyn/cm^2^ and 15 dyn/cm^2^ are highlighted by the grey shaded region. **B**. Data-driven vector field 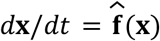 from a representative 12 dyn/cm^2^ bistable replicate (12 dyn/cm^2^ replicate 5 in **A**). The two stable fixed points (θ^∗^, *r*^∗^, ρ^∗^) are shown as blue points, and the saddle point as a white diamond. Arrows are normalized to unit length and indicate the local direction of the vector field, while the color indicates the magnitude of the vectors on a logarithmic scale, thresholded between 0.05 and 1.5 for visualization. **C**. Example reconstructions of a VE-cadherin patch using stable fixed points (θ^∗^, *r*^∗^, ρ^∗^) from representative 12 dyn/cm^2^ bistable replicate (12 dyn/cm^2^ replicate 5 in **A**) as conditioning vectors. **D**. A 2D streamplot (grey arrows) of a representative bistable 12 dyn/cm^2^ replicate (12 dyn/cm^2^ replicate 5 in **A**) is shown. The 2D view is obtained by projecting (θ, *r*, ρ)-space onto the plane spanned by the two stable fixed points (blue points) and the saddle point (white diamond). The resulting 2D streamplot is obtained by projecting the vector field 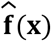 orthogonally onto this plane. The stable and unstable manifolds of the projected saddle point are shown as dashed and solid black lines, respectively. The 2D projection of 2D projection of the stable fixed point (θ^∗^, *r*^∗^, ρ^∗^) = (− 0. 01, 0. 86, 0. 47), with basin of attraction colored in green, is the leftmost stable fixed point in the plot. The 2D projection of the stable fixed point (θ^∗^, *r*^∗^, ρ^∗^) = (1. 01, 1. 12, 0. 51), with basin of attraction colored in purple, is the rightmost stable fixed point in the plot.

Of the replicates that yielded a single stable fixed point (θ^∗^, *r*^∗^, ρ^∗^), which we term “monostable”, those at 12 dyn/cm^2^ displayed fixed point locations consistent with the 6 dyn/cm^2^ cell state (i.e, θ^∗^ ≈ 0 and *r*^∗^ > 1), while those at 15 dyn/cm^2^ displayed fixed point locations consistent with the 21 dyn/cm^2^ cell state (i.e, θ^∗^ ≈ π*/*2 and *r*^∗^ ≳ 1; Figure 4A). Most monostable 12 dyn/cm^2^ replicates exhibit high migration coherence (magenta; >0.7) at the stable fixed point, and most monostable 15 dyn/cm^2^ replicates exhibit low migration coherence (cyan, <0.5) at the stable fixed point (Figure S4A). However, both shear stress conditions show monostable replicates that deviate from this trend, exhibiting either low or high coherence, respectively (Figure 4A, 12 dyn/cm^2^ replicate 4, 15 dyn/cm^2^ replicate 4). Migration coherence for monostable replicates appears related to the value of the stable fixed point location *r*^∗^, with lower *r*^∗^ values corresponding to lower migration coherence (Figure 4A). Because low migration coherence is a key characteristic of the 21 dyn/cm^2^ cell state, this result strengthens the previous observation of a subtly lower value of *r*^∗^at the 21 dyn/cm^2^ cell state compared to the 6 dyn/cm^2^ cell state (Figure 2D). Consistent with previous observations at 6 dyn/cm^2^ and 21 dyn/cm^2^, the locations ρ^∗^ across these monostable replicates did not show a clear dependence on shear stress magnitude (Figure S10B).

Among replicates at 12 dyn/cm^2^ and 15 dyn/cm^2^ that exhibit two stable fixed points, which we term “bistable”, each individual replicate exhibits two distinct stable cell orientations and degrees of elongation, as evidenced by the two stable fixed points having distinct values of ^∗^ and ^∗^ (Figure 4A). Additionally, within a given replicate each fixed point had a distinct migration coherence, and three of the five bistable replicates had one fixed point corresponding to high migration coherence and one fixed point corresponding to low migration coherence (Figure 4A, 12 dyn/cm^2^ replicate 6, 15 dyn/cm^2^ replicates 1 and 2). Bistability is not strongly reflected in the value of ρ^∗^, with the two stable states often exhibiting overlapping locations in terms of ρ (Figure S10B). This observed pattern of bistability within a given replicate is consistent with the cell state bifurcation hypothesis (Figure 3A).

To further examine how bistability manifests in the data-driven vector field, we performed an in-depth analysis of a single bistable replicate at 12 dyn/cm^2^ (Figure 4A, replicate 5). A plot of the data-driven vector field 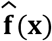 shows the two distinct stable fixed points as sinks of the vector field (Figure 4B). This replicate represents one of the simplest cases, in which θ^∗^ most clearly distinguishes the two states, while *r*^∗^ and migration coherence are similar and ρ^∗^ is indistinguishable between states (overlapping confidence intervals, Figure 4A, Figure S10B). To confirm this interpretation, we used the generative capabilities of the DiffAE model to produce VE-cadherin patch reconstructions at each of these stable fixed points. We found that the morphologies of the stable fixed points correspond to the expected difference in cell orientation between the two states (Figure 4C). From the direct visualization of 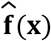 in three dimensions, however, it was difficult to further resolve how bistability structures the morphological state space in this replicate (Figure 4B). To address this, we sought a two-dimensional projection of 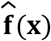 that preserves key structural features of the full three-dimensional vector field while enabling clearer visualization. We found that for the selected 12 dyn/cm^2^ replicate, an orthogonal projection of 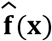 onto the plane defined by the two stable fixed points and the single saddle point between them was suitable because it preserved the local linear stability of the fixed points: the stable fixed points of 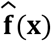 remained stable and the saddle point remained a saddle (Methods, Figure 4D). More broadly, finding a suitable projection for low-dimensional visualization of this manner is related to the non-trivial problem of selecting an appropriate reduced-order-model for a high-dimensional, non-linear dynamical system.^70,71^ The two-dimensional projection obtained for this replicate provides a clear and intuitive visualization of a bistable dynamical system, mirroring the streamline portrait for the bifurcation hypothesis in Figure 3A.

### An N-terminal truncating VE-cadherin mutation alters cell aspect ratio but not migration coherence of hiPSC-ECs exposed to 6 dyn/cm^**2**^

Given the strong correlation observed between orientation (θ^∗^) and migration coherence at 6 dyn/cm^2^ and 21 dyn/cm^2^, it was striking that at the intermediate shear stress magnitudes of 12 dyn/cm^2^ and 15 dyn/cm^2^, migration coherence and orientation could become uncoupled (Figure 4A, 12 dyn/cm^2^ replicates 4 and 5, 15 dyn/cm^2^ replicate 1 and 4). We aimed to mechanistically decouple orientation and coherent migration in hiPSC-ECs by truncating the N-terminal of VE-cadherin. The N-terminal extracellular cadherin (EC) domain, EC1, is primarily responsible for homophilic VE-cadherin binding in trans with neighboring cells to enable the formation of adherens junctions,^72–74^ and is necessary for collective cell migration in vasculogenesis.^75^ However, the EC domains are dispensable for endothelial alignment under shear stress.^76^ Therefore, we hypothesized that hiPSC-ECs expressing an N-terminal truncation of VE-cadherin and subject to 6 dyn/cm^2^ shear stress would still align with the fluid flow direction but no longer migrate coherently.

To investigate this hypothesis, we utilized a cell line in which we had deleted exon 3 in the mEGFP-tagged VE-cadherin hiPSC line (Ex3Del cells, Methods). This mutation introduced a premature stop codon, and western blots indicated that the Ex3Del cells express an N-terminal truncation product of VE-cadherin that excludes EC1 and EC2 (Figure S11A-D). Because the Ex3Del cells lacks the VE-cadherin epitope used to sort the mEGFP-tagged VE-cadherin line and were instead sorted for CD-31 expression, we compared the mutant line to both VE-cadherin-mEGFP cells sorted for VE-cadherin (VE-cadherin-mEGFP) and CD-31 expression (Sorting control) as controls. All cells were subject to 6 dyn/cm^2^ using the standard assay.

The GFP signal of Ex3Del cells is significantly decreased compared to the control cells, indicating a drastic reduction in VE-cadherin expression, although localization to downstream-polarized blobs is still observed (Figure 5A). When subjected to 6 dyn/cm^2^ shear stress, Ex3Del hiPSC-ECs align parallel to the direction of fluid flow and migrate upstream, as we had hypothesized (Figure 5A). Contrary to our hypothesis, however, the Ex3Del cells appeared to maintain coherent migration. Migration speed of Ex3Del cells also appears slightly increased compared to control cells. Compared to controls the Ex3Del cells exhibit distinct geometries and packing. Although they still appear elongated under response to shear stress, they were more polygonal in shape than the roughly ellipsoid control cells (Figure 5A). The Ex3Del cells display rounder morphologies that appear brighter in the brightfield standard deviation projection (Figure 5A). Consistent with this observation, Ex3Del cells decrease in cell density after the steady state timepoints used for analysis, contrasting with the increase in density and cell crowding observed in control data at these late timepoints.

**Figure 5.**
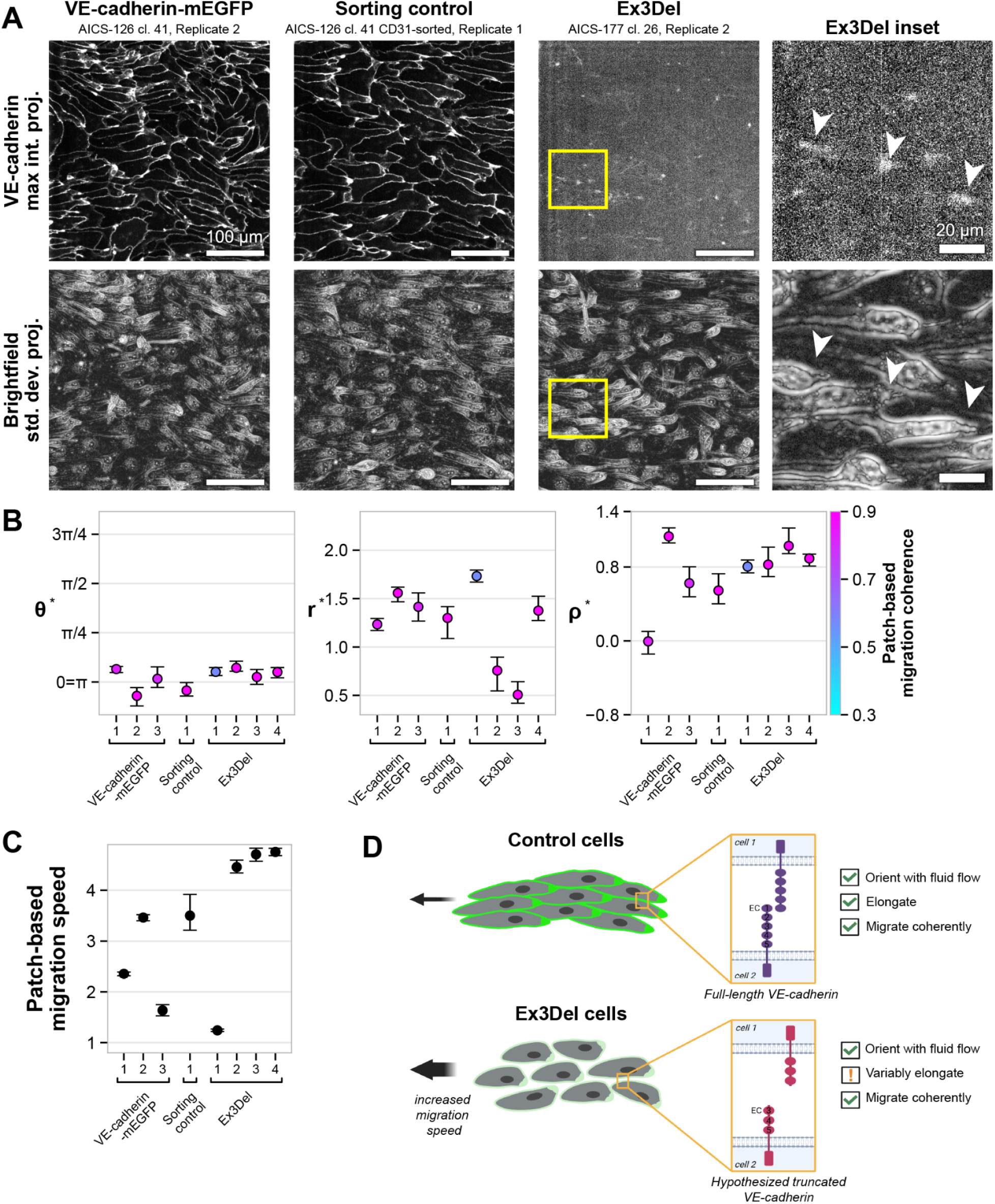
Genetic perturbation of VE-cadherin selectively disrupts features of cell state under 6 dyn/cm^2^ shear stress. **A.** Representative mEGFP-tagged VE-cadherin maximum intensity Z-projection and brightfield standard deviation Z-projection FOVs of the mEGFP-tagged VE-cadherin cell line (VE-cadherin-mEGFP), CD31-sorting control (Sorting control), and homozygous VE-cadherin exon 3 deletion line (Ex3Del) under 6 dyn/cm^2^ shear stress. The associated VE-cadherin maximum intensity Z-projection and brightfield standard deviation Z-projection timelapses stitched across all FOVs in the shown replicates are provided in Videos S1, S17, S19. The associated VE-cadherin maximum intensity Z-projection, brightfield Z-slice, and brightfield standard deviation Z-projection timelapses of the shown FOVs and inset are provided in Videos S2, S18, S20, S21. Corresponding inset regions (yellow) of Ex3Del images are shown in the rightmost column, with white arrows indicating examples of persistent downstream polarized VE-cadherin blobs. Fluid flow direction is from left to right in all images. Scale bars, 100µm (FOV) and 20µm (insets). **B**. Identified stable fixed point locations θ^∗^, *r*^∗^ and ρ^∗^ across replicates of the VE-cadherin-mEGFP (6 dyn/cm replicates in the *Shear stress grid-based analysis dataset;* Methods), sorting control, and Ex3Del cell lines (in the *VE-cadherin Exon3Del perturbation grid-based analysis dataset;* Methods) colored by the measured migration coherence at each stable fixed point. Error bars show bootstrapped 90% confidence intervals (Methods). **C**. Values of migration speed calculated at the stable fixed point for each replicate in the *VE-cadherin Exon3Del perturbation grid-based analysis dataset* (Methods). Error bars show bootstrapped 90% confidence intervals (Methods). **D**. Schematic showing overview of phenotypic differences between VE-cadherin-mEGFP and Ex3Del cells and hypothesized VE-cadherin truncation product. Schematic in **D**. generated using BioRender.

While segmentation-based quantification of this dataset was not possible due to the low GFP signal, our segmentation- and label-free approach let us extract features based solely on the brightfield channel. We then used the same method as before to estimate the data-driven vector field 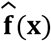 over morphological state space and calculate its stable fixed points (θ^∗^, *r*^∗^, ρ^∗^) for Ex3Del and control cells exposed to 6 dyn/cm^2^ shear stress. A single stable fixed point was identified within each control and Ex3Del replicate (Figure 5B). The values of ^∗^ are similar across cell lines and replicates and are consistent with the visually observed parallel alignment in Ex3Del cells exposed to 6 dyn/cm^2^ (θ^∗^≈ 0 ; Figure 5B).

The values of *r*^∗^ show a larger spread across replicates of Ex3Del cell populations than in the control replicates (Figure 5B). On the low end of this observed range, two of the Ex3Del replicates also displayed fixed point locations *r*^∗^ with significantly lower values than observed in control cells (*r*^∗^ ≈ 0. 5, as compared to *r*^∗^ >1 ; Figure 5B). This result is in agreement with the observation that Ex3Del cells sometimes display a reduced aspect ratio when exposed to 6 dyn/cm^2^ compared to control cells (Figure 5A). On the high end of the range of *r*^∗^ in Ex3Del cells, one replicate displayed an *r*^∗^ value that was significantly higher than any of the control replicates (*r*^∗^ ≈ 1. 75 ; Figure 5B). We hypothesize the larger range and significantly lower observed minimum value of locations *r*^∗^across Ex3Del replicates may be due to a reduction in contact inhibition in these cells.^77,78^ This reduced function has been shown to impact cell spreading^79^ and could make cell morphology more sensitive to initial experimental conditions.

Ex3Del cells displayed a smaller range of values of ρ^∗^ compared to both controls, but the full range of ρ^∗^ across Ex3Del replicates still fell within the observed range of ρ^∗^ across VE-cadherin-mEGFP and sorting control replicates combined (Figure 5B). Closer examination of images from the parental and Ex3Del datasets with similar values of ρ demonstrate that this is not a simple reflection of a more consistently high cell density (Figure S11F). Instead, we hypothesize that the DiffAE model is interpreting the rounded, slightly detached morphology of Ex3Del cells as representative of a higher value of ρ in control cells. Because the DiffAE model was not trained on any Ex3Del data, it is reasonable that the model might not correctly interpret this new cell morphology phenotype. In sum, the fixed points calculated from the Ex3Del data matched our qualitative observations of orientation (θ) and aspect ratio (*r*), but incorrectly characterized local cell density (ρ).

To further characterize changes associated with the VE-cadherin truncation, we again calculated the average measured migration features at the stable fixed points (θ^∗^, *r*^∗^, ρ^∗^). Unlike the consistent migration speed observed in the VE-cadherin-mEGFP line across all shear stress magnitudes studied (Figure S9C), the measured speed of cell migration is higher in Ex3Del cells compared to controls at 6 dyn/cm^2^ (Figure 5C). The increase in migration speed is consistent with previous reports that endothelial cells migrate faster with EC1 functionally blocked^80^ and could be attributed to the lack of cell-cell binding through VE-cadherin, freeing the cells to migrate more quickly.^49,81^ The migration coherence of Ex3Del cells is comparable to both the parental and editing controls, consistent with qualitative observations (Figure 5B). These results together suggest that the EC1 and EC2 domains of VE-cadherin are not required for the observed and measured coherent migration in hiPSC-ECs exposed to shear stress (Figure 5D).

## Discussion

In this work we use an unsupervised ML-based approach to learn biologically-relevant, interpretable representations of cell morphology, which are then combined with a dynamical systems framework to characterize cell states and cell state transitions in hiPSC-ECs under shear stress. In taking a fully unsupervised approach to ML-based feature extraction, we found that a brightfield-conditioned DiffAE model learned nuanced representations of cell orientation, aspect ratio, and local density. Although the PCs in the DiffAE model’s latent space captured important morphological variation, they did not encode aspects of cell behavior, such as migration coherence or the migration speed. Because the model was trained on static image snapshots, this finding highlights the importance of explicitly incorporating temporal information into the model training process when dynamic aspects of cell state are of interest.^82–85^ We could, however, directly map brightfield-derived measured migration features onto the morphological state space. By doing so, we showed a clear phenotypic distinction between the two cell states at 6 dyn/cm^2^ and 21 dyn/cm^2^ and additionally supported interpretation of the bifurcation in the bistable regime at 12 dyn/cm^2^ and 15 dyn/cm^2^. It is worth highlighting that this integration was possible because the model was conditioned on brightfield alone, and its learned features can be paired with virtually any other measurement acquired from the same field of view,^86^ such as immunofluorescence or transcriptomics, making it a flexible foundation for multimodal, integrative characterization of cell state.

For the Ex3Del replicates, evaluating the measured migration features at the stable fixed points revealed the surprising finding that Ex3Del cells displayed migration coherence similar to control cells. Given the reported necessity of the EC1 domain for collective endothelial migration during vasculogenesis,^75^ we propose that the coherent migration observed in Ex3Del cells may reflect cells responding similarly to their microphysiological environment, rather than undergoing canonical cadherin-dependent collective migration. While the level of expressed truncated VE-cadherin was drastically reduced, the protein also continued to localize to the observed persistent downstream-polarized VE-cadherin blobs, suggesting that the EC1 and EC2 domains are not necessary for this localization. Although blob persistence strongly correlates with coherent migration in both control and Ex3Del cells, the present data do not allow us to establish causality. In light of recent papers reporting other proteins localizing to downstream domains in endothelial cells under shear stress, this presents an intriguing subject for further investigation.^56–60^

A key benefit of using hiPSC-ECs as a model system is that the axis along which the state transition occurs, shear stress magnitude, is experimentally controllable. This allowed us to directly test whether the cell state transition going from 6 dyn/cm^2^ to 21 dyn/cm^2^ occurs through a bifurcation by quantifying how the data-driven vector field changed with shear stress. We found replicates at 12 dyn/cm^2^ and 15 dyn/cm^2^ that had two stable fixed points, which is consistent with the hypothesized bifurcation of cell states with respect to shear stress magnitude. Regarding the observed variability between replicates at 12 dyn/cm^2^ and 15 dyn/cm^2^, it is well established conceptually that as a system approaches a critical transition point (i.e., the point at which fixed points appear or disappear), the basins of attraction of a stable fixed point become shallower and wider, leading to a higher degree of variation at the population level.^67,87,88^ As the system approaches the critical shear stress magnitudes, a decrease in the relative stability of the two theoretically co-existing states would allow for other unquantified aspects of biological variation to have an increased effect on which cell states are manifest in the population. This possible sensitivity could explain the complexity of the quantified cell states across replicates at 12 dyn/cm^2^ and 15 dyn/cm^2^, including the presence of both monostable and bistable replicates as well as the decoupling of cell orientation from migration coherence.

This decoupling of key aspects of the hiPSC-EC phenotypic cell state at intermediate shear stress magnitudes highlights the need for deeper mechanistic understanding of this cell state transition. On the theoretical end, the quantitative characterization of morphological and behavioral aspects of the hiPSC-EC phenotypic cell states would benefit greatly from experimentally validated mechanistic models, akin to the well-established dynamical systems models for gene regulatory networks. A large body of work on spatiotemporal active matter models already addresses the dynamic interplay between cell orientation, cell morphology, and cell migration,^89–95^ but these models have yet to be placed in the conceptual context of cell states. On the experimental end, there is significant data shedding light on the molecular mechanisms responsible for the endothelial response to shear stress.^96–99^ Integrating this mechanistic data to achieve a multi-scale understanding of these image-based phenotypic cell states would greatly enrich the quantitative, image-based framework established in this study.

### Limitations of the study

Although hiPSC-ECs have demonstrated utility for drug testing, developmental studies, and disease modeling,^43–46^ they differ from primary endothelial cells in notable ways. They express endothelial markers and flow-sensitive genes, but often at lower levels than in primary ECs.^100,101^ hiPSC-ECs appear to respond to fluid shear stress at lower magnitudes than primary ECs: published data show parallel alignment at shear stresses as low as 0.4 dyn/cm^2^, compared with a low of 10 dyn/cm^2^ in HUVECs, and data in this study similarly show perpendicular orientation emerging at 12 dyn/cm^2^ in hiPSC-ECs versus 20 dyn/cm^2^ in HUVECs.^40,53^ They also exhibit mixed arterial and embryonic-like transcriptional identity,^102,103^ and show blunted inflammatory responses relative to primary endothelial cells.^104,105^ Thus, while they provide an excellent model system for studying morphological cell states, mechanistic conclusions attributed to canonical endothelial cells should be interpreted with these limitations in mind.

## Author Contribution Statement

Conceptualization: E.A., C.L.L., S.E.P., R.J.Z., B.M., E.A.E., C.H., J.M., B.R., J.W., J.A.T., G.D., S.M.R., M.P.V. Investigation: E.A., C.L.L., S.E.P., R.J.Z., T.B., J.C.D., B.M., R.J.D., J.H.E., E.A.E., M.J.H., J.M., E.E.S., M.L.K., J.W., J.A.T., G.D., S.M.R., M.P.V. Methodology: E.A., C.L.L., S.E.P., R.J.Z., T.B., F.M., B.M., J.A., J.H.E., J.P.T., M.P.V. Resources: C.L.L., R.J.Z., T.B., C.S.W., J. Y., J.S.Y. Validation: E.A., C.L.L., S.E.P., R.J.Z., T.B., J.C.D., F.M., B.M., R.J.D., J.H.E., M.J.H., J.M., S.S.M., A.P., E.E.S., J.S.Y. Data curation: C.L.L., S.E.P., R.J.Z., J.S.Y. Software: E.A., C.L.L., S.E.P., F.M., B.M., J.S.Y. Formal analysis: E.A., C.L.L., S.E.P., F.M., S.S.M. Visualization: E.A., C.L.L., S.E.P., F.M. Writing - original draft: E.A., C.L.L., S.E.P., R.J.Z., T.B., J.S.Y., S.M.R., M.P.V. Writing - review & editing: E.A., C.L.L., S.E.P., R.J.Z., M.L.K., E.N., J.W., J.A.T., G.D., S.M.R., M.P.V. Supervision: E.A., C.L.L., S.E.P., R.J.Z., J.C.D., C.H., B.R., M.L.K., E.N., J.A.T., G.D., S.M.R., M.P.V. Project administration: R.J.Z., C.R.G., M.P.V.

## Declaration of generative AI and AI-assisted technologies in the manuscript preparation process

During the preparation of this work, the author(s) used Claude and ChatGPT as part of the editing process. The author(s) carefully reviewed and edited the output and take full responsibility for the content of the published article.

## Acknowledgements

We thank Helen Anderson, Jamie Gehring, and Susan Ludmann for exploring the feasibility of endothelial cell culture on glass substrates. We thank Gaudenz Danuser, Margaret Gardel, Quincey Justman, and Wallace Marshall for helpful scientific discussions. The WTC-11 hiPS cell line used to create the gene-edited cell lines of the Allen Cell Collection was provided by the B. R. Conklin laboratory at the Gladstone Institute and University of California, San Francisco. This research was supported by the Allen Institute, founded by Jody Allen – chair and co-founder of Allen Family Philanthropies, and the late Paul G. Allen – investor, philanthropist, and co-founder of Microsoft. We gratefully acknowledge their vision and generosity, which make this work possible.

## Materials and Methods

### Cell lines

#### Generation of CDH5-mEGFP hiPS cell line

The Allen Cell Collection consists of gene-edited human induced pluripotent stem (hiPS) cell lines, generated from the wild-type WTC-11 background (AICS-0), in which proteins marking distinct cellular structures are endogenously fused to fluorescent proteins via gene tagging. For this study, we used AICS-0126 cl. 41, which endogenously expresses mEGFP-tagged vascular endothelial (VE)-cadherin. This cell line was newly generated in this study for the Allen Cell Collection, using AAV6 DNA donors. In brief, cells underwent nucleofection using a Lonza P3 Primary Cell 4D Nucleofector X Kit S (Cat # V4XP-3032, Lonza) and a Lonza 4D system (Lonza 4D Core Unit, Cat # AAF-1003B; Nucleofector X Unit Cat # AAF-1003X, Lonza) with Cas9/sgRNA ribonuclear proteins, then transduced with a custom designed AAV6 vector (Addgene plasmid # 217425) at 0.5×10^3^ MOI.^106^ Following gene editing, cells from the modified populations were identified with custom-designed droplet digital (ddPCR) assay surveillance.^107^ Subsequently, PCR screening was used to identify clones. AICS0-126 cl. 41 was indicated as integrating the mEGFP tag at both alleles, although further genotyping indicated that ∼3.5% of cells integrated the mEGFP tag on only one allele. The donor plasmids essential for creating these cell lines can be obtained through Addgene (https://www.addgene.org/The_Allen_Institute_for_Cell_Science/). Comprehensive details regarding these cell lines are accessible via the Allen Cell Collection (https://www.allencell.org/), and cells can be acquired through Coriell (https://www.coriell.org).

#### Generation of CDH5 exon 3 deletion mutant hiPS cell line

AICS-0126 cl. 41 hiPS cells were additionally targeted with paired CRISPR/Cas9 ribonuclear protein complexes designed to delete exon 3 of gene model NM_001795 at genomic coordinates –66,386,774 and +66,387,161 respectively using sgRNAs (Synthego) targeting the genomic sequences 5’ TGACGTGAGCTGGGAATGGG 3’ and 5’ ATCAGCCACTTCACTGCCTG 3’ (hg38 human genome assembly). This approach replicated the CDH5 knockout strategy used previously in a murine model system,^108^ but resulted in an N-terminal truncation product of VE-cadherin being expressed (see *Characterization of Ex3Del cells)*. The anticipated 407 bp deletion of the exon was confirmed with PCR and electrophoresis in both transfected populations and monoclonal cells derived from the edited population. Sanger sequencing confirmed the presence of two deletion alleles in monoclonal culture. The resultant cell line was denoted as AICS-0177 cl. 26, referred to as Ex3Del, and is available upon request.

### hiPS cell culture

WTC-11 hiPS cells were maintained in mTeSR^™^1 (Cat # 85850, STEMCELL Technologies) supplemented with 1% penicillin/streptomycin (P/S) on 0.337 mg/mL growth factor-reduced Matrigel (Cat # 356231, Corning)-coated 10-cm dishes (Cat # 353003, Corning). Media was exchanged daily. When cells were approximately 80% confluent (every 3-4 days), media was removed from the dish and 10 mL of Dulbecco’s Phosphate Buffered Saline (DPBS), no calcium, no magnesium (Cat # 14190144, Gibco) was used to wash cells. The hiPS cells were then dissociated into single cells using 3 mL StemPro^™^ Accutase^™^ (Cat # A11105-01, Gibco) incubation for 4 minutes at 37 °C and 5% CO_2_. The dissociation reaction was stopped by adding DPBS to the lifted cell suspension, then spun down at 1000 rpm for 3 minutes in a 15 mL conical tube. The supernatant was discarded, and the cell pellet was resuspended in 10 mL of mTeSR^™^1 medium with 1% P/S and 10 μM Y-27632 (Cat # 72308, STEMCELL Technologies). Cells were counted using a Vi-Cell XR cell counter (Beckman Coulter) and 0.50 x 10^6^ – 1.00 x 10^6^ cells were seeded into a new Matrigel-coated 10-cm dish (56.7 cm^2^) containing 10 mL mTeSR^™^1 medium with 1% P/S and 10 μM Y-27632. A comprehensive cell hiPS cell culturing protocol is available at https://www.allencell.org/written-protocols.html.

### Endothelial cell differentiation and magnet-assisted cell sorting

The endothelial cell (EC) differentiation protocol was adapted from the methods described by Patsch *et al*. (2015)^109^ with modifications made to cell plating density and CHIR 99021 concentrations. First, hiPS cells were plated as single cells at 2.5 x 10^5^ – 1.00 x 10^6^ cells per well (9.6 cm^2^) of a 0.337 mg/mL Matrigel-coated 6-well plate (Cat # FB012927, Fisher Scientific) in mTeSR^™^1 supplemented medium with 1% P/S and 10 µM Y-27632. Initial cell density was adjusted regularly to maintain >10% yield from magnet-assisted cell sorting. After 24 hours (denoted as day 0 of the protocol), media was replaced with 3 mL/well of mesoderm induction media: 4.5-6 µM CHIR 99021 (Cat # 4423, Tocris; concentration adjusted on a per-lot basis) and 25 ng/mL Recombinant Human BMP-4 Protein (Cat # 314-BP, R&D Systems) in “N2B27 media”, which was composed of: 1X B-27^™^ supplement (Cat # 17504044, Gibco), 1X N-2 supplement (Cat # 17502048, Gibco), 1X GlutaMAX^™^ Supplement (Cat # 35050061, Gibco), 48 mL DMEM/F-12 (Cat # 11320033, Gibco), and 48 mL Neurobasal^™^ Medium (Cat # 21103049, Gibco). After 72 hours, this mesoderm induction media was exchanged with 3 mL/well endothelial induction media consisting of 200 ng/mL Human VEGF-165 Recombinant Protein (Cat # 100-20, PeproTech) and 2 µM Forskolin (Cat # F6886, Sigma-Aldrich) in StemPro^™^-34 serum-free medium (Cat # 10639011, Gibco) supplemented with 1X GlutaMAX (Cat # 35050061, Gibco) to drive cells toward endothelial specification. The cells were replenished with another 3 mL/well endothelial induction media on day 4. On day 5 of the differentiation protocol, cells were harvested as single cells using 1 mL/well StemPro^™^ Accutase^™^ and magnetically sorted for CD144^+^ expression with CD144 (VE-Cadherin) MicroBeads (Cat # 130-097-857, Miltenyi-Biotec) as described in the manufacturer’s protocol (CD144 (VE-Cadherin) MicroBeads human Data Sheet, Miltenyi-Biotec). Alternatively, AICS-0177 CDH5 exon 3 deletion cells and AICS-0126 sorting control cells were sorted using CD31 (PECAM-1) MicroBeads (Cat # 130-091-935, Miltenyi-Biotec). Briefly, “MACS buffer” was prepared with the addition of 5% MACS BSA Stock Solution (Cat # 130-091-376, Miltenyi-Biotec) into autoMACS^®^ Rinsing Solution (Cat # 130-091-222, Miltenyi-Biotec), then passed through a 0.2 µm filter to prevent clogging of the MS Columns (Cat # 130-042-201, Miltenyi-Biotec). The cell suspension, kept on ice, was diluted in MACS buffer and passed through a 40 µm cell strainer to remove large cell debris and clumps. The cells were centrifuged for 10 minutes at 300 x g, and the supernatant was discarded afterwards. The remaining cell pellet was resuspended in 20 µL of CD144 or CD31 MicroBeads and 80 µL of MACS buffer for every 1.00 x 10^7^ cells. After incubating the cell suspension at 4 °C for 15 minutes, 2 mL of MACS buffer was added prior to centrifugation at 300 x g for 10 minutes. The supernatant was discarded carefully, and the pellet was resuspended in 0.5 mL of MACS buffer for every 1.00 x 10^8^ cells. MS Columns were mounted on an OctoMACS^®^ Separator (Cat # 130-042-109) and MACS MultiStand (Cat # 130-042-303, Miltenyi-Biotec), then pre-washed with MACS buffer prior to addition of the cell suspension. The suspension was then washed through each column completely 3 times with MACS buffer, allowing the CD144^-^ or CD31^-^ cells to be discarded whilst the CD144^+^ or CD31^+^ cells remained bound to the magnetic column. The columns were then removed from the magnetic separator, and the plunger was used to dispense CD144^+^ or CD31^+^ cells from the columns into a 15 mL conical tube. The resulting, purified hiPS cell-derived ECs were manually counted with a hemocytometer and either cryogenically banked at 3.00 x 10^5^ cells per 300 µL of CryoStor^®^ CS10 (Cat # 07930, STEMCELL Technologies) or replated for endothelial marker validation via flow cytometry.

#### Endothelial cell marker validation

Flow cytometry for CD31 and CD144 expression was performed to confirm endothelial cell identity. ECs were replated at 1.00 x 10^5^ cells per well (9.6 cm^2^) onto a 5 µg/mL fibronectin (Cat # 33016015, Gibco) coated 6-well plate. When cells were at least 80% confluent (approximately 3-5 days), ECs were dissociated into a single-cell suspension using 1 mL of StemPro^™^ Accutase^™^ and centrifugated in a 15 mL conical tube at 300 x g for 5 minutes in DPBS. The cell pellet was resuspended with 4% paraformaldehyde (Cat # 15710, Electron Microscopy Sciences) in DPBS and incubated at room temperature for 10 minutes for fixation prior to spinning down at 90 x g for 3 minutes. The supernatant was discarded, and the pellet was resuspended in DPBS for another round of centrifugation at 200 x g for 3 minutes. The supernatant was aspirated off, and the cell pellet was resuspended in 5% Fetal Bovine Serum (FBS) (Cat # A1895604, Gibco) in DPBS. The resultant fixed samples were stored at 4 °C prior to running flow cytometry.

Fixed cells were aliquoted at 2.00 x 10^5^ – 2.50 x 10^5^ cells per well of a U-bottom 96-well plate (Cat # 353077, Falcon). 100 µL of 2% FBS in DPBS was added to each well, and the plate was centrifuged at 500 x g for 4 minutes. The supernatant was discarded, then 200 µL of 2% FBS in DPBS was used to resuspend cells, with another centrifugation at 500 x g for 4 minutes. The supernatant was discarded again. Subsequently, cell pellets were resuspended in 2% FBS in DPBS with either 1.3% added BD Horizon^™^ BV786 Mouse Anti-Human CD144 antibody (Cat # 565672, BD Biosciences), BD Horizon^™^ BV786 Mouse IgG1, k Isotype Control (Cat # 563330, BD Biosciences), CD31 (PECAM-1) Monoclonal Antibody APC, (Cat # 17-0319-42, Invitrogen), or Mouse IgG1 kappa Isotype Control APC (Cat # 17-4714-82, Invitrogen). Cells in their antibody cocktails were allowed to stain for 30 minutes at room temperature in the dark prior to adding another 100 µL of 2% FBS in DPBS per well and centrifugating at 500 x g for 4 minutes. The supernatant was discarded, and stained samples were then resuspended in 200 µL of 2% FBS in DPBS containing 20 µg/mL DAPI (Cat # D3571, Invitrogen). Fluorescence intensities of samples in the U-bottom plate were measured using a CytoFLEX S Flow Cytometer (Beckman Coulter). Resulting data was analyzed using FlowJo (BD Biosciences, v10) software to determine the percentage of populations that were double positive for CD31 and CD144. DAPI, forward scatter, and side scatter measurements were first used to gate for intact, single cells. Isogenic controls for each sample were used to gate for APC (CD31) and BV786 (CD144). Reported CD31/CD144 double positive populations were those that measured above the isogenic control thresholds.

### Western Blot

For preparation of the blots staining the N-terminus of VE-cadherin and GFP, whole cell lysates were extracted directly from AICS-0000, AICS-0126, and AICS-0177 hiPS cell-derived EC monolayers using M-PER buffer (Cat # 78501, Thermo Scientific) for 15 minutes on ice. 1X LI-COR SDS sample buffer (LI-COR 4X Protein Sample Loading Buffer, Cat #928-40004, LICORbio) and 10X BOLT Sample Reducing Agent (Cat #B0009, Invitrogen) was added to 70 µL of cell lysate, and the samples were boiled for 10 minutes on a 70 °C heat block.

Samples were separated on 7% Tris-Acetate gels (Cat # EA03555BOX, ThermoFisher). After transfer onto nitrocellulose membranes, a total protein stain was applied and used as a loading control (LI-COR Revert Total Protein Stain, Cat # 926-11011, LICORbio; not shown). After imaging the total protein stain, blots were incubated for 30 minutes at room temperature with LI-COR Intercept TBS Blocking Buffer (IBB, Cat # 927-85001, LICORbio). Blots were then incubated in IBB + 0.2% Tween20 solution containing primary antibodies to detect the N-terminus of VE-Cadherin (1:500; mouse VE-cadherin monoclonal antibody; Cat # MA5-46904, Thermo Fisher), and GFP (1:2000; goat GFP antibody, Cat # 600-101-215, Rockland) at 4 °C overnight. Next, blots were incubated in IBB + 0.2% Tween containing respective LI-COR IRDye 800CW secondary antibodies (IRDYE® 800CW Goat anti-mouse IGG secondary antibody, Cat # 926-32210; IRDYE® 800CW Goat anti-mouse IGG secondary antibody, Cat # 926-32214) at 1:5000 for 1 hr at room temperature. Resulting blots are shown in Figure S11.

### Characterization of Ex3Del cells

The CDH5 editing strategy followed to generate the Ex3Del cells replicated the VE-cadherin knockout strategy used previously in a murine model system.^108^ In hiPSC-ECs, this exon deletion produced cells that expressed an N-terminal truncated form of VE-cadherin. Western blots indicated that Ex3Del hiPSC-ECs expressed N-terminal truncated forms of VE-cadherin with and without an mEGFP tag (Figure S11B-C). The size of the detected protein products indicate that the EC1 and EC2 domains of VE-cadherin are absent (Figure S11B-D). We confirmed the presence of a monoallelic mEGFP tag on VE-cadherin in Ex3Del cells using a custom-designed ddPCR assay.^107^ A summary of modifications to CDH5 in Ex3Del cells is depicted in Figure S11E.

### Endothelial cell culture in glass bottom µ-Slide

ECs were thawed at 1.00 x 10^5^ cells per well (9.6 cm^2^) onto a 5 µg/mL fibronectin coated 6-well plate. Cells were replenished with 2 mL of “EC media” composed of: StemPro^™^-34 serum-free medium (Cat # 10639011, Gibco) supplemented with 1X GlutaMAX (Cat # 35050061, Gibco) and 50 ng/mL Human VEGF-165 Recombinant Protein (Cat # 100-20, Gibco, PeproTech), every other day for 5 days. Additional EC media was then pre-equilibrated overnight in a loosely-capped 15- or 50-mL conical tube at 5% CO_2_ and 37 °C in the absence of VEGF, which was subsequently added to media at 50 ng/mL on the day of replating to minimize growth factor degradation. Pre-equilibration played a key role in minimizing bubble formation within downstream tubing and slide components of the ibidi-based flow assay, which prevented EC attachment to or directly detached the ECs from the fibronectin-coated glass surface. When the ECs reached 80-100% confluency in each well (approximately 5 days after thaw), cells were dissociated into a single-cell suspension using 1 mL of StemPro^™^ Accutase^™^ and prepared at 4.00 x 10^4^ – 6.00 x 10^4^ cells in every 162.5 µL (channel volume) of pre-equilibrated EC media. Pre-equilibrated µ-Slides (0.6 #1.5H µ-Slide I Luer Glass Bottom channel slides, Cat # 80187, ibidi) were coated with 162.5 µL of 42.67 µg/mL fibronectin for 1 hour, then the excess coating was displaced by washing through the slide 3 times with 162.5 µL of the prepared cell suspension to achieve an even distribution of cells across the slide. ECs were allowed to settle on the slide for 15 minutes in an incubator prior to adding 60 µL of pre-equilibrated EC media to both the left and right channel ports. Cells on the µ-Slides were incubated overnight at 5% CO_2_ and 37 °C to form a 90-100% confluent monolayer prior to applying shear stress.

### Preparation of fluidic unit and pump (ibidi Pump System)

A µ-Slide with a 90-100% confluent EC monolayer (prepared as described in *Endothelial cell culture in glass bottom µ-Slide)* was attached to a fluidic unit (Cat # 10903, ibidi) with yellow-green Perfusion Set tubing (Cat # 10964, ibidi) and syringe reservoirs (Cat # 10964, ibidi). The reservoirs and tubing were filled with 16 mL of overnight pre-equilibrated EC media with 50 ng/mL VEGF and 1X of 100X P/S added freshly such that there was approximately 6 mL of volume in each syringe reservoir.

The pump (Cat # 10902, ibidi) was used to maintain unidirectional laminar flow across ECs plated in µ-Slides. Air intake tubing was placed in the incubator such that warm 5% CO_2_ air could pass through a drying bottle filled with orange Silica gel beads (Cat # 13767-1KG-R, Sigma-Aldrich) prior to entering the pump. The pump and drying bottle were kept outside of the incubator. The air pressure output tube was placed in the incubated environment. The prepared fluidic unit was attached to the pump using both the air pressure output tube and electrical connector, then placed in the incubator.

### Specification and validation of shear stress

The ibidi PumpControl software (v1.6.1) was used to regulate pressure through the ibidi Pump System to achieve desired shear stresses. For the µ-Slides, the shear stress (τ; dyn/cm^2^) was calculated using the equation:

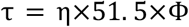

where η is the dynamical viscosity (dyn • s/cm^2^), 51.5 is the slide-specific rectangular geometry factor provided by the manufacturer, and Φ is the flow rate (mL/minute). The desired shear stress was set in the PumpControl software, which automatically calculated the flow rate according to Poiseuille’s law for laminar flow:

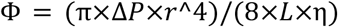

where ΔP is the pressure differential applied, L and r are the length and radius of the connecting tubing, respectively, and η is the dynamical viscosity of the fluid. The yellow-green Perfusion Set tubing used in this setup was 50 cm in total length with an inner radius of 0.8 mm. The dynamical viscosity of the EC medium containing 10% serum at 37 °C was assumed to be 0.0072 dyn • s/cm^2^, the standard media viscosity constant provided by ibidi.

To verify the actual shear stress applied, the flow rate was manually measured by recording the time required to displace 1 mL of medium from the graduated syringe reservoir. The average of three manual flow rate measurements (mL/min) was used to calculate the actual shear stress applied to the cells. Verified shear stress values were rounded to the nearest integer to reflect the level of confidence in the measurement.

### Preparation of shear stress experiments for microscope image acquisition

The pump and fluidic unit were prepared as described in *Preparation of fluidic unit and pump system* with the following additional specifications adapted from ibidi *Application Note 14: Live Cell Imaging Setups Under Flow*. The ibidi Pump System air intake tubing was positioned close to the gas supply port of the microscope stage top incubator (H201-Frame, Okolab) such that maximal 5% CO_2_ air could pass through a drying bottle filled with orange Silica beads with moisture indicator of free heavy metals, (Cat # 13767-1KG-R, Sigma-Aldrich) prior to entering the pump. The air pressure output tube from the pump was placed within the microscope environmental chamber (Okolab). To “prime” the pump with warm air, the ibidi PumpControl software (v1.6.1) was used to supply 50 mbar pressure through the pump system for at least 1 hour before connecting the prepared fluidic unit. This pre-equilibration step was critical in preventing the formation of bubbles in the media during shear stress experiments on the microscope. After pre-equilibration, the prepared Fluidic Unit was attached to the pump using both the air pressure output tube and electrical connector, then placed in the outer environmental chamber. The pump and drying bottle were kept outside of the microscope environmental chamber.

The bottom of the glass channel slide containing ECs was cleaned with 70% ethanol and dried with a Kim wipe prior to being mounted on the microscope stage within the stage-top incubator. As much of the Perfusion Set tubing as possible (approximately 60%) was placed within the lidded stage-top incubator, which was then wrapped in Parafilm to improve stability of environmental conditions and temperature of media within the tubing. The PumpControl software was used to initiate the various shear stress conditions as described in *Specification and validation of shear stress*. The glass channel slide was allowed to equilibrate on the stage with shear stress treatment for at least 1 hour before imaging.

### Preparation of shear stress experiments for incubator control

To validate comparable cell health and alignment response for imaged µ-Slides, a replicate µ-Slide on a separate Fluidic Unit was maintained in a Heracell^™^ VIOS 160i CO_2_ incubator (Cat # 50145502, Thermo Scientific) under the same shear stress conditions simultaneously. These pairwise experiments were necessary to run in the case that we observed excessive cell death on the imaging slide from phototoxicity or issues with the physical set-up that deemed additional troubleshooting with the imaging set-up was needed. The incubator was set to 37 °C and 5% CO_2_. The pump and fluidic unit were prepared as described in *Preparation of fluidic unit and pump system*. The PumpControl software was used to initiate the various shear stress conditions as described in *Specification and validation of shear stress*. The incubator control was identical to the image acquisition set-up (described above, *Preparation of shear stress experiment for image acquisition*) except for the Fluidic Unit being housed in the same incubator as the µ-Slide, negating the need for the 1-hour priming of the pump. Additionally, incubator control set-up was absent of any laser exposure throughout the shear stress experiment.

### Preparation of shear stress experiments for fixation

ECs fixed at specific timepoints underwent shear stress application as described above in *Preparation of shear stress experiment for incubator control* with the addition of using Serial Connectors (Cat # 10830, ibidi) between up to 3 µ-Slides in series for any given Fluidic Unit. Flow was paused to disconnect each µ-Slide for fixation outside of the incubator, then tubing was reconnected in series to commence shear stress treatment.

### Fixation

All solutions were washed through µ-Slides 3 times by pipette to completely exchange reagents while allowing cells to remain attached and bathed in fluid within the channel area, unless otherwise specified. The method of washing is as described by the manufacturer in the document *Instructions µ-Slide I Luer Glass Bottom* for the µ-Slides (Cat # 80187, ibidi). Endothelial cells were washed with 4% paraformaldehyde in DPBS, taking care not to form bubbles in the cell channel area, and incubated at room temperature for 10 minutes. The slides were then washed with 0.02% sodium azide (Cat # BDH7465-2, VWR) in DPBS for storage up to 1 month at 4 °C.

To permeabilize the ECs, 0.5% Triton^®^ X-100 (Cat # A16046.AE, Thermo Scientific) was washed through the slide and incubated at room temperature for 10 minutes. 1% BSA (Cat # 15260-037, Gibco) in 1X BD Perm/Wash^™^ (Cat # 51-2091KZ, BD Biosciences) was washed through the slide, then incubated at room temperature for 30 minutes for the blocking reaction. Fixed and permeabilized cells were either imaged after this step or stained with antibodies for immunofluorescence.

### Immunofluorescence staining

Cells were fixed and permeabilized as described above (*Fixation*). Primary antibodies were diluted as follows in 1X BD Perm/Wash^™^, then washed 3 times with 162.5 µL (0.6 channel volume) of Anti-Sox17 Clone OTI3B10 Mouse Monoclonal Antibody (1:100; Cat # TA500044, OriGene) and Anti-NR2F2 rabbit monoclonal antibody (1:100; Cat # EPR18443, abcam]. 1X BD Perm/Wash^™^ was washed through for no primary antibody control samples. Slides were then stored overnight at 4 °C. After 24 hours, the slides were washed with DPBS. Secondar antibodies were washed 3 times through the slides at 162.5 µL and used at 1:500 in 1X BD Perm/Wash^™^ as follows: Goat anti-Mouse IgG (H+L) Alexa Fluor^™^ Plus 555 (Cat # 32727, Invitrogen), Goat anti-Rabbit IgG (H+L) Alexa Fluor^™^ Plus 647 (Cat # A32733, Invitrogen). Nuclear Violet^™^ LCS1 (Cat # 17543, AAT BioQuest; “NucViolet”) was used at 1:1500 in the secondary antibody mixture. The hiPS cell-derived ECs were incubated with the secondary antibody mixture at room temperature for 2 hours, prior to washing and storing with 0.02% sodium azide in DPBS. Slides were imaged within 2 weeks of fixing and staining.

### Image acquisition

#### Spinning-disk confocal microscopy

Imaging was performed on a Zeiss AxioObserver 7 inverted microscope coupled to a Yokogawa CSU-W1 confocal scan head using a 50 µm spinning disk (system was integrated by 3i - Intelligent Imaging Innovations, Inc.). The microscope was outfitted with an environmental chamber (Okolab) and a stage top incubator (H201-Frame, Okolab) to maintain the cells at 37 °C and provide humidified air with 5% CO_2_ during imaging. A 0.83X magnification changer was used to enlarge the field of view. A 20X Plan-Apochromat objective (numerical aperture 0.8, Zeiss) was used to perform timelapse imaging. A 10X Plan-Apochromat (NA 0.45, Zeiss) was used for position selection. Image acquisition routines were designed in Slidebook software (Intelligent Imaging Innovations, versions from 2023 to 2025). A custom built beam expansion device (“mesa” by 3i) was placed between the laser fiber and Yokogawa W1 scanhead to provide a uniform illumination profile across the field of view. Optical control images were acquired regularly to monitor field uniformity and camera alignment. A 488 nm laser (LuxX Diode laser series) was used to provide excitation at a nominal laser power of 2 mW, optimized for long live-cell timelapse imaging. Laser power was measured monthly at the 10X objective to calibrate power settings in Slidebook. Across timelapse acquisitions, the measured 488 nm laser power averaged 2.08 ± 0.12 mW. An LED light source (Lambda TLED+, Sutter Instruments) with a peak emission of 740 nm was used to provide illumination for transmitted light images. mEGFP fluorescence and transmitted light signals were split by a 660LP dichroic beam splitter and filtered through a 525/50 emission filter and a 706/95 emission filter, respectively, and were simultaneously detected by two Prime BSI sCMOS cameras (Photometrics) with an exposure time of 100 ms or 101 ms, with the exposure time chosen to minimize stripe artifacts due to disk-camera synchronization. The intensity histogram of brightfield images was adjusted to peak at around ∼14,000 in grayscale value. The nominal Z-step size for 3D image stacks was 0.53 µm for the 20X objective and a total of 25 Z-steps was acquired. A piezo Z-stage (Mad-City Labs) was used for fast acquisition of Z-stacks.

#### Imaging position selection

Imaging positions were selected near the central region of the µ-Slide where the shear stress is noted as homogenous in the ibidi *Application Note 11: Shear Stress and Shear Rates for ibidi µ-Slides Based on Numerical Calculations* (Figure S1B). The port on the left side of the slide was used as a reference point, and the imaging field was positioned approximately one-quarter of the slide length toward the right. One region per experiment was chosen that contained a confluent, uniform endothelial monolayer without visible gaps or patches across the length of 6 tiled fields of view, arranged sequentially along the axis of fluid flow. Positions were preferentially selected toward the left side of the slide, as cells exposed to 6 dyn/cm^2^ shear stress migrate upstream, resulting in increased patchiness at the downstream end over time. Areas containing visible dust particles or cellular debris were avoided.

#### Acquisition of 20X 48-hour 3D timelapse data

hiPS-EC monolayers were cultured and imaged as described above (see *Endothelial cell culture in glass bottom µ-Slide, Preparation of shear stress experiment for image acquisition,Spinning-disk confocal microscopy, Imaging position selection*). Timelapses were acquired at shear stress conditions ranging from 0-24 dyn/cm^2^. To capture cells along the long axis of the flow chamber, image montages were generated by tiling six horizontally arranged fields of view (FOVs; each 1744 pixels x 1712 pixels, XY pixel size = 0.382 µm) with 10% overlap. The SlideBook Focus Surface tool was used to correct sample tilt and maintain consistent focus across the montage. Timelapses were acquired for 48 hours at 5-minute intervals. If substantial cell crowding occurred, imaging was terminated early (typically ∼45 hours; Figure S1F). When shear stress is applied, fresh media is perfused through the slide. The shear stress is applied ∼1 hour before the start of imaging to allow the slide to equilibrate to incubator conditions before imaging. To extend the variation in cell morphology seen by the DiffAE model we collected data of cells exposed to shear stress that changed magnitude during timelapse acquisition, and cells cultured without fluid flow. To change shear stress during timelapse acquisition, flow was stopped between imaging timepoints. Media was slowly rebalanced by toggling 30 mbar of pressure on and off to match the fluid height in each syringe to the 6 mL gradation. Subsequently, the desired new shear stress was initiated before the start of the next imaging timepoint and actual shear stress was re-measured as described in *Specification and validation of shear stress*. For timelapses with no shear stress applied, fluidic units were not attached to slides. To maintain cell health, under the no–shear stress condition, media was replenished manually every 3 hours between imaging timepoints during business hours over the 48-hour timelapse. At each replenishment, 162.5 µL of pre-equilibrated EC media was washed through the slide three times, followed by the addition of 60 µL of fresh media to both the left and right ports. To verify that cell morphology, alignment, and overall cell health in the imaged µ-Slides were comparable to non-imaged controls, a replicate µ-Slide prepared on the same day and subjected to the same shear stress conditions was maintained in a standard cell culture incubator (see *Preparation of shear stress experiment for incubator control*). Timelapse images of hiPS cell-derived ECs were subjected to manual quality control checks. Timelapses were excluded if cells exhibited poor health as shown by poor attachment to the glass substrate, significant cell death above normal levels, or extremely low or high starting density. Resulting in a total of 32 timelapse replicates with 28 timelapses of AICS-0126 at shear stress conditions ranging from 0-24 dyn/cm^2^ and four timelapses of AICS-0177 at 6 dyn/cm^2^ acquired for the data set in this study (Table S1-5).

#### Acquisition of 20X immunofluorescence images

A total of 4 fixed hiPS cell-derived EC monolayers were prepared as described in *Endothelial cell culture in glass bottom µ-Slide, Fixation*, and *Immunofluorescence staining*. Shear stress experiments were performed as described in *Preparation of shear stress experiment for fixation*. Imaging was performed as described in *Spinning-disk confocal microscopy*, with laser powers optimized for each fluorescent channel: 0.8 mW for the NucViolet nuclear stain (405 nm), 11 mW for Anti-Sox17 (561 nm), and Anti-NR2F2 (640 nm) with imaging positions selected as detailed in *Imaging position selection*. A total of four µ-Slides of AICS-0126 subjected to shear stress levels ranging from 0–24 dyn/cm^2^ were imaged, with 6–50 single-FOV positions selected per µ-Slide.

### Image processing of 3D timelapse data for analysis

#### Single timepoint filter

To identify and exclude transient imaging artifacts from downstream analysis, including rapid changes in illumination, striping artifacts, and bubbles passing through the field of view, we implemented automated single time point quality control filters for both the brightfield and fluorescent channels. To detect Z-slices with brightfield artifacts, the average pixel intensity of each bright-field Z-slice was measured. These values were flattened into a sequence across Z-slices and timepoints, where each timepoint consists of 25 Z-slices (Figure S12A). A rolling median was computed across this flattened index using a window of 100 Z-slices (equivalent to 4 timepoints; Figure S12A). A timepoint was considered an outlier if it contained one or more Z-slices with a mean intensity greater than 1% from the rolling median (bright BF outlier) and less than 0.04% from the rolling median (dark BF outlier; Figure S12A). Bright outliers capture images where the transmitted light came on more intensely than programmed. Dark outliers capture images containing striping artifacts from variation in spinning disk speed, bubbles passing through the field of view (FOV), and when the transmitted light sporadically did not come on. This automated detection method detected brightfield artifacts in 1.4% of all 96420 timepoints. Identified timepoints were programmatically annotated and excluded from model training and downstream analysis.

A similar method was applied to the mEGFP fluorescence channel. While mEGFP fluorescence image artifacts were more rare than brightfield ones, their magnitude was such that they could be detected on a per timepoint basis. Therefore, to detect timepoints with artifacts in the mEGFP fluorescence channel, the mean pixel intensity was measured over the Z-stack for each timepoint (Figure S12B). A rolling median of 12 timepoints was calculated across the mean intensity per Z-stack over time (Figure S12B). Dark outliers and bright outliers were identified by timepoints deviating from the rolling median by more than ±1% (Figure S12B). This automated detection method detected fluorescence artifacts in 0.002% of all 96420 timepoints. Identified timepoints were programmatically annotated and excluded from model training and downstream analysis.

#### Position filter

If dust or debris was present on the bottom of the coverslip glass, all images from that position would contain bright-field artifacts from light scattered by the particles. FOVs were visually inspected and if dust or debris on the glass was detected, the position was annotated and excluded from model training and downstream analysis.

#### Unfed timepoint filter

In the no-shear stress condition, media is replenished every 3 hours only during business hours over the 48-hour timelapse (see *Acquisition of 20X 48-hour 3D timelapse movies*). Accordingly, any timepoints occurring more than 3 hours after a media change (i.e. acquired overnight), were annotated as unfed and excluded from model training and downstream analysis.

#### Determining the in-focus Z-plane

To find the in-focus plane at a given timepoint, the standard deviation of pixel intensities for each brightfield Z-slice was measured. The in-focus plane within a Z-stack was defined as the Z-slice with the minimum standard deviation (Figure S12C). To reduce sensitivity to transient brightfield artifacts, including bubbles and debris, and because the in-focus plane was highly consistent over time for a given position, the in-focus plane for each timelapse position was defined as the mean in-focus plane across all timepoints rather than on a per-timepoint basis (Figure S12D). The in-focus plane was found to consistently detect a slice close to the glass, as opposed to the center of cells.

#### Z-slice selection filter

To ensure that blur captured in the brightfield image and projected in the standard deviation Z-projection reflects biological differences in cell height and morphology rather than technical differences in the position of cells within the Z-stack, we generated consistent subsets of the full Z-stacks (“sub-Z-stacks”) relative to the identified in-focus plane near the glass surface. Across all replicates, at least 11 Z-slices were available above the in-focus plane (Figure S12E); therefore, a consistent upper offset of +11 slices was used (Figure S12F). To provide a comparable amount of out-of-focus information above and below the cells, a lower offset of -4 slices was used across all replicates (Figure S12F).

#### Visual steady state filter

Each FOV from all timelapse replicates was manually annotated using VE-cadherin-mEGFP maximum intensity Z-projections to identify the timepoint at which approximately 75% of imaged area reached steady state alignment and behavior. Steady state for each FOV was first determined by observing alignment and migratory behavior after 24 hours of culture under the specified shear stress condition. The timepoint when approximately 75% of the FOV by area first exhibited that alignment and behavior was then annotated as the beginning of steady state. In replicates where the shear stress condition was changed during the timelapse experiment, this process was repeated following the shear stress change. All timepoints prior to the onset of this visual steady state in each FOV were annotated and excluded from downstream analyses.

#### Cell crowding filter

Each FOV from all timelapse replicates was manually annotated using both the brightfield standard deviation Z-projection and the VE-cadherin-mEGFP maximum intensity Z-projection to demarcate when approximately 30% of the imaged area showed cell crowding. Cell crowding was characterized by loss of cell elongation and alignment and disruption of migratory behavior of cells, as well as an increase in high-intensity patches of VE-cadherin-mEGFP signal and qualitative difficulty determining cell-cell boundaries via VE-cadherin-mEGFP signal (Figure S12G). All timepoints occurring after the onset of cell crowding in each FOV were annotated and excluded from both model training and downstream analysis. If cell crowding was present, its onset defined the end of the steady-state regime.

### Creation of image datasets

#### DiffAE model training FOV dataset

Eight of the 32 timelapse movie replicates acquired as described in *Acquisition of 20X 48-hour 3D timelapse movies* were selected for downstream diffusion autoencoder (DiffAE) model training (Table S2). One representative replicate was chosen for each of the following shear stress conditions: 0, 6, 9, 12, 15, 21, 6–20, and 20–6 dyn/cm^2^. Movies were selected to capture biological variation in initial cell density, alignment relative to flow (parallel, perpendicular, or mixed), instances of cell death and division, and degrees of cell crowding. Z-stacks subsets were created consistently around the in-focus brightfield plane (see *Z-slice selection filter*), and FOVs were filtered according to the criteria in *Single timepoint, Unfed timepoint, Position*, and *Cell crowding filter* resulting in 46 FOVs in the quality-controlled “DiffAE model training FOV dataset” (Table S2).

#### Shear stress FOV dataset

23 of the 32 timelapses movie replicates acquired as described in *Acquisition of 20X 48-hour 3D timelapse movies* were selected for downstream analysis comparing shear stress conditions (Table S1). The dataset included AICS-0126 cells imaged under five shear stress conditions: 6 dyn/cm^2^ (n = 3), 9 dyn/cm^2^ (n = 2), 12 dyn/cm^2^ (n = 7), 15 dyn/cm^2^ (n = 9), and 21 dyn/cm^2^ (n = 2). Z-stacks subsets were created consistently around the in-focus brightfield plane (see *Z-slice selection filter*), and FOVs were filtered according to the criteria described in the *Single timepoint, Position*, and *Cell crowding* filters, resulting 131 FOVs in the quality-controlled “Shear stress FOV dataset” (Table S1).

#### VE-cadherin Exon3Del perturbation FOV dataset

Five of the 32 timelapse movies acquired as described in *Acquisition of 20X 48-hour 3D timelapse movies* were collected as a perturbation for downstream analysis (Table S3). Four timelapse movies of the AICS-0177 VE-cadherin Exon3Del cell line and 1 timelapse movie of AICS-0126 sorted for CD-31 expression were collected at 6 dyn/cm^2^. Z-stacks were centered on the in-focus plane (see Z-slice selection filter), and FOVs were filtered according to the criteria in *Single timepoint, Position, and Cell crowding* filters. For the AICS-0177 replicates, each FOV was further manually annotated for cell delamination. Unlike AICS-0126, which exhibited cell crowding at later timepoints, these replicates instead showed loss of monolayer integrity due to apparent cell detachment from the substrate. The onset of delamination was defined as the timepoint at which approximately 30% of each FOV exhibited detachment, and all subsequent timepoints were excluded from downstream analysis. This resulted in 30 FOVs in the quality-controlled “VE-cadherin Exon3Del perturbation FOV dataset” (Table S3).

#### Label-free nuclear segmentation training FOV dataset

Three of the nine datasets, corresponding to 159 FOVs, acquired as described in *Acquisition of 20X immunofluorescence* were used to generate ground-truth nuclear segmentations, utilizing the NucViolet-stained nuclei as targets and the corresponding brightfield images as input (Table S4).

### Generating label-free nuclear segmentations

To enable label-free detection of nuclei throughout timelapse experiments, we trained a model to generate nuclear segmentations from the brightfield channel. Using the images in the *Label-free nuclear segmentation training* FOV dataset (Table S5), ground-truth segmentations of nuclei were generated from maximum intensity Z-projections of the NucViolet stained channel using the default Cellpose “nuclei” model (Cellpose version 3.0.1.1).^110^ All resulting segmentations were visually inspected for correctness. Next, brightfield standard deviation Z-projections and their corresponding Nuc-Violet-based ground-truth segmentations were split into an 80:20 train:test set and passed to the default Cellpose “nuclei” model to finetune it for label-free nuclei detection using data normalization, stochastic gradient descent, a learning rate of 0.1, a weight decay of 0.001, and 300 training epochs. Inference was then performed on all brightfield images in the *DiffAE model training* and *Shear stress FOV datasets*. To evaluate performance, all timelapses were visually inspected every 48 timepoints to confirm the absence of false nuclear predictions and to estimate the fraction of nuclei that were successfully predicted. These predicted nuclear segmentations were used to refine cell segmentations and for other downstream analyses.

### Generating cell segmentations via VE-cadherin

To generate cell segmentations from VE-cadherin-mEGFP signal, a hysteresis threshold with a high threshold value of 80% of the image intensity and a low threshold value of 66% of the image intensity was applied to the VE-cadherin-mEGFP maximum intensity Z-projection. The inverse of the thresholded image was used to generate a distance transform for peak detection and seed generation. These seeds were used to perform an initial watershed segmentation. Adjacent watershed regions were merged when VE-cadherin-mEGFP intensity along their shared boundaries was low, indicating that they belonged to the same cell. Nuclei segmentations (described above) were then used to identify multinucleated cell regions. For regions containing a single nucleus, a morphological skeleton was generated to preserve cell shape. The skeletons and nuclear segmentations were subsequently used as seeds for a second watershed segmentation to refine the final cell boundaries (Figure S2A). All timelapse images from the *DiffAE model training* and *Shear stress FOV datasets* were segmented (Table S5). Segmentations were visually inspected every 48 timepoints, confirming that ∼70% of each FOV was accurately segmented.

### Single cell measured features

For each segmented cell, several morphological and intensity features were computed using scikit-image’s region property measurements.^111^ These included “cell centroid coordinates,” defined as the geometric center of each segmentation; “cell orientation,” defined as the angle of the major axis of a fitted ellipse relative to the horizontal axis; “cell aspect ratio,” defined as the ratio of the major to minor axis lengths of the fitted ellipse; and “cell area,” defined as the 2D area of the segmentation. Intensity features included “cell mean fluorescence intensity,” computed as the mean intensity within each segmentation, and “cell edge fluorescence intensity,” computed as the mean intensity within a 2-pixel-wide perimeter of the segmentation.

To further characterize cellular responses to shear stress, we computed additional orientation-based features. The “cell–nucleus angle relative to migration” quantifies nuclear orientation within each cell relative to its direction of migration, defined as the angle between the nucleus–centroid vector and the cell segmentation centroid displacement vector, mapped from ±180° to 0° (with 0° corresponding to the direction of migration).

### Single-cell tracking

Single-cell trajectories were generated from tracking cell segmentations over time. To tolerate transient segmentation errors, cells were linked across short segmentation gaps by matching to the first subsequent segmentation with >50% pixel overlap within the next four timepoints. Matching was done reciprocally (forward and backward) to ensure that small regions were not incorrectly assigned as fully overlapping with larger regions.

To obtain reliable single-cell trajectories for analysis, a series of quality-control filters were applied to the single-cell tracks. Trajectories were required to have a minimum duration of at least 24 timepoints. Timepoints where segmentations touched the image boundaries were excluded. Mis-segmentations were detected automatically based on a change in area. To do so, cell area over time was smoothed using a one-dimensional Gaussian filter (σ = 2), and normalized area was computed as the ratio of raw area to smoothed area. Timepoints with >10% deviation in normalized area were removed. After filtering, trajectories had to retain at least 20 valid timepoints; those that did not were excluded. This procedure removes transient segmentation errors while preserving most of each trajectory, typically yielding ≥100 trajectories per FOV, or ∼600 per timelapse (Table S5).

### Single cell measured dynamic features

The “cell migration angle” captures the direction of cell migration relative to flow. It was calculated as the angle of the displacement vector of the cell centroid between consecutive segmented timepoints relative to the x-axis. If a segmentation was missing, the next available timepoint was used (maximum gap of three timepoints, ΔT ≤ 3), and the displacement was divided by the number of timepoints spanned to obtain an average per-timepoint movement in microns. The average was not interpolated for missing segmentations. Angles were mapped from ±180° to 0° (with 0° representing the direction of flow along the X-axis).

The “cell-nucleus angle relative to migration” quantifies the orientation of the nucleus relative to the direction of cell movement. It was computed as the angular difference between the nuclear orientation and the migration angle, such that 0° indicates alignment between the nucleus and the direction of migration.

### Diffusion autoencoder for unsupervised representation learning

#### Model architecture

To learn unsupervised representations of the brightfield images, we used a diffusion autoencoder (DiffAE), which is a type of image-to-image conditioned diffusion model.^61^ Unlike a traditional autoencoder where the input and output images are identical, we constructed a model in an image-to-image translation framework where a different paired image is used as the conditioning object. Specifically, we conditioned on the brightfield image while generating and computing the loss on the paired VE-cadherin image, rather than using the VE-cadherin image as both the conditioning input and reconstruction target.

For the semantic encoder, we used a tiny ConvNext^112^ with a latent dimension of 512, modifying the first layer of the ConvNext to accept a 1-channel grayscale brightfield input. We used the MONAI^113^ implementation of the conditional DDIM^114^ U-Net with [32, 64, 128, 256] feature channels per resolution level. For the cross-attention conditioning mechanism, we again used the MONAI implementation for conditional diffusion models,^115^ incorporating 64 channels per head at the second, third, and fourth levels of the U-Net. We additionally applied adaptive group normalization for temporal conditioning and used combined linear projections to efficiently fuse the query, key and value computations. During training, we sampled timesteps for the diffusion process from 0 to 1000 to match the implementation of Song *et al*. (2022).^114^ Similarly, for image reconstruction, we used the 50 timesteps for evaluation.^114^ To accelerate training, we used the Min-SNR weighting strategy from Hang *et al*. (2024)^116^ with γ = 5.0.

#### Data preprocessing

The model was trained on paired brightfield/VE-cadherin images from the *DiffAE model training FOV dataset* (Table S2). Images were down sampled to half their original XY resolution (from 1744 x 1712 pixels to 872 × 856 pixels). Next, consistent sub-Z-stacks were created as described (*Z-slice selection*). Each bright-field FOV sub-Z-stack was then converted into a standard deviation Z-projection, log normalized, clipped to the 1st and 98^th^ percentiles, and Z-score normalized (Figure S4A). Log normalization prevented bright debris from dominating the intensity histogram and increased the contrast of cellular features, reducing the skewness of the intensity distribution and improving representation of the cellular signal. Each mEGFP-VE-cadherin fluorescence FOV sub-Z-stack was converted into a maximum intensity Z-projection and then clipped to the 0.1 and 99^th^ percentiles and rescaled to the range [-1, 1] (Figure S4B).

#### Training strategy

During training, at every epoch, 8 random patches of size 128 pixels × 128 pixels (∼98 µm × 98 µm) were selected from each processed FOV. MONAI’s SmartCache Dataset was used to optimize data loading. All replicates in the *DiffAE model training FOV dataset* (processed as described above) were preloaded into cache memory at the start of training. During each epoch, FOV images were retrieved from this cache and the randomly sampled patches were cropped before being transferred to GPU memory for training. A batch of 12 processed FOVs, and correspondingly 96 patches, were used in an epoch (Figure S4E, F). PyTorch’s Distributed Data Parallel (DDP) was used to train both the semantic encoder and the DDIM U-Net jointly for 4602 epochs using four A100 80GB GPUs. A single epoch in this case corresponds to a pass through a quarter of the *DiffAE model training FOV dataset* due to the way the distributed sampler split the FOVs across the various ranks (GPUs). A custom data sharding schedule ensured that each rank saw a non-overlapping portion of the dataset. Therefore, with shuffling, near complete sampling of the FOV dataset would theoretically be achieved after ∼40 epochs. A one-cycle learning rate was used^117^ with the learning rate increasing to a maximum of 1e-4 during a warmup period of 200 epochs, then decaying for the remaining duration of training.

#### Model validation

To evaluate the learned representations and generative capabilities of the trained DiffAE, we performed qualitative assessments on representative images from 0, 6, and 21 dyn/cm^2^ timelapse datasets not used during DiffAE model training (see *Acquisition of 20X 48-hour 3D timelapse movies*; Figure S2C). We performed two controls to confirm that the reconstructed images are actively driven by the encoded latent vector. First, the brightfield conditioning image was scrambled before it was passed through the semantic encoder to obtain representations for denoising (Figure S2C). Second, the latent vector output from the semantic encoder was directly scrambled before the denoising process (Figure S2C). The output of each control was compared to both the standard inference output obtained from the unmodified brightfield conditioning image and the ground-truth VE-cadherin image (Figure S2C).

#### Latent-dimension sweep and model selection

To identify the optimal diffusion-autoencoder configuration, we performed a hyperparameter sweep over the latent dimension, training models with numbers of latent dimensions ranging from 8 to 1024 in powers of two. Each model was evaluated on representative images from timelapse replicates not used during DiffAE model training (see *Acquisition of 20X 48-hour 3D timelapse movies*) by reconstructing VE-cadherin crops from corresponding brightfield latent vectors (Figure S4A). Image reconstruction was assessed both visually and by computing the Pearson correlation coefficient between generated and ground-truth crops (Figure S4). As a positive control and estimate of the upper bound on reconstruction performance, we trained additional DiffAE models conditioned directly on VE-cadherin image patches with latent dimensions of 512 and 1024, using mEGFP-VE-cadherin fluorescence images as both the semantic encoder input and the diffusion decoder target. This control has a one-to-one correspondence between the input and target image, providing an estimate of the maximum achievable reconstruction accuracy. To account for stochasticity inherent to the diffusion sampling process, all evaluations were repeated across ten independent random seeds; we report the mean ± standard deviation (Figure S4B).

### Patch-based measured features

To quantify migration coherence and migration speed, we measured optical flow between consecutive timepoint images. We calculated optical flow vectors pixelwise on the log-normalized standard deviation bright-field projection of all FOV timepoints using the TV-L1 algorithm,^118^ as implemented in scikit-image following Sánchez Pérez *et al*. (2013).^119^ We selected TV-L1 over other classical methods (e.g., Horn-Schunck and Lucas-Kanade) because its total-variation regularization preserves motion discontinuities at cell boundaries while remaining robust to brightness-constancy violations in bright-field microscopy. We computed the optical flow fields at the FOV-level before cropping to the patch level (rather than directly computing flow at the patch level) to avoid edge artifacts arising from incomplete local information (Figure S5A).

We quantified “patch-based migration speed” as the mean magnitude of the unnormalized flow vectors within each patch. For each patch and time interval, we normalized the per-pixel flow vectors to unit directions and then averaged these unit vectors to obtain a mean unit vector for the entire patch. The magnitude of this mean unit vector gives us the *mean resultant length*, 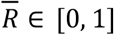, for the patch. Values close to 1 indicate high coherent migration in a common direction, whereas values close to 0 indicate directional cancellation consistent with low migration coherence (Figure S5B).

We found that frame-to-frame variation in 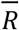 exceeds what would plausibly reflect changes in collective cell behavior, indicating sensitivity to subtle subcellular motion. Thus, we applied an exponential moving average (α = 0. 1) along the time axis to smooth the coherence feature. This moving average yields a smoothed per-patch feature capturing the degree of collective migration coherence, which we called “patch-based migration coherence”. The lower and upper limits of the migration coherence colormap were set to the 5th and 95th percentiles (0.3 and 0.9, respectively) of the migration coherence values across all analysis datasets (Figure 2-5, Figure S5).

“Number of nuclei in a patch” was quantified as the sum of nuclear segmentations within each patch and used as a measure of local cell density.

### Generating grid-based features

To generate a set of features agnostic to cell segmentations, patches of 128 pixels × 128 pixels were extracted from the full, preprocessed FOV (*Data preprocessing*) using a non-overlapping sliding window, yielding 36 samples per timepoint (“grid-based patches”). Each patch was then passed through the semantic encoder to obtain a 512-dimensional latent vector. All images from the *DiffAE model training FOV dataset* were processed in this way to generate the “DiffAE model training grid-based analysis dataset” (Table S2).

### Generating cell-centered features

To quantitatively validate ML-based features we additionally constructed cell-centered feature datasets. Patches for model inference were obtained in this context by taking the centroids of quality-controlled segmentations after trajectory filtering (*Single-cell trajectory filters*) and extracting image patches of 128 pixels × 128 pixels from the full, preprocessed FOV (*Data preprocessing*) centered on these segmentation centroids (“cell-centered patches”). Each patch was then passed through the semantic encoder to obtain a 512-dimensional latent vector.

In order to create an analysis dataset that integrates measured features with ML-based latent vectors, segmentation-based features (*Single-cell measured features* and *Single-cell dynamic features)* were computed for the segmentation used to define each cell-centered patch. The patch-based measured features (*Patch-based measured features*) of migration coherence and migration speed were also computed for each cell-centered patch.

All images from the *DiffAE model training FOV dataset* were processed in this way to generate the “DiffAE model training cell-centered analysis dataset” (Table S2).

### Principal component analysis of DiffAE features

We performed principal component analysis (PCA) on the *DiffAE model training grid-based analysis dataset* using the scikit-learn implementation to reduce dimensionality in the DiffAE latent feature space (Table S2). We then applied the PCA transformation that was fit on this reference dataset to all downstream grid-based and cell-centered feature datasets without recomputing PCs. The first 97 PCs capture 95% of the variance (Figure S3A). Individual PCs explain relatively small fractions of variance, with PC1 accounting for 3.61% and the top 10 PCs explaining 2.04% on average.

### Generative latent walks for interpretation of PC axes

In order to visually interpret the PCs, we performed generative “latent walks” along each of the feature axes. For each latent walk, we began by fixing the PCs except for the one being traversed to the mean value across the *DiffAE model training grid-based analysis dataset* (Table S2). We stepped through each PC from –3σ to +3σ in increments of σ, where σ denotes one standard deviation of the data along a given PC within the *DiffAE model training grid-based analysis dataset*. We then converted each of these points in PC space back to the original 512-dimensional latent space coordinate system using the inverse PCA transformation. Finally, we used these coordinates as conditioning for the Conditional DDIM U-Net to generate a representative VE-cadherin patch reconstruction from that point. We used a fixed noise sample for each reconstruction to enable comparison across all points in the latent walks. Repeating this process for each PC allowed visual interpretation of the learned latent space (Figure S3B).

### Coordinate transformation of PC-based features

To disentangle the aspects of PC1 and PC2 that encode orientation versus cell elongation (Figure S3B) within a patch we performed a polar coordinate transformation on the coordinate plane (PC1, PC2). We computed the polar angle as the four-quadrant inverse tangent of (PC1, PC2) using the numpy function atan2:

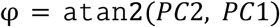

This coordinate is 2π-periodic and takes on a range of values between − π and π.

Initial investigation indicated that φ indeed captures the orientational aspect of PC1 and PC2, with φ = π ≡− π corresponding to parallel orientation relative to flow and φ = 0 corresponding to perpendicular orientation relative to flow. In order to get a feature variable that more intuitively mapped to the measured feature of cell orientation (see *Single cell measured features*), we shifted and rescaled this polar angle:

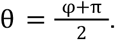

Like cell orientation relative to flow, the coordinate θ takes on values between 0 and π radians, with θ = 0 ≡ π corresponding to parallel orientation to flow and θ = π*/*2 corresponding to perpendicular orientation to flow (Figure S3C). We computed the polar radius *r* as 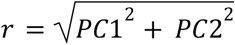 . This component captures cell elongation within a patch (Figure S3C). In order to get a feature variable that increases with increasing local density, we defined ρ =− *PC*3 (Figure S3B).

### Generative latent walks for interpretation of transformed feature variables

In order to visually interpret the transformed coordinates θ, *r*, and ρ, we again performed generative “latent walks” along each of these three feature axes. As we did with the PCs, we stepped through each axis while holding the other two features constant at the mean value across the *DiffAE model training grid-based analysis dataset* (Table S2). Because θ and *r* both have a priori defined ranges, we stepped along the full range present in the *DiffAE model training grid-based analysis dataset* at fixed increments instead of using the standard deviation. For consistency, we did the same with ρ, again using the full range of values present in *DiffAE model training grid-based analysis dataset*. Before we could map these coordinates back to the original 512-dimensional latent space coordinate system using the inverse PCA, we had to convert (θ, *r*) back to (PC1, PC2) and ρ back to PC3:

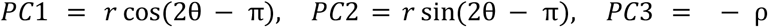

Once transformed back into latent coordinates, we used each point as conditioning for the conditional DDIM U-Net to visualize one sample from that point in latent space using a fixed noise sample (Figure 1D).

### Correlation between ML-based and measured features

In order to validate the qualitative interpretation of the generative latent walks, we computed Pearson correlation coefficients for all pairs of ML-based (see *Principal component analysis of DiffAE features* and *Coordinate transformation of PC-based features*) and measured features calculated on the same cell-centered patch in the *DiffAE model training cell-centered analysis dataset* (Figure 1E, Figure S3D, Table S2).

### Generating steady-state dynamics analysis datasets

We performed inference on the *Shear stress* and *VE-cadherin Exon3Del perturbation FOV datasets* as described in *Generating grid-based features*. We then transformed the resulting latent vectors to the 3-dimensional morphological state space of (θ, *r*, ρ) as described in *Principal component analysis of DiffAE features* and *Coordinate transformation of PC-based features*. Patch-based measurements of migration coherence and speed were calculated as described in *Patch-based measured features*. To restrict analysis to the steady-state response to shear stress, we filtered timepoints as described in *Visual steady-state filter* to exclude non-steady-state intervals, yielding the “Shear stress grid-based analysis dataset” and the “VE-cadherin Exon3Del perturbation grid-based analysis dataset” (Table S1, Table S3).

Similarly, we performed inference on the *Shear stress FOV dataset* as described in *Generating cell-centered features*. We transformed and filtered the resulting latent vectors using the same procedures, yielding the “Shear stress cell-centered analysis dataset” (Table S1).

### Dynamical systems characterization of cell state

#### Obtaining trajectories from grid-based patches

For the *Shear stress grid-based analysis dataset* and *VE-cadherin Exon3Del perturbation grid-based analysis dataset*, we defined trajectories in the morphological state space as the (θ, *r*, ρ)-values for a given grid-based patch with a fixed XY-location over time (Figure S4A). That is, given *m* grid-based patch locations in a replicate, we treated the values of **x** = (θ, *r*, ρ) from each grid-based patch over each of *n* timepoints in the timelapse as an independent observation of a vector-valued stationary process **x**(*t*) = (θ(*t*), *r*(*t*), ρ(*t*)) (Figure S4A):

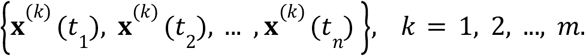

We base the assumption of stationarity on the fact that we are using only timepoints annotated as being at “steady state” as described in *Generating steady-state dynamics analysis datasets*.

#### Conceptual framework for cell state dynamics: stochastic differential equations

In line with the dynamical systems framework for cell states,^6–8^ we modeled the grid-based patch trajectories as realizations of an underlying Langevin-like stochastic differential equation (SDE):

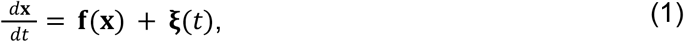

where *d***x***/dt* denotes the velocity of the state vector **x, f**(**x**) is a vector field that maps points in state space to deterministic velocities, and **ξ**(t) represents a noise process that is uncorrelated in time. The vector field **f**(**x**), referred to as the “theoretical vector field” in this work, is sometimes referred to as the drift coefficient (or vector of drift coefficients in each variable) of the resulting “drift-diffusion process.” Our goal was to then estimate this theoretical vector field in a data-driven manner using established properties of SDEs of this form.

In particular, established methods for obtaining an estimate 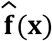 to the vector field 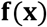 leverage the fact that random processes governed by equation (1) are Markovian.^120–123^ To estimate this data-driven vector field **f**(**x**) from trajectory data, these methods use the definition of the theoretical vector field **f**(**x**) as the first *Kramers-Moyal coefficient* of the vector-valued random process **x**(*t*):

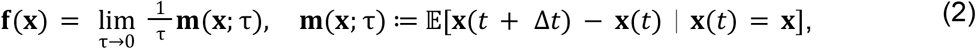

where the operator E denotes an average over all timepoints *t* and sampled trajectories (i.e., both a time and an ensemble average).

In order to determine whether these methods could be applied to the grid-based trajectories in morphological state space, we analyzed the autocorrelation functions (ACFs) of these trajectories to assess whether or not their temporal correlations were consistent with the assumption of Markovian dynamics. For a Markovian random process, the ACF is expected to be well approximated by a single exponential decay function as a result of the fluctuation-dissipation theorem.^124^ In contrast, deviations from single-exponential behavior or improved fits to multi-exponential functions indicate the presence of multiple characteristic correlation timescales.^124^

For a stationary scalar valued process *x*(*t*), the autocorrelation function (ACF) at lag time τ is defined as:

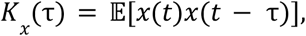

where the operator E denotes the average over timepoints *t*. In practice, this quantity is normalized to the range [− 1, 1] to get a scale-free measure of correlation (analogous to the Pearson correlation coefficient):

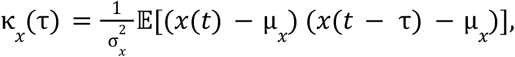

where μ_*x*_ and 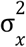 are the mean and variance of *x*(*t*), respectively. Note that by definition of stationarity, these are necessarily constant.

We computed the normalized ACF κ_*x*_ (τ) via the Fourier transform by noting that the autocorrelation is a convolution of *x*(*t*) with itself. Applying the convolution theorem for Fourier transforms to the unscaled ACF, we get:

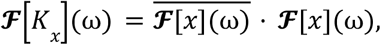

where 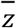 denotes the complex conjugate of *z* ∈ ℂ. Applying this theorem to the normalized ACF and using the linearity of the Fourier transform, we have:

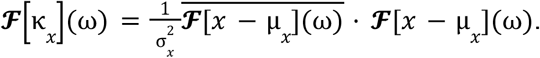

Let us denote 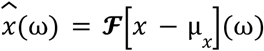. Then we have that:

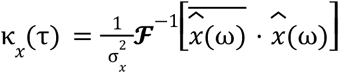

In Python, we performed this calculation using the numpy implementation of the Fast Fourier Transform (FFT) algorithm.

We computed κ^*x*^(τ) for each of *x* = θ, *r*, ρ in this manner for each individual grid-based patch trajectory (see *Obtaining stationary time series data from grid-based patches*) on a per-replicate basis in the *Shear stress* and *VE-cadherin Exon3Del perturbation grid-based analysis datasets*. We then computed the average and the (5^th^, 95^th^) percentiles across the given dataset at each lag τ, giving us a mean ACF curve and distribution for visualization (Figure S4B). We used this mean ACF curve to fit an exponential decay curve of the form *Ae*^−*B*τ^ + *C* (Figure S4B). We then computed the coefficient of determination (R^2^) as a measure of the goodness-of-fit of this exponential decay curve (Figure S4B).

#### Kernel-convolution based method for estimating the data-driven vector field from trajectory data

Based on the result that the ACF is well fit by a single exponential function across all replicates (Figure S4B), we proceeded with the approach defined in equation (2) to estimate a data-driven vector field 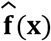 from grid-based trajectories in morphological state space. To deal with the limit of Δ*t* → 0 in equation (2), we used the assumption that the time Δ*t* between consecutive points along a trajectory was sufficiently small and took the approximation:

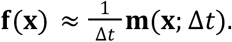

For the grid-based trajectories, Δ*t* is the time between consecutive frames in the timelapse data (1 frame = 5 minutes; Figure S4A). In order to obtain estimates that are more numerically stable, in addition to being more interpretable, we converted from units frames to units hours by taking:

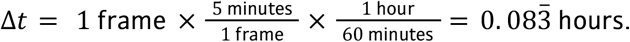

We computed the conditional average on the right-hand side of the above equation using a kernel-convolution based method developed by Gorjão and Meirinhos (2019)^121^ by adapting the accompanying implementation at https://github.com/LRydin/KramersMoyal (Figure S4C).

This method calculates a smooth numerical estimate of the vector field **f**(**x**) at each point in a discretized grid over (θ, *r*, ρ) by approximating a quantity called the *Nadaraya-Watson estimator*. Given random variables **X** and **Y** and a set of observations {(**X**_*n*_, **Y**_*n*_)} of those random variables, the Nadaraya-Watson estimator of the conditional expectation

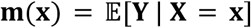

is defined as

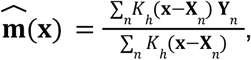

where *K*_*h*_ is a *kernel function* with *bandwidth h*. For the vector field **f**(**x**), the independent “predictor” variable is the trajectory position, **X**_*n*_ = **x**(*t*), and the dependent “response” variable is the displacement from this position, **Y**_*n*_ = Δ**X**_*n*_ ≔ **x**(*t* + Δ*t*) − **x**(*t*), where *n* indexes the unique observations (timepoint and trajectory index). Thus,

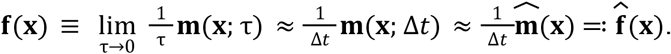

For computational efficiency, the method developed by Gorjão and Meirinhos (2019)^121^ uses properties of convolutions to compute the estimator 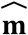 on a discrete grid instead of evaluating the kernel at each observation n and summing.

We began by discretizing the morphological state space into uniform bins of width 0. 05 in each of θ, *r*, and ρ centered at grid points {**x**_*k*_}. We then computed the following weighted and unweighted count histograms (Figure S4C):

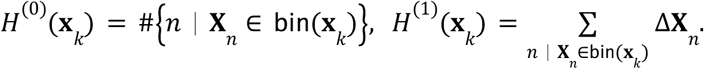

Consider a discrete convolution of the histogram *H*^(1)^ with a kernel *K*_*h*_ on the grid {**x** _*k*_} (Figure S4C), denoted by the operator ∗:

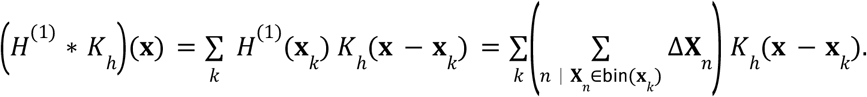

Because each point along a trajectory **X** _*n*_ can fall into only one bin *k*_*n*_, the above expression can be simplified to a sum over observations *n*:

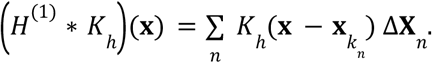

Furthermore, 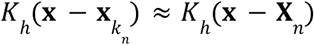 for 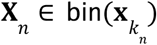 when the bin width is sufficiently small compared to the kernel bandwidth *h*. Under this assumption,

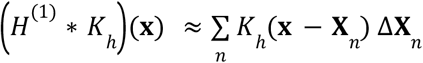

A similar identity holds for the denominator of the Nadaraya-Watson estimator:

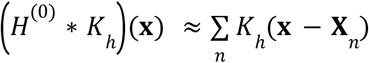

Thus, we can compute these discrete convolutions and take their quotient to get an approximation of the estimator 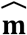:

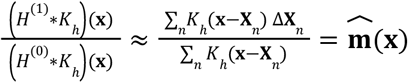

In order to compute the discrete convolutions, we evaluated the kernel function on a grid twice the size of the histogram grid to avoid numerical artifacts at the boundary (Figure S4C). We then used the function convolve from scipy.signal with the argument mode set to “same” to obtain the discrete convolution on the original grid over morphological state space (Figure S4C).

So that we could then specify different kernel functions in each of (θ, *r*, ρ), we used a *product kernel* formulation for the multivariate kernel *K*:

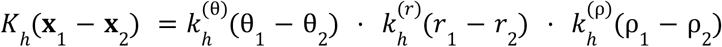

We used a Gaussian kernel for *r* and ρ:

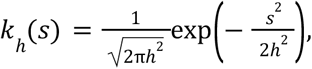

and a π-periodic exponential sine squared kernel for θ:

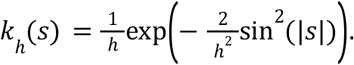

We chose a uniform bandwidth of *h* = 0. 15 for all variables, which is 3 times the bin width Δ*x* = 0. 05 of the discretization of the state space. This magnitude is sufficiently large to make the convolution approximation step mentioned above, providing sufficient smoothing of the vector field estimates for numerical stability without risking loss of accuracy caused by over-smoothing (Figure S4C).

For quantitative analysis, we computed the data-driven vector field in the full 3D space:

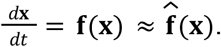

For qualitative visual analysis at 6 dyn/cm^2^ and 21 dyn/cm^2^, we computed the data-driven velocity vector fields in 2D in *r* and ρ:

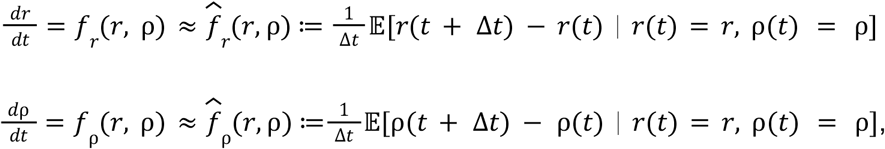

separately from the 1D velocity field in θ:

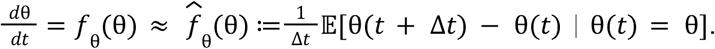

To account for the periodic nature of the variable θ, we “unwrapped” individual trajectories along θ using numpy’s unwrap function before computing the forward differences θ(*t* + Δ*t*) − θ(*t*). Doing so ensures that differences across the periodic boundary at 0 ≡ π are not misleadingly large. We performed each of these computations separately on a per-replicate basis.

#### Fixed point identification and classification

After obtaining the numerical values of the data-driven vector field on a grid over the specified state space (in 1, 2, or 3D, as outlined above), we identified and characterized the fixed points of this vector field to characterize the long-term behavior of the dynamics:

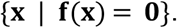

Because the method used to estimate **f**(**x**) only provides numerical estimates on a fixed grid, we could not apply a root finding algorithm directly to those outputs. Instead, we needed to get a *callable* estimate of **f** that could be evaluated at any point **x**, not just the chosen grid points. We did so via linear interpolation of the numerical estimates of **f**(**x**) between the values at the fixed grid points.

We then took the resulting callable function and used scipy’s optimize.fsolve method to find the fixed points as defined above. To ensure that the obtained roots were not biased by the selection of initial guess, we re-ran the root finding algorithm using 250 initial points sampled from the density of the data. We accumulated the output of this root-finding sweep by defining roots to be unique only up to 4 decimal places. In order to further keep the results robust against numerical fluctuations, we filtered out any found roots that fell outside of the 5^th^ to 95^th^ percentiles of the data along each axis.

Finally, we classified the local stability of these fixed points using a linear stability analysis. We took the callable vector field function and obtained the Jacobian matrix of partial derivatives using the Jacobian method from the Python package numdifftools. This method numerically estimates the partial derivatives using a central finite difference method, guaranteeing at least second order accuracy. At each fixed point, we evaluated this matrix function and numerically computed the eigenvalues, which determine the local linearized stability of the fixed point (i.e., whether it is an asymptotically stable, unstable, or saddle point). We then took the subset of these fixed points that were stable, denoted **x**^∗^, and used them for visualization and analysis.

#### Validation of low dimensional vector field visualizations by comparing fixed point locations

To validate the visualization and qualitative analysis of the dynamics in θ separate from those in (*r*, ρ), we compared the locations of the stable fixed points derived in the 1D (θ) and 2D (*r* and ρ) cases to those derived in the full 3D case (θ, *r*, and ρ together). Let 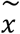 denote the location of the stable fixed point along *x* = θ, *r*, ρ as identified for the “low dimensional” (i.e., 1D or 2D) dynamics. That is, 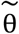 is the fixed point of the 1D system 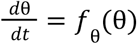, and 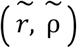 is the fixed point of the 2D system 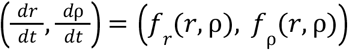. Likewise, let **x**^∗^ = (θ^∗^, *r*^∗^, ρ^∗^) denote the location of the stable fixed point as identified for the full 3D system 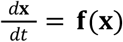. For each of the replicates analyzed in this manner (one each at 6 dyn/cm^2^ and 21 dyn/cm^2^), we computed the absolute error between these fixed point locations:

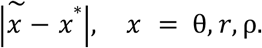

We found that the maximum error across both datasets and all features was ∼ 0.089, indicating that the location of the stable fixed point as identified by quantifying dynamics in θ separate from (*r*, ρ) is accurate to the stable fixed point as identified by quantifying the dynamics of all three features together.

#### Bootstrapping algorithm for fixed point location confidence intervals within a replicate

To get an estimate of uncertainty in the locations of the fixed points, we used bootstrap resampling (500 iterations with replacement) at the trajectory level. Let *m* denote the number of grid-based patch trajectories (i.e., unique patch locations) in a given dataset. For each bootstrap iteration, we sample *m* grid-based trajectory indices with replacement from the set {1, 2, …, *m*}:

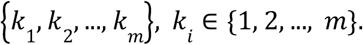

For each replicate, the number *m* of grid-based trajectories is equal to 36 times the number of FOV positions in the replicate after filtering (Table S1). We then pass the corresponding set of resampled trajectories into the data-driven vector field and fixed point estimation pipeline (see *Kernel-convolution based method for estimating the data-driven vector field from trajectory data* and *Fixed point identification and classification*). Based on the assumption of the stationarity of the data, resampling at the trajectory level is equivalent to resampling at the individual timepoint level.

For each bootstrap iteration we obtained a set of identified fixed points and their classified stability. We then developed a method to determine what is “the same” fixed point between each iteration, as we expect there to be some numerical fluctuation of the exact coordinates of a fixed point from resample to resample. For each “baseline” fixed point (as identified from the original unsampled set of trajectories), we took the identified stability (stable, unstable, saddle) and only considered fixed points from the bootstrap iterations with that same stability. Looping over the bootstrap iterations, if the current iteration had fixed points with the given stability, we computed the distance between each of these bootstrap fixed points and the current baseline fixed point. We then took the bootstrap fixed point with the smallest distance to the fixed point as that bootstrap iteration’s “match” to the baseline fixed point. We implemented this algorithm in a way such that a baseline fixed point can only have one match from a bootstrap iteration, and each bootstrap fixed point can only be matched to one baseline fixed point.

For a given baseline fixed point, we took the set of identified bootstrapped “matches” to be the distribution of the location of the fixed point in morphological state space. We used the (5th, 95th) percentiles of this distribution in each coordinate to define 90% confidence intervals for the locations of each fixed point. As an additional measure of confidence, we also computed for each fixed point the percentage of bootstrap iterations for which that point was assigned a match (“bootstrap detection rate”). We filtered out all fixed points with bootstrap detection rate < 40% for all downstream analysis.

#### Coloring morphological stable fixed points by behavioral measured features

To map migration coherence onto stable fixed points (θ^∗^, *r*^∗^, ρ^∗^) in morphological state space, we first associated each timepoint *t* along the grid-based trajectories with the migration coherence value computed for that patch between frames *t* and *t* + 1 (see *Patch-based measured features*). In this manner, we assigned each point **x**(*t*) = (θ(*t*), *r*(*t*), ρ(*t*)) in morphological state space along a grid-based trajectory with the migration coherence value measured over frames *t* to *t* + 1 (Figure S8D). Migration coherence associated with a stable fixed point (θ^∗^, *r*^∗^, ρ^∗^) was then computed as the mean value within a 0.25 × 0.25 × 0.25 bin centered on the point (θ^∗^, *r*^∗^, ρ^∗^) (Figure S8D). This procedure was repeated for all fixed points in the *Shear stress* and *VE-cadherin Exon3Del grid-based analysis datasets*. Fixed points were colored according to their associated migration coherence value. For consistency across summary plots, color scales were limited to the 5th and 95th percentiles of migration coherence values across all replicates in the *Shear stress* and *VE-cadherin Exon3Del grid-based analysis datasets*. The same procedure was used to map migration speed onto stable fixed points.

#### Validation of dynamics from grid-based patch trajectories

To compute the first-passage times of trajectories starting from the same binned location in morphological state space to reach a stable fixed point (θ^∗^, *r*^∗^, ρ^∗^) of the grid-based data-driven vector field 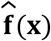 for a given replicate, grid-based and cell-centered trajectories from the respective *shear stress grid-based* and *cell-centered analysis datasets* were filtered to obtain only those trajectories with durations of at least 6 hours. These sets of trajectories were further restricted to just include trajectories that “reached” (θ^∗^, *r*^∗^, ρ^∗^), as these were the trajectories that by definition had a definite first passage time to the stable fixed point. For purposes of this analysis, a trajectory was considered to have reached the fixed point when it first came within a radius of 0.25 of the fixed point (Figure 2H). To investigate the potential bias the choice of radius has on the analysis, we conducted a parameter sweep. We found that this radius lay within a range of values in which the distribution of first-passage times to the fixed point stabilized while still maintaining a sufficient number of trajectories reaching the fixed point, indicating that the measurements were insensitive to small changes in the chosen radius (Figure S9C, D).

We discretized the morphological state space into bins of width 0.26 rad, 0.25, and 0.5 in θ, *r*, and ρ, respectively, and assigned trajectories to all bins they passed through before reaching the fixed point. These bin sizes were chosen to balance the need for robust statistics (i.e., sufficient number of trajectories for bin from which to compute a mean first passage time) versus a fine-grained resolution of state space. We treated these bins as sampling locations along continuous trajectories, with trajectory “start” points defined by entry into a given bin. Individual trajectories had one start point for each bin they visited. To ensure robust statistics, we excluded bins with fewer than 10 contributing trajectories from either the grid-based or cell-centered datasets.

We defined the mean first-passage time (MFPT) as the mean time over all trajectories in a bin from entry into that bin to reaching the fixed point. We computed the MFPT for a bin separately for grid-based and cell-centered trajectories, denoting them MFPT^*i*^_grid_ and MFPT^*i*^_cell_, where *i* indexes the bin in discretized feature space. We computed the Pearson correlation between MFPT^*i*^_grid_ and MFPT^*i*^_cell_ and performed a weighted linear regression using orthogonal distance regression (ODR), which accounts for the standard error of the mean within each bin (Figure S9A, B). We computed the Pearson correlation coefficients and ODR slopes for each timelapse in the *Shear stress grid based* and *cell-centered analysis datasets* (Figure S9E, F). To estimate the deviation of the binned MFPTs from the line ODR lines of best fit, we computed the shortest distance from each MFPT to this line of best fit.

#### Orthogonal projection of 3D vector field onto 2D plane for streamline visualization

To provide a more interpretable 2D view of the structure of the data-driven vector field for a selected example bistable replicate (12 dyn/cm^2^ replicate 5), we took the plane defined by the two stable fixed points (denoted 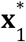 and 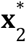) and the single identified saddle point that separates them (denoted **x**^*s*^) and orthogonally projected both the 3D coordinates **x** = (θ, *r*, ρ) as well as the data driven vector field 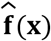 onto this plane. The first step of this process was identifying an orthonormal basis from which to define the in-plane 2D coordinates, which we obtained by performing the Gram-Schmidt orthogonalization process on the two in-plane vectors 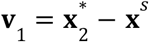 and 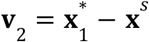:

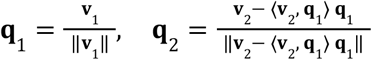

where ⟨·, ·⟩ denotes the vector dot product and ‖ · ‖ denotes the vector norm (magnitude). The change-of-basis matrix going from 3D (θ, *r*, ρ)-coordinates to 2D in-plane coordinates is defined as the 2 × 3 matrix *Q* with rows given by **q** and **q** . For any point **x** = (θ, *r*, ρ) in the plane defined by 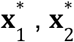, and **x**^*s*^, its 2D coordinates **u** = (*u*_1_, *u*_2_) with respect to the basis vectors **q**_1_ and **q**_2_ are obtained via the change-of-basis formula:

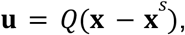

where **x**^*s*^ is subtracted so that the saddle point acts as a point of reference for the axes in the 2D plane. Because the basis **q**, **q** is orthonormal, the matrix *Q* satisfies the property that *Q*^*T*^*Q* is equal to the 3 × 3 identity matrix *I*_3_ and *QQ*^*T*^ is equal to the 2 × 2 identity matrix *I*_2_, where *Q*^*T*^ is the 3 × 2 matrix transpose of *Q*. Therefore, any point **u** in the 2D plane coordinate can be mapped back into (θ, *r*, ρ) coordinates via:

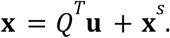

We then used the change-of-basis matrix *Q* to project the 3D data-driven vector field 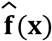 into the 2D plane, to obtain a visual representation of the in-plane dynamics. Note that, by properties of linear transformations and the matrix *Q*, it follows from the equation 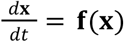 that

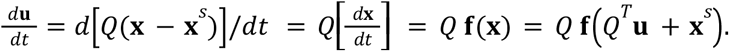

That is, if we define the 2D vector field **g**(**u**) as follows:

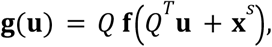

Then 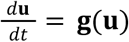. Therefore, the 2D-projected version of the data-driven vector field 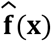 in the plane is given by

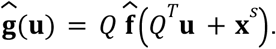

Note that fixed points of 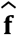 are also fixed points of 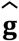:

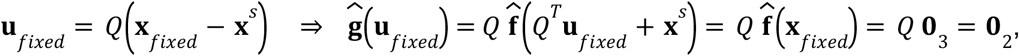

where **0**_*n*_ denotes the *n*-dimensional zero vector. Although this projection does not necessarily preserve local linear stability of fixed points, we found that it does for the chosen example replicate (Figure 4D).

To visualize the behavior of the projected dynamical system 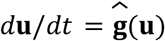, we defined a square grid over 2D plane-space with spacing 0.0125 a.u. between points, evaluated 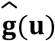 on this grid, and used matplotlib’s streamplot function to visualize the streamlines of the 2D vector field. The extent of the grid was determined as the range of values sufficient to visualize all projected fixed points with an additional margin. We visualized the basins of attraction of the projected stable fixed points 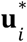 by labeling each point **u**_0_ in the 2D grid by which stable fixed point (*i* = 1 or 2) the solution **u**(*t*) obtained by integrating 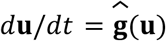 forward from **u**_0_ first “reaches.” A trajectory **u**(*t*) was classified as reaching stable fixed point if it came within within a radius of 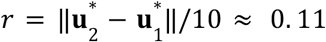 a.u. of that point. Solutions were computed numerically with a vectorized implementation of the fourth-order Runge-Kutta (RK4) method. The spacing of 0.0125 a.u. for the 2D mesh grid was chosen so that when points were colored by the labeled basin of attraction, there was sufficiently high resolution at the boundary between the basins for clear visualization.

For additional visualization, we computed the unstable and stable manifolds of the projected saddle point **u**^*s*^ = **0**. For the unstable manifold, we computed the Jacobian of 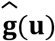 at **u**^*s*^ = **0** and took the eigenvector **v**^+^ of this matrix corresponding to the eigenvalue with positive real part. We then numerically computed the unstable manifold by integrating the ODE 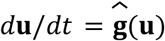 forward from initial points **u**(0) =± (0. 001)**v**_+_ using scipy’s solve_ivp method. Similarly, to compute the stable manifold we took the eigenvector **v** corresponding to the eigenvalue with negative real part and integrated the ODE 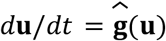 backward from initial points **u**(0) =± (0. 001)**v**_ using scipy’s solve_ivp method.

## Resource availability

### Lead contact

Further information and requests for resources and reagents should be directed to and will be fulfilled by the lead contact, Matheus Viana.

### Materials availability

Using the wild-type WTC-11 hiPS cell line background, we previously generated the Allen Cell Collection of hiPS cell lines in which each gene-edited cell line harbors a fluorescent protein endogenously tagged to a protein representing a distinct cellular structure of the cell. The cell line AICS-0126 cl. 41, which expresses mEGFP-tagged VE-cadherin are described at https://www.allencell.org and are available through Coriell at https://www.coriell.org/1/AllenCellCollection. For all non-profit institutions, detailed MTAs for each cell line are listed on the Coriell website. Please contact Coriell regarding for-profit use of the cell lines, as some commercial restrictions may apply.

### Data and code availability

We release all timelapse data used in this study in the OME-Zarr format to democratize their access. All the timelapse, training, and analysis datasets are publicly available on S3 at https://open.quiltdata.com/b/allencell/tree/aics/endo_cell_state_dynamics/. The data are also available via the AWS S3 API directly at s3://allencell/aics/endo_cell_state_dynamics. To facilitate data exploration, all datasets are accessible with BioFileFinder (BFF) at this link. BFF is an open-use web application that enables rich metadata search, filtering, and sorting of datasets that can be downloaded or viewed directly.^125^ Individual timelapses can be viewed using https://volumeviewer.allencell.org/ through BFF by selecting *Open with* > *Vol-E*. Users can additionally interactively explore the image and feature data simultaneously using Timelapse Feature Explorer (TFE). Image data overlayed with features from grid-based patches and cell segmentations are available in TFE: link for grid-based patches and link for cell segmentations.

Custom-written code was central to the analysis and conclusions of this paper. A subset of the code was generated with the assistance of Claude Sonnet 4.6. All necessary code to reproduce the results of this paper has been shared publicly on GitHub (https://github.com/AllenCell/endothelial-pipeline/). Any additional information required to reanalyze the data reported in this paper is available from the lead contact upon request.

