## Supplemental Figures for "Dynamics of ML-based Morphological Features Indicate a Shear Stress-Dependent Bifurcation of hiPSC-Derived Endothelial Cell States"

### Supplemental Figures & Tables

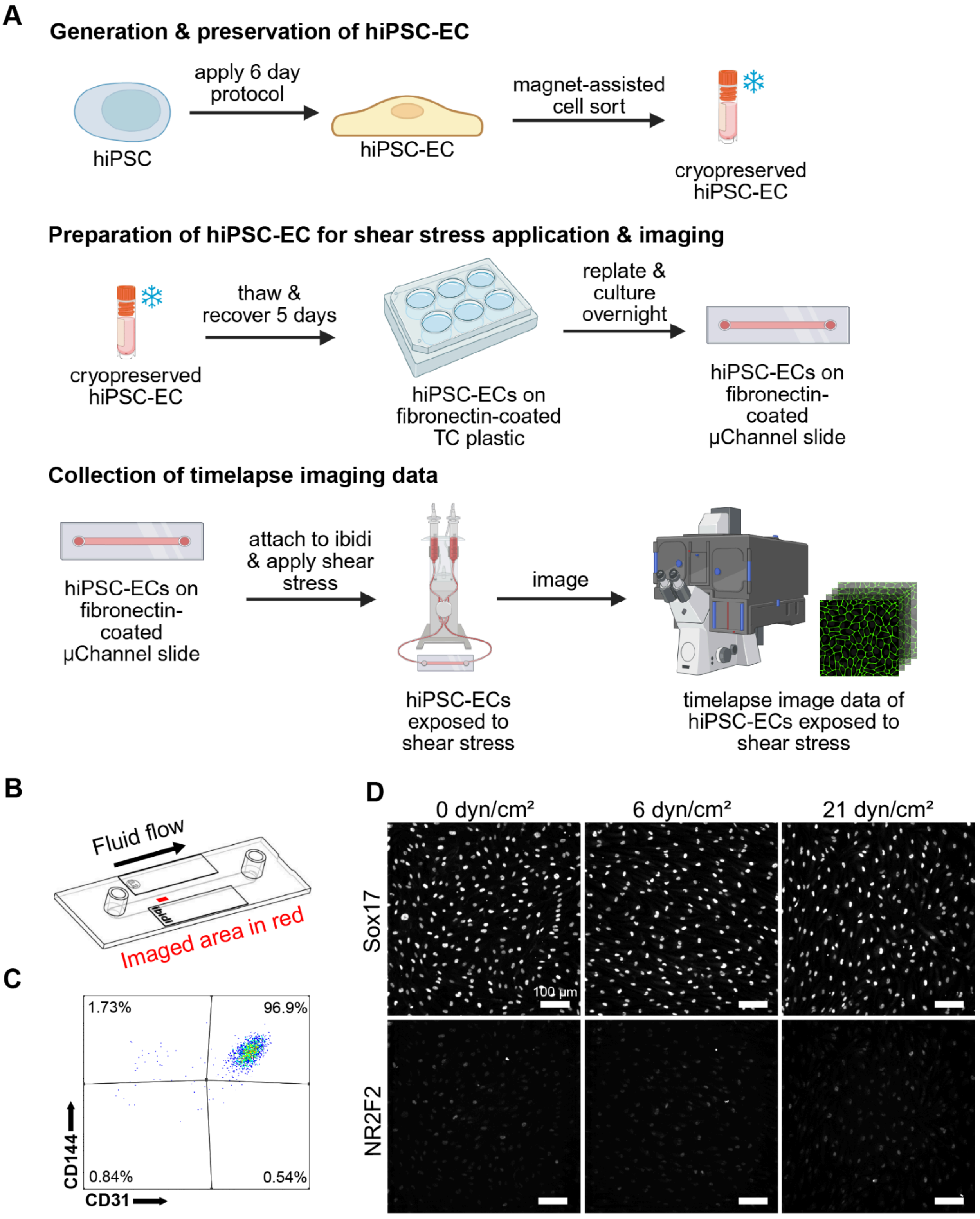


**Supplemental Figure 1. Generation, imaging, and characterization of hiPSC-ECs, showing cell purity, morphology, and marker expression under static and flow conditions. A.**

Experiment overview for generation of timelapse data. hiPSCs were differentiated into hiPSC-ECs using a 6 day protocol. hiPSC-EC populations were purified using magnet-assisted cell sorting for CD-144 or CD-31, and resulting sorted cells were cryopreserved. Cells were thawed into fibronectin-coated 6 well tissue culture (TC) plastic plates and recovered for 5 days before replating into fibronectin-coated glass-bottom ibidi µChannels. After cells were incubated in the µChannel overnight, shear stress was applied using the ibidi Pump System while timelapse imaging was performed. All procedures are detailed in Methods. **B.** Illustration of area of ibidi µChannel slide imaged during studies (red rectangle). **C.** Representative flow cytometry data of hiPSC-ECs stained with CD31 and CD144 indicate high purity of cell populations used for this study. **D.** hiPSC-ECs fixed and stained with SOX17 (arterial marker) and NR2F2 (COUP-TFII, venous marker) after 24 hours exposed to 0 dyn/cm$^2$, 6 dyn/cm$^2$ and 21 dyn/cm$^2$ of shear stress demonstrate that cells maintain a predominantly arterial phenotype after exposure to 6 dyn/cm$^2$ or 21 dyn/cm$^2$ shear stress (Methods). Maximum intensity Z-projections of Sox17 and NR2F2 immunofluorescence (Alexa Fluor 555 and 647) are contrast matched. Fluid flow direction is from left to right in all images. Scale bars, 100 µm. Schematic in **A.** generated using BioRender.

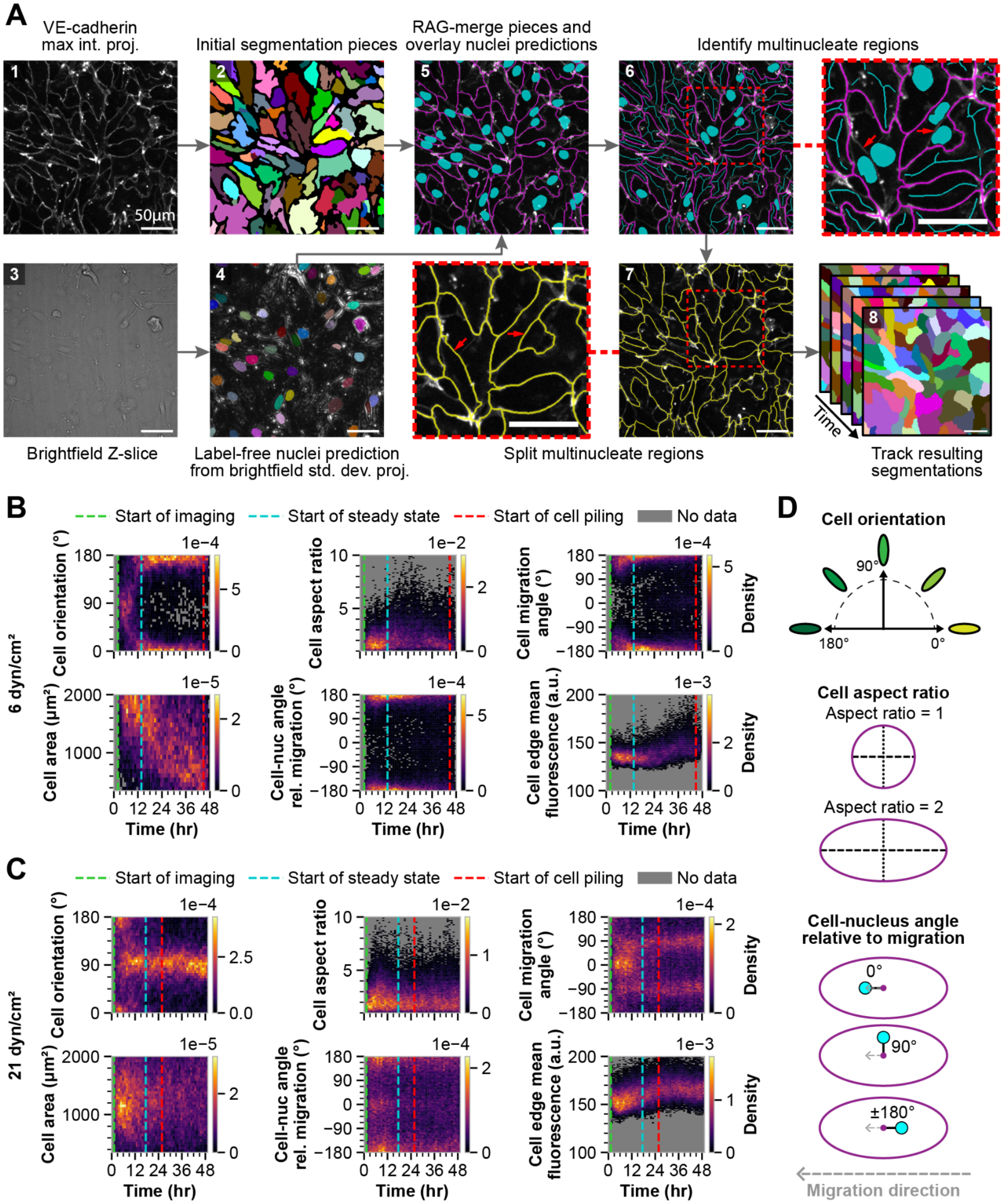


**Supplemental Figure 2. Generation of tracked 2D cell segmentations and measurements using VE-cadherin. A.** Panel 1) mEGFP-tagged VE-cadherin maximum intensity Z-projections of mEGFP-tagged VE-cadherin were thresholded to generate an initial binary mask. Panel 2) Initial segmentations were generated using a watershed algorithm using seeds derived from peak detection of a distance transform on the inverted initial binary mask. Panel 3) A representative Z-slice of the brightfield channel from panel 1 is shown. Panel 4) a standard deviation Z-projection of the brightfield channel is generated and used to create label-free nuclei predictions via a fine-tuned Cellpose "nuclei" model (Methods). Panel 5) A region adjacency graph (RAG) was constructed from the initial watershed segmentation, and adjacent regions were merged when VE-cadherin intensity along shared boundaries was below a threshold, producing the segmentation outlines shown in magenta. Label-free nuclei predictions are shown in cyan. Panel 6) Cell regions (magenta) containing a single nucleus are skeletonized (cyan lines) and used as seeds to preserve morphology, while multinucleate regions have their label-free nuclei predictions used as seeds (cyan filled regions). Inset highlights multinucleated cell segmentations where VE-cadherin signal was missed (red arrow). Panel 7) A second watershed segmentation was performed to split the multinucleate region along the missed VE-cadherin boundary. Inset highlights the corrected, final VE-cadherin–based segmentation. Panel 8) Single-cell trajectories were generated from tracking cell segmentations over time. To tolerate transient segmentation errors, cells were linked across short segmentation gaps by matching to the first subsequent segmentation with >50% pixel overlap within the next four timepoints (Methods). Scale bars, 50 um. **B.** Measurements of cell orientation, aspect ratio, migration angle, cell area, the cell-nucleus angle relative to the direction of migration, and edge mean fluorescence for a single imaging position from a representative replicate exposed to 6 dyn/cm$^2$ shear stress. Measurement definitions are described in Methods and illustrated in **D**. Time 0 denotes the onset of fluid flow. The green dashed line indicates the start of imaging. Density histograms show the distribution of each measurement over time during the 48-hour experiment, where yellow indicates higher density and purple indicates lower density. The blue dashed line indicates the start of steady state and the red line indicates the start of cell crowding (Methods). Grey regions indicate the absence of data. **C.** Same as **B** for a representative replicate at 21 dyn/cm$^2$. **D.** Schematics illustrating how cell orientation, aspect ratio, and cell-nucleus angle relative to migration direction were defined. Orientation is defined from 0° to 180°, where 90° is perpendicular to the direction of fluid flow, and 0° and 180° are parallel to the direction of fluid flow. As aspect ratio increases, the long axis (dashed line) elongates relative to the short axis (dotted line). The cell-nucleus angle relative to migration direction angle is defined as the angle between the line connecting the cell centroid (magenta point) and the nucleus centroid (cyan point) (black line) and the direction of migration (grey dashed line). See Methods for descriptions of all measured features.

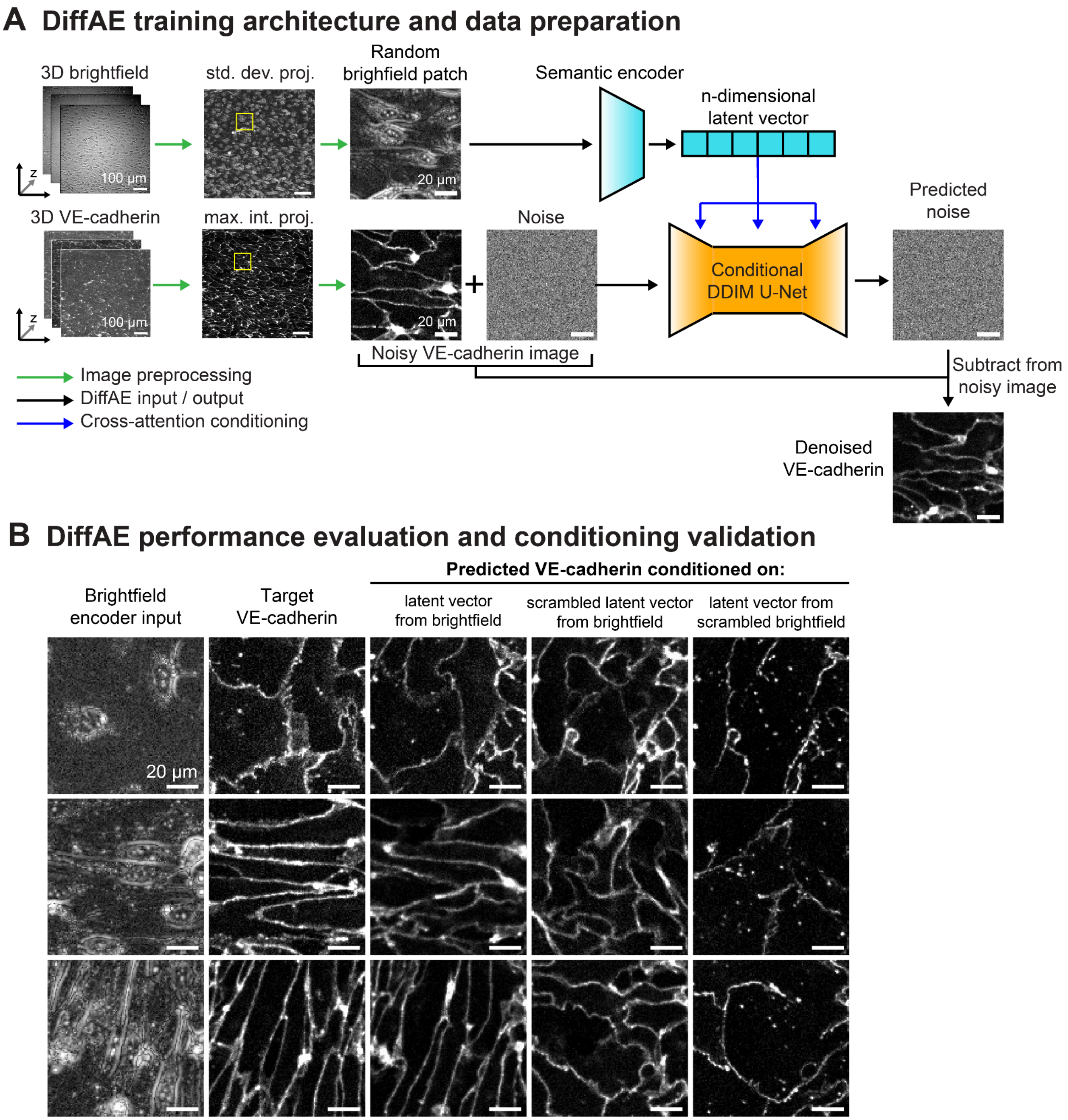


**Supplemental Figure 3. Diffusion autoencoder (DiffAE) training architecture and validation of semantic feature encoding. A.** 3D Z-stacks of brightfield and mEGFP-tagged VE-cadherin images were converted into 2D images using standard deviation and maximum intensity Z-projections, respectively. Paired 128 pixels × 128 pixels 2D brightfield and VE-cadherin-mEGFP patches were extracted at the same location, using an ensemble of randomly selected locations within the FOV, spanning all time points and shear stress conditions represented in the *DiffAE model training FOV dataset*. The semantic encoder compresses each input brightfield patch into a 512-dimensional latent vector. The diffusion model, a denoising diffusion implicit model (DDIM) based on a U-Net architecture, receives the VE-cadherin patch corrupted with added noise. The DDIM U-Net leverages cross-attention layers conditioned by the brightfield latent vector to predict the noise component. This predicted noise is subtracted from the noisy VE-cadherin patch, yielding a denoised prediction (Methods). Scale bars, 100 μm (FOVs) and 20 μm (patches). **B.** Representative patches from replicates not included in the *DiffAE model training FOV dataset* were used to validate the model's performance. The first two columns show the input brightfield patches the corresponding ground-truth VE-cadherin patches (target output). The third column shows the VE-cadherin pattern predicted by the DiffAE. The last two columns show two control experiments designed to test whether the prediction is driven by the semantic code: scrambling the latent vector or scrambling the input brightfield patch before semantic encoding. Scale bars, 20 μm.

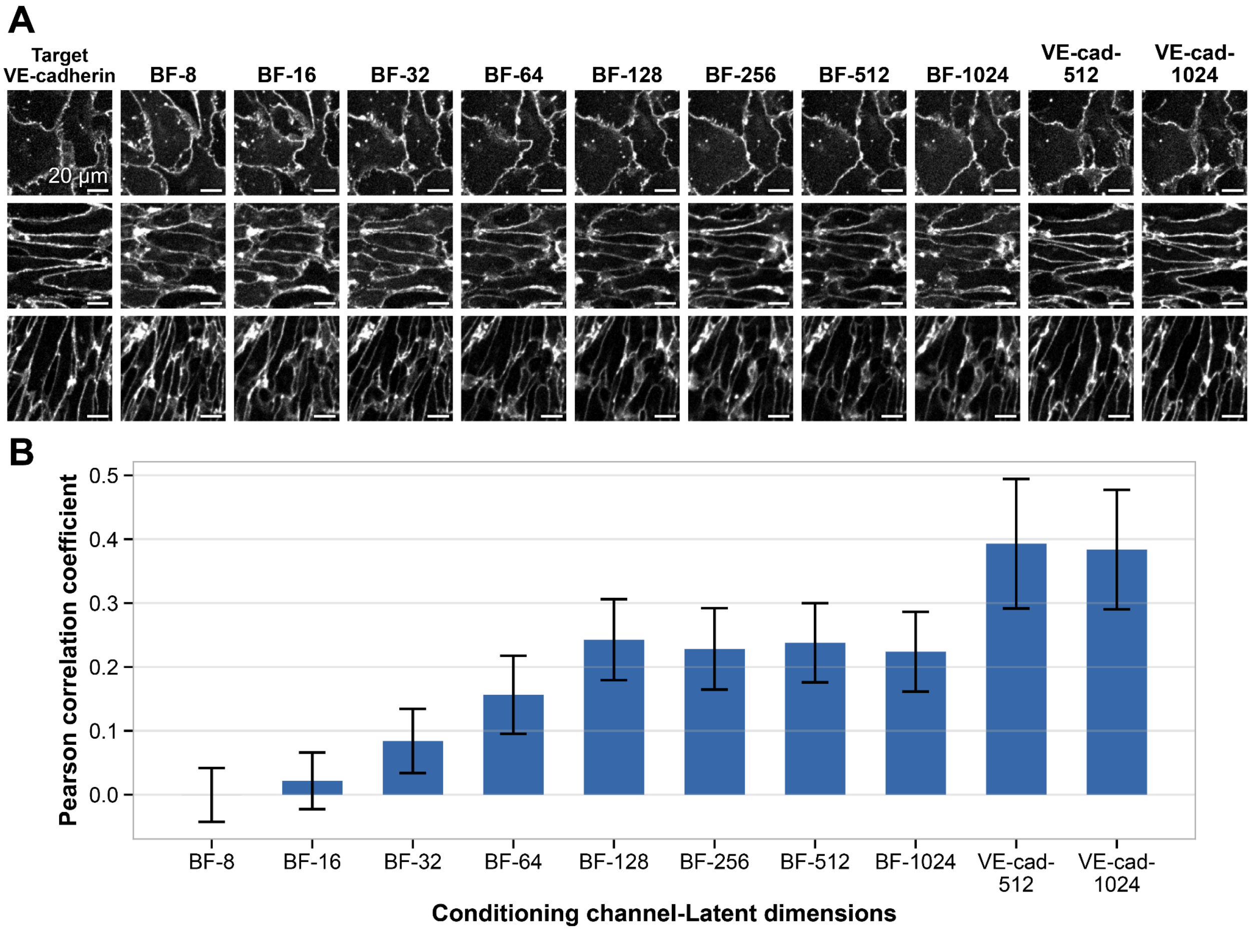


**Supplemental Figure 4. Comparison of VE-cadherin and brightfield-conditioned DiffAE models across latent dimension sweeps. A.** Diffusion autoencoders were trained across a sweep of latent dimensions (8–1024) for brightfield-conditioned models, and at selected latent dimensions (512 and 1024) for VE-cadherin-conditioned models. Representative patches from replicates not included in the *DiffAE model training FOV dataset* were used to assess model performance. The first column shows the ground-truth VE-cadherin target patch. Each row shows the corresponding prediction generated using the same diffusion noise input image across models. For brightfield-conditioned models, the semantic encoder input was the corresponding brightfield patch (shown in Figure S3). For VE-cadherin-conditioned models, the semantic encoder input was the target VE-cadherin patch itself. Scale bars, 20 µm. **B.** Reconstruction accuracy was quantified for the models shown in **A** as the pixel-wise Pearson correlation coefficient (r) between the predicted and ground-truth patches. Models were evaluated using a fixed panel of 10 patches excluded from training spanning distinct shear stress conditions and cell orientations, with each patch denoised using 10 independent noise seeds (n = 100 reconstructions per model). Bars represent the mean r, and error bars denote ±1 standard deviation across all reconstructions.

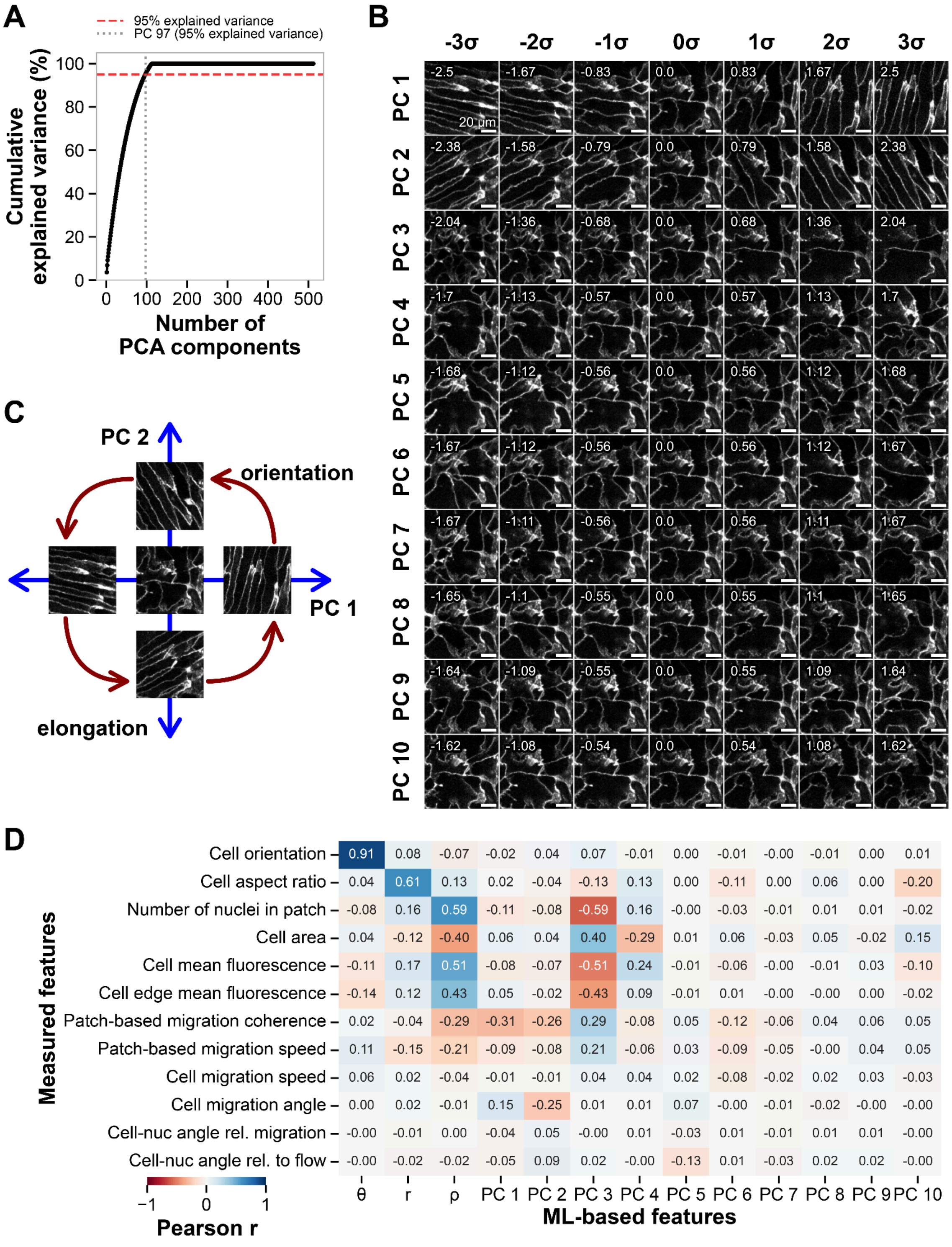


**Supplemental Figure 5. Derivation and validation of a low-dimensional, interpretable morphological feature space from DiffAE latent vector representations. A.** Cumulative explained variance for principal component analysis (PCA), shown as a percentage. The top 97 principal components (PCs) are required to explain 95% of the total variance (dashed red line), with each PC explaining less than 3.7% of the total variance. **B.** Generative latent walk performed along the top 10 PCs, spanning ±3 standard deviations (σ) of the PC value distribution observed in the *DiffAE model training dataset* for each component. **C.** Entanglement of orientation and elongation as respective angular and radial components in the PC1-PC2 plane suggest the need for a polar coordinate transformation. **D.** Pearson correlation coefficients (r) for paired ML-based and measured features from the *DiffAE model training cell-centered feature dataset*. Dark red and blue indicate strong positive and negative correlations, respectively, while white indicates no correlation. Out of the top 10 PCs, the top 3 show the strongest correlations (magnitude > 0.3) with measured features of interest. PCs 11-100 had correlation magnitudes < 0.21.

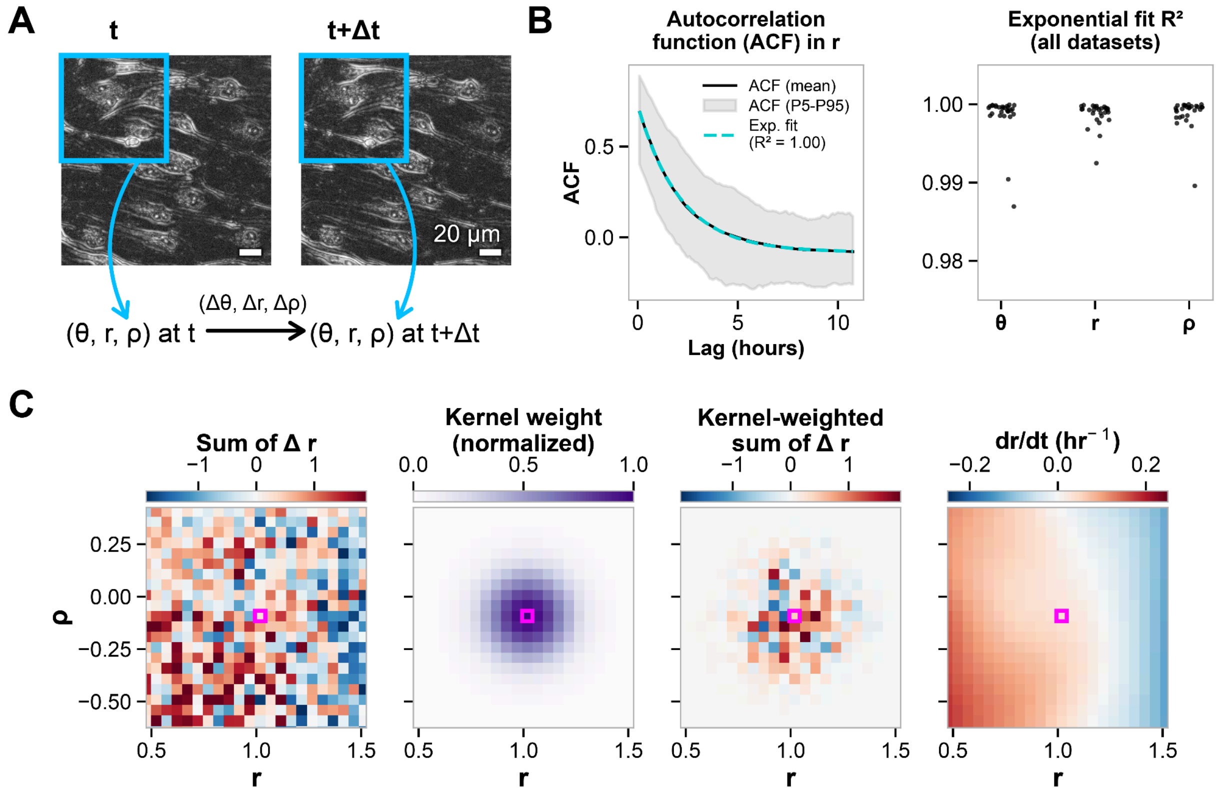


**Supplemental Figure 6. Estimation of a dynamical systems representation of cell state from grid-based patch trajectories. A.** Definition of trajectories and single-frame displacement vectors in ML-based feature space for grid-based patches. **B.** Characterization of trajectory fluctuations and correlation timescales. *Left:* Example autocorrelation function (ACF) for the variable $r$ as computed from grid-based trajectories. For each dataset and feature variable, we computed the ACF over a range of time lags for each individual trajectory and plotted the mean (solid black line) and 5$^{th}$ to 95$^{th}$ percentiles (gray shaded area). We then fit an exponential decay function to the mean ACF curve (dashed turquoise line) and computed the coefficient of determination ($R^2$) for the fit. *Right:* Summary plot of coefficient of determination ($R^2$) values for exponential decay fits to the mean ACF in θ, $r$, and ρ for each replicate in the *Shear stress grid-based analysis dataset* and *VE-cadherin Exon3Del perturbation grid-based analysis dataset* (each black point represents one replicate). **C.** Kernel-convolution-based method for estimating the data-driven vector field of morphological state space velocities as conditional expectation of forward differences in feature space. *Left:* For a given feature variable (shown: $r$), we computed single-timepoint displacements in that variable (shown: $\Delta r = r(t+1) - r(t)$). We then binned feature space (shown: $r$ and ρ) and computed the sum of these displacements in each bin. That is, each bin is colored by the sum of all $\Delta r = r(t+1) - r(t)$ such that $r(t)$ falls in that bin. *Center-left*: In order to take a locally weighted average to compute the data-driven vector field component (e.g., $dr/dt = \widehat{f}(r, \rho)$) for a given bin, we centered a multivariate kernel at that bin. *Center-right:* We then re-weighted the histogram based on this kernel. *Right:* Finally, we compute the data-driven vector field component for the given bin as the sum of the kernel-reweighted bin values. In practice, we do this process simultaneously for all bins in a computationally efficient manner using a discrete convolution of the original binned sum-of-displacements with the kernel function evaluated on the grid (Methods).

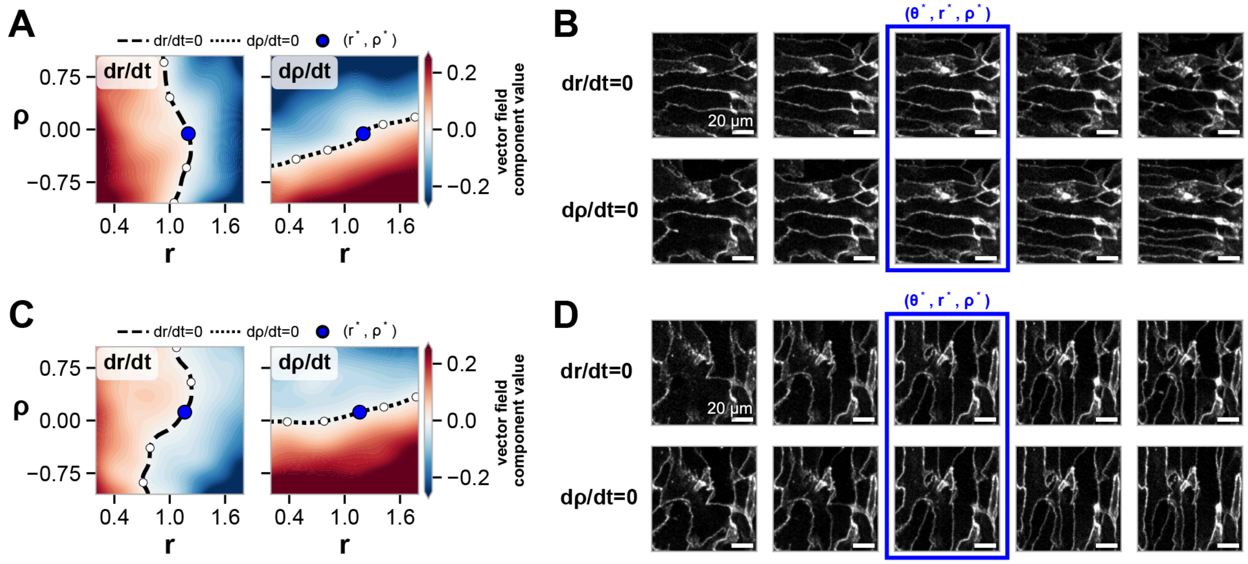


**Supplemental Figure 7. Latent walks along nullclines of data-driven vector fields. A** and **C.** Contour plots of the values of $dr/dt = \hat{f}_r(r, \rho)$ and $d\rho/dt = \hat{f}_\rho(r, \rho)$, respectively, as heatmaps for a representative replicate at 6 dyn/cm$^2$ (**A**) and 21 dyn/cm$^2$ (**C**) as shown in Figure 2B. The nullcline curves defined by $dr/dt = 0$ and $d\rho/dt = 0$ are shown as black dashed ($r$-nullcline) and black dotted (ρ-nullcline) lines, respectively, overlaid on the contour plots. The stable fixed point $\left(r^*, \rho^*\right)$ is shown as a blue point. **B** and **D.** Reconstructions along the $r$- (top row, increasing ρ left to right) and ρ- (bottom row, increasing $r$ left to right) nullclines at 6 dyn/cm$^2$ (**B**) and 21 dyn/cm$^2$ (**D**). Points in $(r,\ \rho)$-space used for these reconstructions are highlighted in white on the contour plots in panels **A** and **C**. The reconstructed patches highlighted inside the blue box are identical reconstructions at the stable fixed point $(r^*, \rho^*)$. All reconstructions are performed with $\theta = \theta^*$.

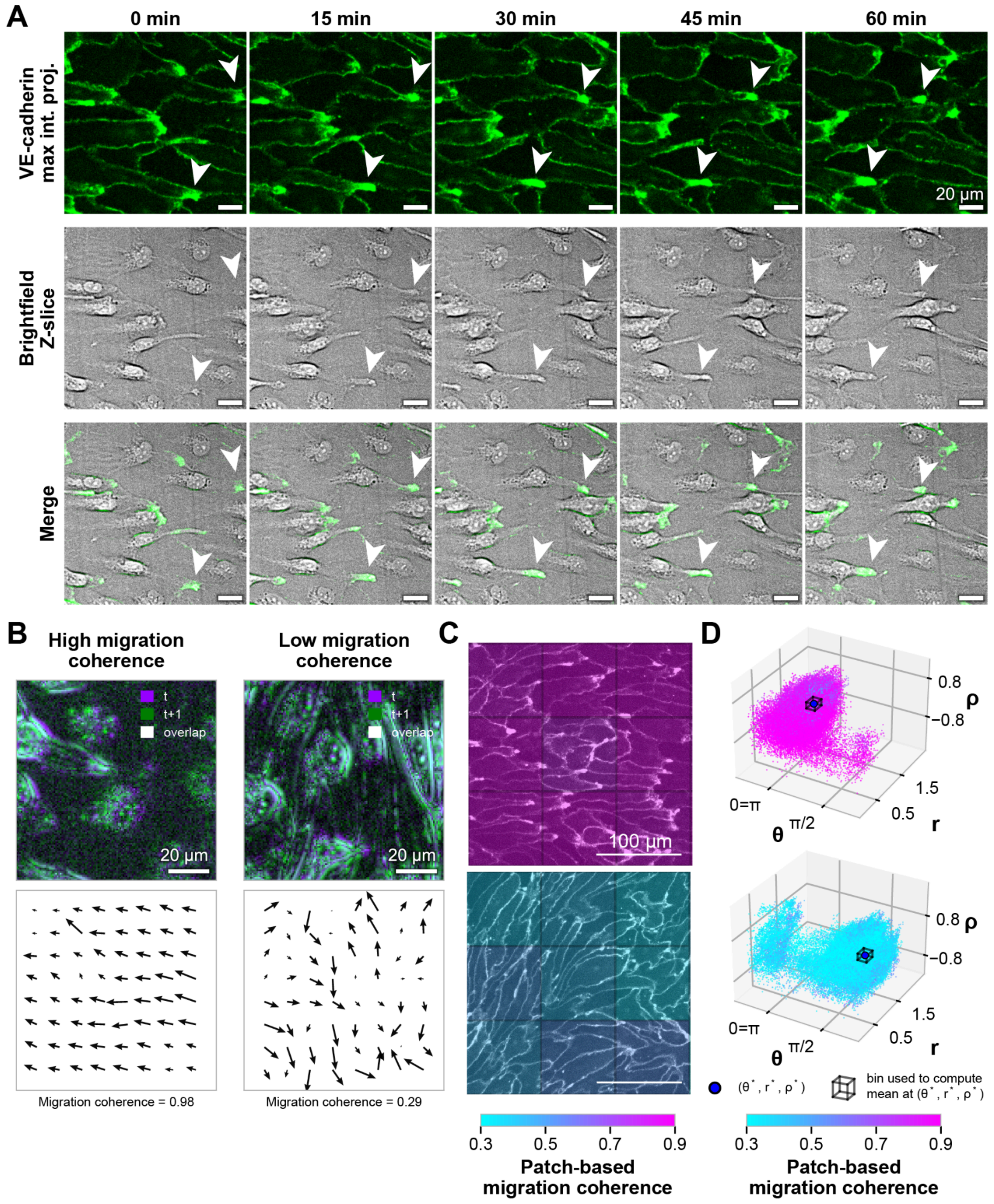


**Supplemental Figure 8. Quantifying migration coherence and mapping it onto stable fixed points in morphological state-space. A.** Representative images of mEGFP-tagged VE-cadherin maximum-intensity Z-projections, brightfield Z-slices, and merged images of cells exposed to 6 dyn/cm² shear stress over time (15-minute intervals shown). mEGFP-tagged

VE-cadherin is shown in green. Fluid flow direction is from left to right in all images. White arrows highlight two examples of VE-cadherin blobs that co-localize with retraction fibers as cells migrate against the direction of fluid flow (to the left). Scale bar, 20 µm. Associated timelapse of the mEGFP-tagged VE-cadherin maximum-intensity Z-projection, brightfield Z-slice, and merge is provided in Video S22. **B.** Representative examples of high migration coherence (left) and low migration coherence (right). Composite images of brightfield standard deviation Z-projections show paired patches at timepoint t (purple) and t+1 (green), where individual timepoints are separated by a 5-minute interval (1 frame). Pixels with no motion appear white. Scale bars, 20 µm. Migration coherence was quantified by computing optical flow between consecutive images, and normalized per-pixel flow directions were averaged within each patch. Values near 1 indicate high migration coherence in a common direction, whereas values near 0 indicate directional cancellation consistent with low migration coherence. Quiver plots show a subset of the un-normalized vectors for visualization (also used to calculate migration speed, see Methods). **C.** Representative 3 x 3 grids of mEGFP-tagged VE-cadherin maximum intensity Z-projection patches at 6 dyn/cm$^2$ (top) and 21 dyn/cm$^2$ (bottom) colored by the measured patch-based migration coherence for each patch using Timelapse Feature Explorer (Methods). Scale bars, 100 µm. **D.** Three-dimensional scatter plots of $(\theta, r, \rho)$ values for representative 6 dyn/cm$^2$ and 21 dyn/cm$^2$ replicates. Each point represents the $(\theta, r, \rho)$ -values for a single patch at a single timepoint t and is colored by the measured migration coherence calculated between timepoints t and t+1. Migration coherence was associated with each stable fixed point by averaging migration coherence values within a bin (black box) of size 0.25 in $\theta$, $r$, and $\rho$ around the fixed point $(\theta^*, r^*, \rho^*)$.

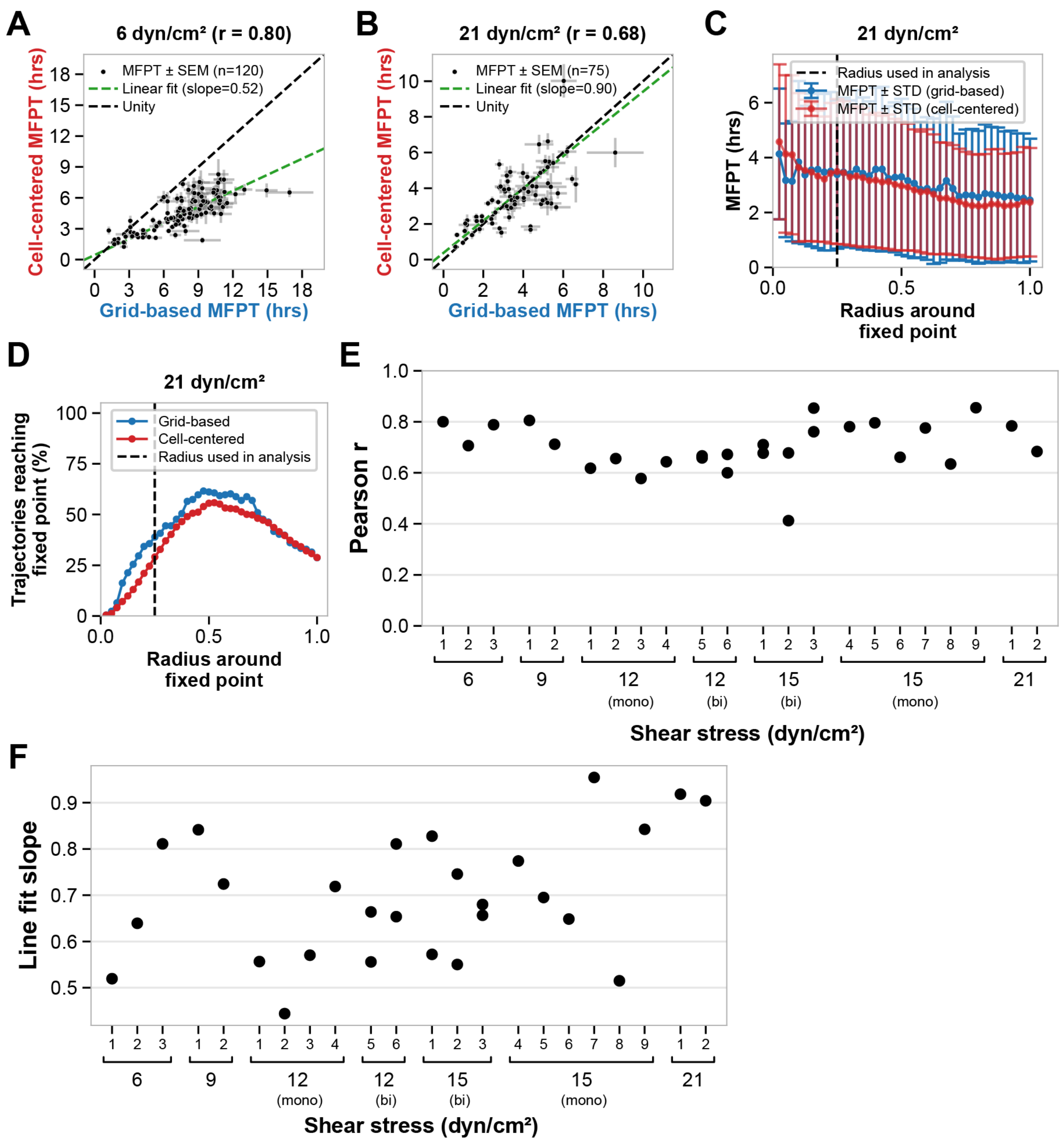


**Supplemental Figure 9. Grid-based dynamics capture behavior of features from cell-centered patches across timescales. A.** and **B.** The mean first passage time (MFPT) is computed as the average time trajectories take to reach the stable fixed point from starting locations binned over $(\theta, r, \rho)$-space (Methods). Shown are scatter plots of the MFPTs (black points) averaged for each starting bin, comparing grid-based and cell-centered trajectories originating from the same binned location in morphological state space and reaching the stable fixed point $\left(\theta^*, r^*, \rho^*\right)$ of the grid-based data-driven vector field $\hat{\mathbf{f}}(\mathbf{x})$ for representative 6 dyn/cm$^2$ (**A**) and 21 dyn/cm$^2$ (**B**) replicates. Grey error bars indicate the standard error of the mean

(SEM) for the grid-based and cell-centered MFPTs. A weighted linear fit using orthogonal distance regression is shown as a green dashed line, and the unity line (black dashed line) is shown for reference. N is the number of starting bins, and r is the Pearson correlation coefficient. **C.** Parameter sweep of the MFPTs for trajectories in the representative 21 dyn/cm$^2$ replicate (grid-based, blue; cell-centered, red) as a function of the radius used to define reaching the stable fixed point in $(\theta, r, \rho)$-space. Error bars indicate the standard deviation (STD) across trajectories. The selected radius (0.25) is indicated by the black dashed line within the plateau region of the MFPTs. **D.** Percentage of trajectories reaching the stable fixed point as a function of the threshold radius around the stable fixed point in $(\theta, r, \rho)$-space. The selected radius (0.25) is indicated by the black dashed line. The percentage initially increases with radius, then decreases above ~0.7 as more trajectories originate within the selected radius and are excluded from the analysis (Methods). **E.** Pearson correlation coefficients (r) and **F.** slopes of the linear fits for all replicates in the *Shear stress grid-based* and *cell-centered analysis datasets*, computed as described in **A** and **B**. Representative examples shown in **A** and **B** correspond to 6 dyn/cm$^2$ replicate 1 and 21 dyn/cm$^2$ replicate 1.

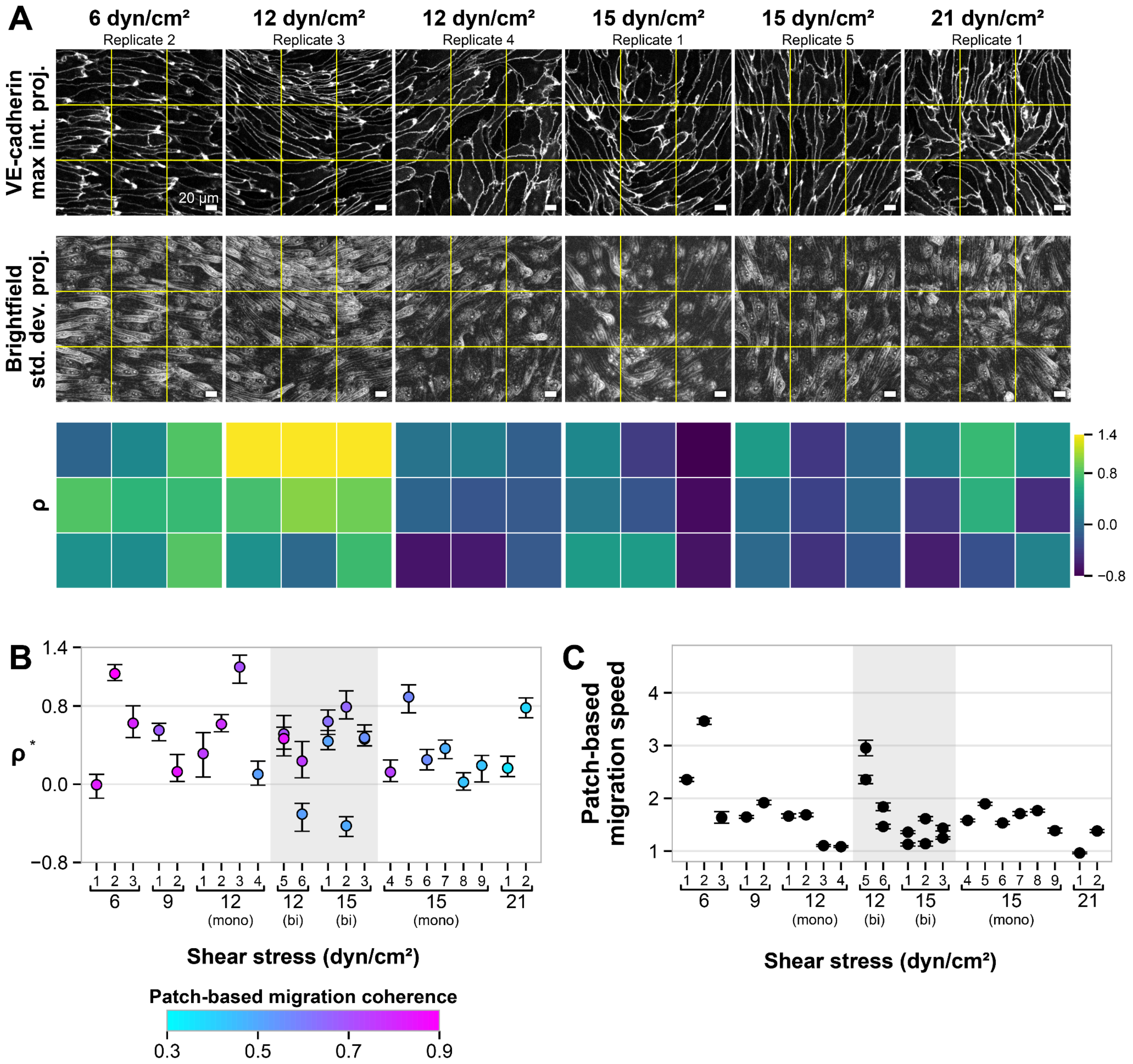


**Supplemental Figure 10. Changes in ρ across the transition from 6 dyn/cm$^2$ to 21 dyn/cm² in imaging data and stable fixed-point values. A.** Representative mEGFP-tagged VE-cadherin maximum intensity Z-projections and brightfield standard deviation Z-projections at steady state timepoints from 48-hour 3D timelapse imaging of hiPSC-ECs at 6 dyn/cm$^2$, 12 dyn/cm$^2$, 15 dyn/cm$^2$, and 21 dyn/cm$^2$ shear stress magnitudes are shown. The associated VE-cadherin maximum intensity Z-projections and brightfield standard deviation Z-projection timelapses stitched across all FOVs for the 6 dyn/cm$^2$, 12 dyn/cm$^2$, 15 dyn/cm$^2$, and 21 dyn/cm$^2$ replicates are provided in Videos S1, S4, S8, S10, S12, S14. The associated VE-cadherin maximum intensity Z-projection, brightfield Z-slice, and brightfield standard deviation Z-projection timelapses of the shown images for the 6 dyn/cm$^2$, 12 dyn/cm$^2$, 15 dyn/cm$^2$, and 21 dyn/cm$^2$ replicates are provided in Videos S7, S9, S11, S13, S15, S16. Fluid flow direction is from left to right in all images. Scale bars, 20 µm. Grid-based patch locations are shown in yellow, with corresponding ρ value for each patch colorized using viridis colormap shown below (for $\theta$, $r$, and patch-based migration coherence; see Figure 3B). **B.** Identified stable fixed point locations $\rho^*$ across replicates at 6 dyn/cm$^2$, 9 dyn/cm$^2$, 12 dyn/cm$^2$, 15 dyn/cm$^2$ and 21 dyn/cm$^2$, colored by the measured migration coherence at each stable fixed point for all replicates in the *Shear stress grid-based analysis dataset* (for $\theta^*$ and $r^*$, see Figure 4A). Error bars show bootstrapped 90% confidence intervals (Methods). Replicates exhibiting bistable fixed points at 12 dyn/cm$^2$ and 15 dyn/cm$^2$ are highlighted by the grey shaded region. **C.** Values of migration speed calculated at the stable fixed point for each replicate in the *Shear stress grid-based analysis dataset* (Methods). Error bars show bootstrapped 90% confidence intervals (Methods).

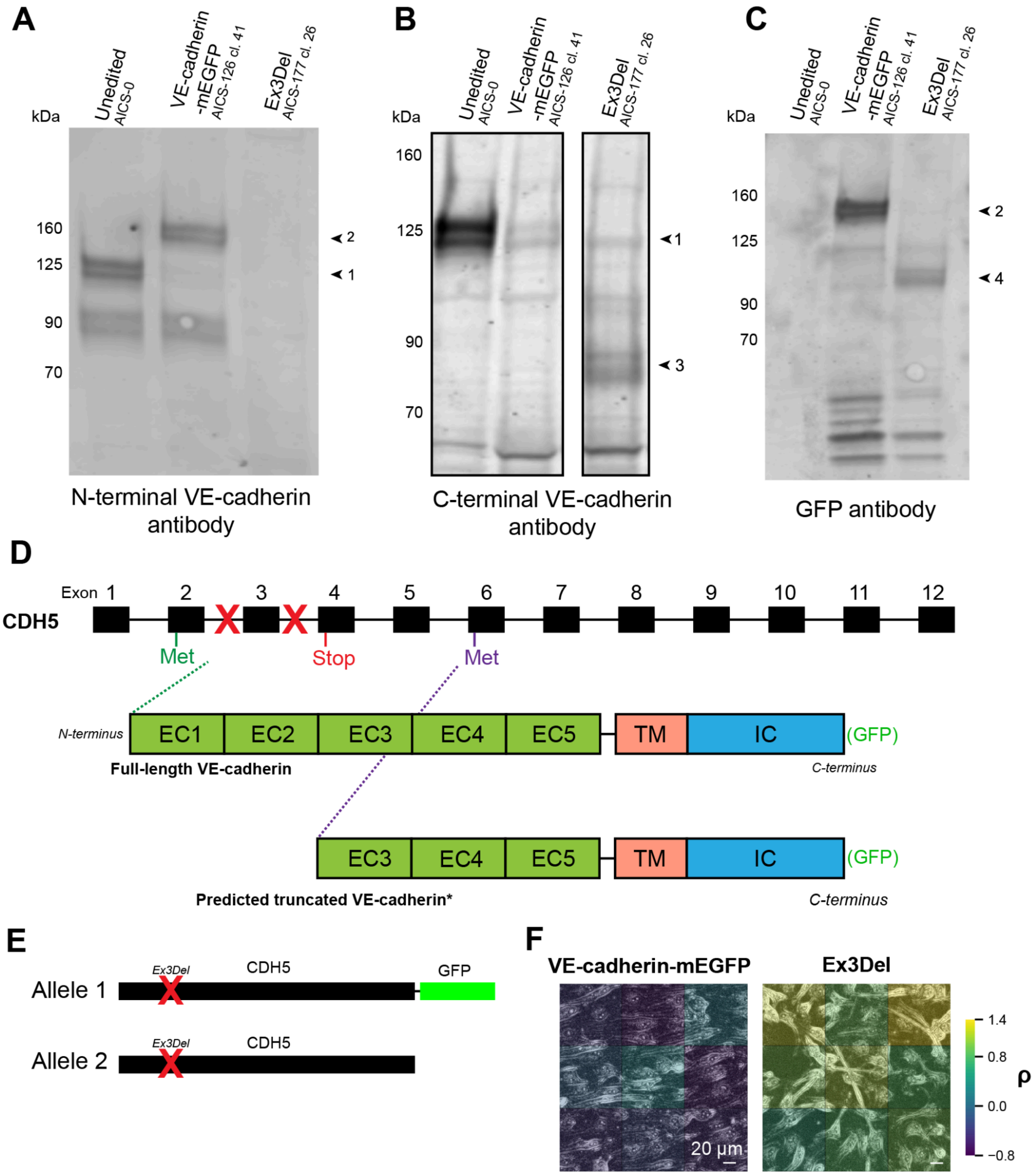


**Supplemental Figure 11. Western blots to characterize VE-cadherin exon 3 deletion hiPSC-ECs. A.** Western blot for VE-cadherin using an N-terminal antibody to stain hiPSC-ECs differentiated from unedited AICS-0, VE-cadherin-mEGFP, and Ex3Del cell lines. In unedited cells, a band is detected at ~125 kDa, consistent with full-length VE-cadherin (labeled as band 1). In the VE-cadherin-mEGFP cell line a band is observed at ~150 kDa, corresponding to mEGFP-tagged full-length VE-cadherin (labeled as band 2). No band is detected in the VE-cadherin exon 3 deletion cell line consistent with the N-terminal target of the antibody missing from this protein. **B.** Western blot for VE-cadherin using a C-terminal antibody in unedited, VE-cadherin-mEGFP, and Ex3Del cells. In addition to unspecific background bands observed in all samples, two additional bands are present. Once again, in unedited cells a ~125 kDa band is observed (band 1). Bands are not detected in the parental cell line, consistent with the GFP protein hindering binding of the antibody to the C-terminus of VE-cadherin. In the VE-cadherin exon 3 deletion cell line, a band is observed at ~75 kDa, consistent with an N-terminal truncated fragment of untagged VE-cadherin (band 3). **C.** Western blot using a GFP antibody in unedited, VE-cadherin-mEGFP, and Ex3Del cells. Bands are not detected in the unedited cells. In the VE-cadherin-mEGFP cell line, a band is detected at ~150 kDa, corresponding to full-length mEGFP-tagged VE-cadherin (band 2). In theEx3Del cells, a band is observed at ~110 kDa, consistent with a truncated version of mEGFP-tagged VE-cadherin (band 4). **D.** Schematic indicating the editing approach taken and hypothesized truncated form of VE-cadherin expressed in Ex3Del cells. The canonical start methionine residue of CDH5 is located in exon 2. In Ex3Del cells, exon 3 was removed, causing a frameshift and subsequent premature stop codon in exon 4. Due to the molecular weight of observed bands in **B** and **C**, we hypothesize that in Ex3Del cells a start methionine residue in exon 6 is utilized to express an N-terminal truncated version of VE-cadherin that is missing the extracellular cadherin (EC) domains 1 and 2. TM: transmembrane domain; IC: intracellular domain. **E.** Schematic of Ex3Del genotype. Exon 3 was deleted in both copies of CDH5. mEGFP was introduced into one allele of CDH5. **F.** Representative standard deviation Z-projection of brightfield images of VE-cadherin-mEGFP and Ex3Del cell lines colored by grid-based values of ρ using Timelapse Feature Explorer (Methods).

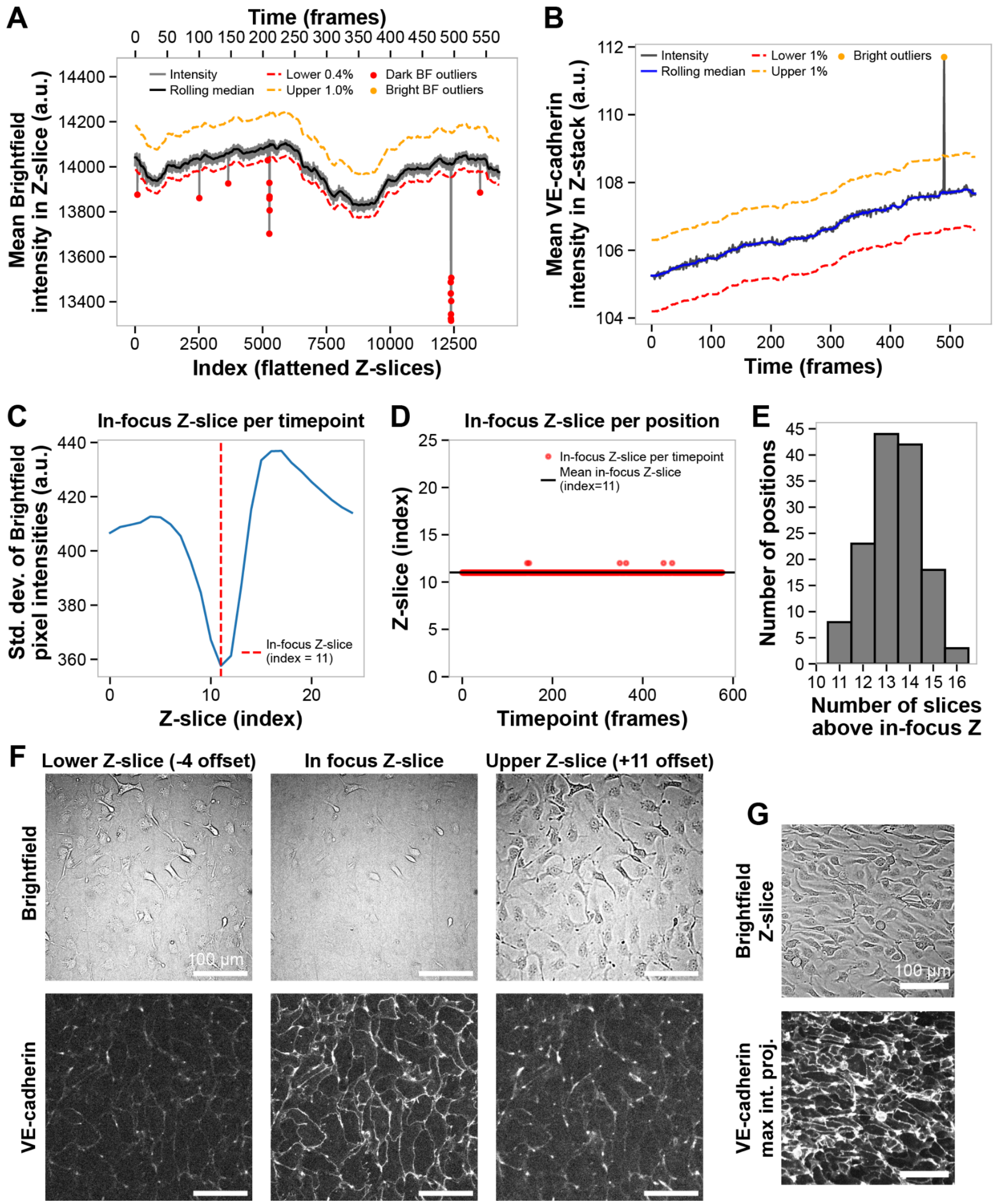


**Supplemental Figure 12. Filtering image data for downstream analysis. A.** Representative trace of the mean brightfield intensity in each Z-slice, plotted as a flattened index across time and Z-slice (grey). The rolling median of the average brightfield intensity, calculated over a window of 100 Z-slices, is shown in black. Timepoints were considered outliers if any Z-slice exceeded +1% from the rolling median (bright brightfield outlier, yellow dashed line) or fell below 0.04% of the median (dark brightfield outlier, red dashed line). **B.** Representative trace of the mean VE-cadherin-mEGFP pixel intensity across the Z-stack for each timepoint (grey). A rolling median over 12 timepoints was calculated for the mean intensity per Z-stack (blue). Timepoints deviating by more than ±1% from this rolling median were designated as bright (yellow dashed line) or dark (red dashed line) outliers. **C.** Standard deviation of brightfield pixel intensities at each Z-slice for a representative timepoint. The in-focus slice for that timepoint is defined as the Z-slice with the minimum standard deviation (dashed red line). **D.** The in-focus plane for a given position was defined as the mean of all in-focus planes across timepoints (shown in black). **E.** Histogram of calculated in-focus planes across all replicates. All replicates contained at least 11 Z-slices above the in-focus plane. **F.** Representative images show the calculated in-focus plane for a given timepoint, along with the upper and lower offsets used to generate consistent sub-Z-stacks near the glass. The in-focus plane consistently identifies a slice close to the glass rather than the cell centers. To balance the amount of out-of-focus blur above the cells, a lower offset of -4 slices was applied across all replicates. This approach ensures that the blur captured in the bright-field standard deviation z-projections reflects true differences in cell height and morphology, rather than technical variation in cell positioning within the Z-stack. Images are contrast-matched to the in-focus plane. **G.** Example images of cell crowding. VE-cadherin-mEGFP channel is shown as a maximum intensity Z-projection of all Z-slices; brightfield channel is shown as a single Z-slice. Scale bar, 100 μm.

### Table S1: Shear stress datasets

| Date | Cell line | Shear stress (dyn/cm²) | Replicate | N FOV positions | N Timepoints per FOV position | Position filter (N removed) | Single timepoint filter (N removed) | Steady-state filter (N removed) | Cell crowding filter (N removed) | Shear stress FOV dataset (N timepoints) | Shear stress grid-based analysis dataset (N patches) | Shear stress cell-centered analysis dataset (N patches) |
|---|---|---|---|---|---|---|---|---|---|---|---|---|
| 20250409 | AICS-126 cl. 41 | 6 | 1 | 6 | 544 | 0 | 98 | 918 | 572 | 1744 | 62784 | 168118 |
| 20250402 | AICS-126 cl. 41 | 6 | 2 | 6 | 571 | 1 | 48 | 268 | 1530 | 1041 | 37476 | 96620 |
| 20250618 | AICS-126 cl. 41 | 6 | 3 | 6 | 577 | 0 | 94 | 803 | 1926 | 717 | 25812 | 213077 |
| 20250428 | AICS-126 cl. 41 | 9 | 1 | 6 | 577 | 0 | 35 | 1051 | 975 | 1423 | 51228 | 208777 |
| 20250716 | AICS-126 cl. 41 | 9 | 2 | 6 | 577 | 0 | 70 | 685 | 1397 | 1354 | 48744 | 186707 |
| 20250604 | AICS-126 cl. 41 | 12 | 1 | 6 | 577 | 0 | 41 | 401 | 1842 | 1216 | 43776 | 176602 |
| 20260126 | AICS-126 cl. 41 | 12 | 1 | 6 | 577 | 1 | 45 | 495 | 1637 | 747 | 26892 | 190240 |
| 20260209 | AICS-126 cl. 41 | 12 | 2 | 6 | 577 | 0 | 36 | 702 | 2142 | 609 | 21924 | 130321 |
| 20260304 | AICS-126 cl. 41 | 12 | 3 | 6 | 564 | 0 | 13 | 918 | 984 | 1474 | 53064 | 282538 |
| 20260121 | AICS-126 cl. 41 | 12 | 4 | 6 | 577 | 0 | 9 | 994 | 121 | 2343 | 84348 | 326063 |
| 20250319 | AICS-126 cl. 41 | 12 | 5 | 6 | 543 | 2 | 9 | 396 | 957 | 951 | 34236 | 131546 |
| 20260216 | AICS-126 cl. 41 | 12 | 6 | 6 | 577 | 0 | 43 | 630 | 1758 | 1061 | 38196 | 201108 |
| 20250813 | AICS-126 cl. 41 | 15 | 1 | 6 | 577 | 0 | 75 | 688 | 476 | 2254 | 81144 | 394924 |
| 20260114 | AICS-126 cl. 41 | 15 | 2 | 6 | 577 | 0 | 30 | 570 | 1618 | 1271 | 45756 | 214786 |
| 20260211 | AICS-126 cl. 41 | 15 | 3 | 6 | 577 | 0 | 20 | 666 | 1866 | 921 | 33156 | 203153 |
| 20260225 | AICS-126 cl. 41 | 15 | 4 | 6 | 577 | 0 | 4 | 654 | 1446 | 1359 | 48924 | 255050 |
| 20250326 | AICS-126 cl. 41 | 15 | 5 | 6 | 577 | 0 | 18 | 522 | 667 | 2273 | 81828 | 306731 |
| 20260204 | AICS-126 cl. 41 | 15 | 6 | 6 | 577 | 0 | 39 | 633 | 1542 | 1275 | 45900 | 211861 |
| 20260202 | AICS-126 cl. 41 | 15 | 7 | 6 | 577 | 0 | 14 | 585 | 2154 | 722 | 25992 | 154069 |
| 20260128 | AICS-126 cl. 41 | 15 | 8 | 6 | 577 | 0 | 6 | 595 | 1782 | 1084 | 39024 | 193905 |
| 20260302 | AICS-126 cl. 41 | 15 | 9 | 6 | 577 | 0 | 13 | 918 | 1158 | 1376 | 49536 | 244920 |
| 20251001 | AICS-126 cl. 41 | 21 | 1 | 6 | 577 | 3 | 17 | 285 | 2 | 1427 | 51372 | 208353 |
| 20250611 | AICS-126 cl. 41 | 21 | 2 | 6 | 577 | 0 | 79 | 880 | 1777 | 786 | 28296 | 180907 |

### Table S2: DiffAE datasets

| Date | Cell line | Shear stress (dyn/cm²) | Replicate | N FOV positions | N Timepoints per FOV position | Position filter (N removed) | Single timepoint filter (N removed) | Cell crowding filter (N removed) | DiffAE FOV dataset (N timepoints) | DiffAE grid-based analysis dataset (N patches) | DiffAE cell-centered analysis dataset (N patches) |
|---|---|---|---|---|---|---|---|---|---|---|---|
| 20250618 | AICS-126 cl. 41 | 6 | 3 | 6 | 577 | 0 | 94 | 1926 | 1498 | 53928 | 213077 |
| 20250428 | AICS-126 cl. 41 | 9 | 1 | 6 | 577 | 0 | 35 | 975 | 2459 | 88524 | 208777 |
| 20250319 | AICS-126 cl. 41 | 12 | 5 | 6 | 543 | 2 | 9 | 957 | 1343 | 48348 | 131546 |
| 20250813 | AICS-126 cl. 41 | 15 | 1 | 6 | 577 | 0 | 75 | 476 | 2922 | 105192 | 394924 |
| 20250611 | AICS-126 cl. 41 | 21 | 2 | 6 | 577 | 0 | 79 | 1777 | 1663 | 59868 | 180907 |
| 20250818 | AICS-126 cl. 41 | 0 | 2 | 6 | 577 | 0 | 163 | 0 | 3299 | 118764 | 124257 |
| 20250714 | AICS-126 cl. 41 | 6, 24 | 1 | 6 | 577 | 0 | 89 | 140 | 3236 | 116496 | 336899 |
| 20250827 | AICS-126 cl. 41 | 21, 6 | 1 | 6 | 577 | 0 | 69 | 1122 | 2308 | 83088 | 278326 |

### Table S3: VE-cadherin Exon3Del perturbation datasets

| Date | Cell line | Shear stress (dyn/cm²) | Replicate | N FOV positions | N Timepoints per FOV position | Position filter (N removed) | Single timepoint filter (N removed) | Steady-state filter (N removed) | Delamination filter (N removed) | Cell crowding filter (N removed) | FOV dataset (N timepoints) | Grid-based analysis dataset (N patches) |
|---|---|---|---|---|---|---|---|---|---|---|---|---|
| 20260309 | AICS-126 cl. 41 CD31-sorted | 6 | 1 | 6 | 577 | 0 | 72 | 870 | 0 | 1890 | 683 | 24588 |
| 20250908 | AICS-177 cl. 26 | 6 | 1 | 6 | 577 | 0 | 98 | 828 | 0 | 0 | 2553 | 91908 |
| 20251029 | AICS-177 cl. 26 | 6 | 2 | 6 | 577 | 0 | 120 | 702 | 1659 | 0 | 1085 | 39060 |
| 20251119 | AICS-177 cl. 26 | 6 | 3 | 6 | 577 | 0 | 60 | 1036 | 1070 | 0 | 1280 | 46080 |
| 20260325 | AICS-177 cl. 26 | 6 | 4 | 6 | 577 | 0 | 75 | 582 | 1668 | 0 | 1191 | 42876 |

### Table S4: Nuclear label-free model training datasets

| Date | Cell line | Shear stress (dyn/cm$^2$) | Sample type | N FOV positions |
|---|---|---|---|---|
| 20240328 | AICS-126 cl. 41 | 0 | fixed | 5 |
| 20240328 | AICS-126 cl. 41 | 12 | fixed | 4 |
| 20250415 | AICS-126 cl. 41 | 15 | fixed | 50 |
| 20250415 | AICS-126 cl. 41 | 6, 21 | fixed | 50 |
| 20250415 | AICS-126 cl. 41 | 6 | fixed | 50 |

### Table S5: Segmentation summary

| Date | Cell line | Shear stress (dyn/cm$^2$) | Replicate | N FOV positions | N Timepoints per FOV position | N Nuclear segmentations | N Cell segmentations | N cell segmentations (after segmentation QC filtering) | Included in Shear stress datasets | Included in DiffAE datasets |
|---|---|---|---|---|---|---|---|---|---|---|
| 20250409 | AICS-126 cl. 41 | 6 | 1 | 6 | 544 | 949814 | 964072 | 213055 | X | |
| 20250402 | AICS-126 cl. 41 | 6 | 2 | 6 | 571 | 1576860 | 1309445 | 138305 | X | |
| 20250618 | AICS-126 cl. 41 | 6 | 3 | 6 | 577 | 1351990 | 1242228 | 437264 | X | X |
| 20250428 | AICS-126 cl. 41 | 9 | 1 | 6 | 577 | 1067172 | 1075882 | 278302 | X | X |
| 20250716 | AICS-126 cl. 41 | 9 | 2 | 6 | 577 | 1331755 | 1228908 | 289576 | X | |
| 20250604 | AICS-126 cl. 41 | 12 | 1 | 6 | 577 | 1283703 | 1226865 | 328864 | X | |
| 20260126 | AICS-126 cl. 41 | 12 | 1 | 6 | 577 | 1397673 | 1331459 | 488332 | X | |
| 20260209 | AICS-126 cl. 41 | 12 | 2 | 6 | 577 | 1334524 | 1236181 | 341144 | X | |
| 20260304 | AICS-126 cl. 41 | 12 | 3 | 6 | 564 | 1659968 | 1470560 | 454235 | X | |
| 20260121 | AICS-126 cl. 41 | 12 | 4 | 6 | 577 | 821207 | 955192 | 369706 | X | |
| 20250319 | AICS-126 cl. 41 | 12 | 5 | 6 | 543 | 1078697 | 1031648 | 188772 | X | X |
| 20260216 | AICS-126 cl. 41 | 12 | 6 | 6 | 577 | 1206034 | 1187816 | 389688 | X | |
| 20250813 | AICS-126 cl. 41 | 15 | 1 | 6 | 577 | 1143157 | 1205274 | 500952 | X | X |
| 20260114 | AICS-126 cl. 41 | 15 | 2 | 6 | 577 | 1268393 | 1242397 | 400788 | X | |
| 20260211 | AICS-126 cl. 41 | 15 | 3 | 6 | 577 | 1252582 | 1248522 | 440457 | X | |
| 20260225 | AICS-126 cl. 41 | 15 | 4 | 6 | 577 | 1029397 | 1120511 | 502829 | X | |
| 20250326 | AICS-126 cl. 41 | 15 | 5 | 6 | 577 | 1350019 | 1295331 | 376507 | X | |
| 20260204 | AICS-126 cl. 41 | 15 | 6 | 6 | 577 | 1221293 | 1229995 | 405769 | X | |
| 20260202 | AICS-126 cl. 41 | 15 | 7 | 6 | 577 | 1408965 | 1303962 | 382261 | X | |
| 20260128 | AICS-126 cl. 41 | 15 | 8 | 6 | 577 | 1197851 | 1186965 | 484879 | X | |
| 20260302 | AICS-126 cl. 41 | 15 | 9 | 6 | 577 | 880466 | 989945 | 427992 | X | |
| 20251001 | AICS-126 cl. 41 | 21 | 1 | 6 | 577 | 1028763 | 1115681 | 230680 | X | |
| 20250611 | AICS-126 cl. 41 | 21 | 2 | 6 | 577 | 1245752 | 1236845 | 364907 | X | X |
| 20250728 | AICS-126 cl. 41 | 0 | 1 | 6 | 577 | 999809 | 1139708 | 265291 | | |
| 20250818 | AICS-126 cl. 41 | 0 | 2 | 6 | 577 | 1260132 | 1358818 | 346989 | | X |
| 20250714 | AICS-126 cl. 41 | 6, 24 | 1 | 6 | 577 | 915066 | 1022164 | 400600 | | X |
| 20250827 | AICS-126 cl. 41 | 21, 6 | 1 | 6 | 577 | 1006866 | 1100116 | 459127 | | X |
